# The novel compensatory reciprocal interplay between neutrophils and monocytes drives cancer progression

**DOI:** 10.1101/2022.07.21.500690

**Authors:** Zhihong Chen, Nishant Soni, Gonzalo Pinero, Bruno Giotti, Devon J. Eddins, Katherine E. Lindblad, James L Ross, Nadejda Tsankova, David H. Gutmann, Sergio A. Lira, Amaia Lujambio, Eliver E.B. Ghosn, Alexander M. Tsankov, Dolores Hambardzumyan

## Abstract

Myeloid cells comprise the majority of immune cells in tumors, contributing to tumor growth and therapeutic resistance. Incomplete understanding of myeloid cells response to tumor driver mutation and therapeutic intervention impedes effective therapeutic design. Here, by leveraging CRISPR/Cas9-based genomic editing, we generated a mouse model that is deficient of all monocyte chemoattractant proteins (MCP). Using this strain, we effectively abolished monocyte infiltration in glioblastoma (GBM) and hepatocellular carcinoma (HCC) murine models, which were enriched for monocytes or neutrophils, respectively. Remarkably, eliminating monocyte chemoattraction invokes a significant compensatory neutrophil influx in GBM, but not in HCC. Single-cell RNA sequencing revealed that intratumoral neutrophils promoted proneural-to-mesenchymal transition in GBM, and supported tumor aggression by facilitating hypoxia response via TNF production. Importantly, genetic or pharmacological inhibiting neutrophil in HCC or qMCP-KO GBM extended the survival of tumor-bearing mice. Our findings emphasize the importance of targeting both monocytes and neutrophils simultaneously for cancer immunotherapy.

**In Brief:** Eliminating monocyte chemoattraction invokes compensatory neutrophil influx in tumor, and vice versa, rendering current myeloid-targeted therapies ineffective. Using genetic and pharmacological approaches combined with novel mouse models of GBM and HCC, we provide credence advocating for combinational therapies aiming at inhibiting both monocytes and neutrophils simultaneously.

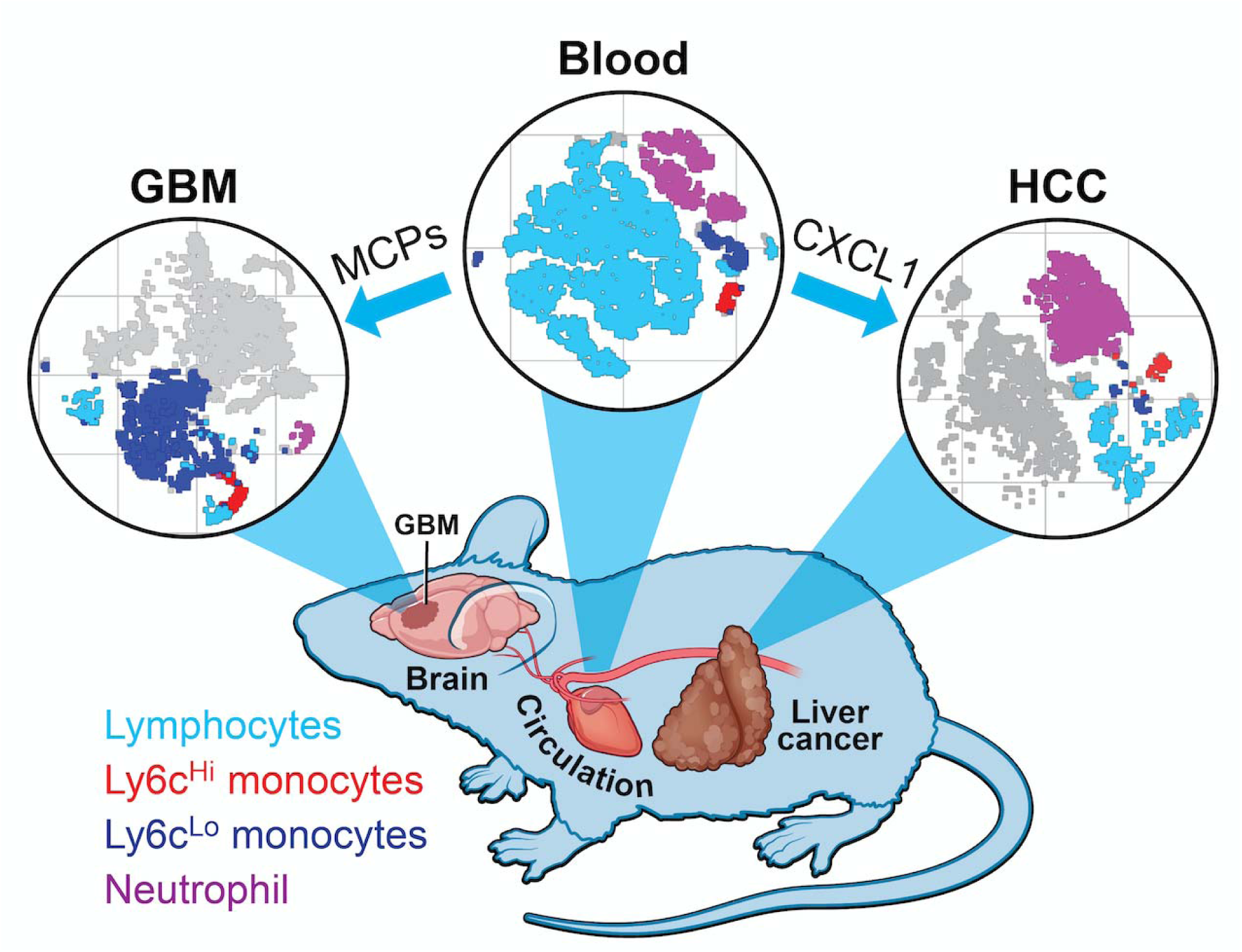

**Highlights:**

• Blocking monocyte chemoattraction results in increased neutrophil infiltration.
• Increased neutrophil recruitment induces GBM PN to MES transition.
• Inhibiting neutrophil infiltration in monocyte-deficient tumors improves mouse GBM survival.
• Blocking neutrophil, but not monocyte, infiltration in HCC prolongs mouse survival.

## INTRODUCTION

The strong interdependence between neoplastic and non-neoplastic cells in the tumor microenvironment (TME) is a major determinant underlying cancer growth. The glioblastoma (GBM) TME is composed of a wide variety of non-neoplastic stromal cells, including vascular endothelia, various infiltrating and resident immune cells, and other glial cell types (Becher et al., 2008; Becher et al., 2010; Jones et al., 2017). The predominant cell type in the GBM TME in both humans and murine tumor models are innate immune cells called tumor-associated myeloid cells (TAMs), which have been shown to promote tumor growth, invasion, and therapeutic resistance (Buonfiglioli and Hambardzumyan, 2021). In GBM, TAMs are composed of mixed populations, the most abundant of which are of hematopoietic origin, including monocytes and monocyte-derived macrophages (MOM). Less abundant, although still a significant presence, are brain intrinsic microglia (Mg) and hematopoietic-derived neutrophils (Chen et al., 2020). As such, it became appealing that treatment aiming at obliterating myeloid cells could offer promising outcomes for GBM patients. However, despite extensive efforts invested in both preclinical and clinical studies in the past decade, macrophage-targeted therapies in GBM have largely failed in clinical trials.

Chemokine gradients are essential for hematopoietic-derived myeloid cells to extravasate blood vessels and reach the tumor parenchyma. Monocyte chemoattractant proteins (MCPs) are a group of four structurally related chemokines that are indispensable for monocyte transmigration. In humans, they are encoded by *CCL2, CCL7, CCL8* and *CCL13* genes that are juxtaposed to each other on chromosome 17; while in mice, MCPs are encoded by *Ccl2, Ccl7, Ccl8* and *Ccl12* genes clustered on chromosome 11. All MCPs function through engaging the CCR2 receptor, but CCL7 may also interact with CCR1, and CCL13 with CCR3 (Proudfoot, 2002). CCL2 has been found to be critical in promoting recruitment of monocytes to the CNS (Fuentes et al., 1995). Neutralizing monoclonal antibodies against CCL2 had been developed and used in clinical trials against metastatic solid tumors but did not produce favorable outcomes. Meta-analysis of these clinical trials indicated that initial CCL2 inhibition may have unexpectedly caused subsequent increases in circulating CCL2 levels, possibly due to a compensatory feedback loop (Lim et al., 2016).

The Cancer Genome Atlas (TCGA) provides robust gene expression-based identification of GBM subtypes, including proneural (PN), mesenchymal (MES), and classical (CL) groups (Brennan et al., 2013; McLendon et al., 2008; Verhaak et al., 2010; Wang et al., 2017). These subtypes were established based upon the dominant transcriptional patterns at the time and location of tumor resection, and are not mutually exclusive of each other, i.e. multiple subtypes can co-exist within a single tumor, both at the regional (Sottoriva et al., 2013) and single-cell levels (Patel et al., 2014). Aimed at defining a unified model of cellular and genetic diversity, one study found that malignant cells in GBM exist in four major plastic cellular states that closely resemble distinct neural cell types, including: neural progenitor-like (NPC-like), oligodendrocyte progenitor-like (OPC-like), astrocyte-like (AC-like), and mesenchymal-like (MES-like) states (Neftel et al., 2019). Tumors with a MES-like state demonstrate striking similarities to the TCGA-MES subtype where both are enriched with TAMs (Hara et al., 2021) (Kaffes et al., 2019) (Wang et al., 2018). Using genetically-engineered mouse models (GEMMs) that closely resemble human PN, CL, and MES subtypes, we previously showed that driver mutations define myeloid cell composition in tumors (Chen et al., 2020). In contrast to *PDGFB*-overexpressing tumors (resembling human PN GBM) or *EGFRvIII*-expressing tumors (resembling human CL GBM) where the majority of myeloid cells are of monocytic linage, *Nf1*-silenced murine tumors (resembling human MES GBM) are enriched with neutrophils and brain-resident microglia (Chen et al., 2020) (Magod et al., 2021), similar to what was shown in human GBM (hGBM) (Gabrusiewicz et al., 2016).

In the current study, we specifically focused on blood-derived myeloid cells to determine the mechanisms of their invasion and the role they play in GBM progression. In addition, we wanted to determine whether there is a causal link between various myeloid cell infiltrates and GBM subtype dominance. By creating a combined all MCP-deficient mouse (qMCP) and generating *PDGFB*-driven gliomas we show that loss of expression of all MCPs in the TME resulted in a decrease of monocyte recruitment and extended survival of tumor-bearing mice. Surprisingly, abolishing all MCPs from the TME *and* tumor cells together resulted in compensatory neutrophil recruitment and a shift from PN-to-MES signature with no effects on survival of tumor-bearing mice. Single-cell RNA sequencing (scRNA-seq) and immunohistochemistry revealed that there is an increased presence of neutrophils in PDGFB-driven tumors when MCPs are abolished, which are predominantly localized in necrotic areas. Pharmacological targeting of neutrophils and their chemokine receptor CXCR2, or genetic ablation of the neutrophil recruiting chemokine *Cxcl1,* resulted in extended survival of PDGFB-driven tumor-bearing qMCP-deficient mice but not WT tumor-bearing mice. Considering GBM contains a mixture of cells with PN and MES gene signatures, these results suggest that effective therapies should target both neutrophils and monocytes. Since GBM is mainly monocyte-enriched, we next wanted to determine whether compensatory recruitment of neutrophils is a GBM-specific or CNS-specific phenomenon. To this end, we used a genetic mouse model of hepatocellular carcinoma (HCC). In contrast to PN and CL GBM, and even more so than MES GBM, the major immune infiltrates in a genetic HCC mouse model are neutrophils. We demonstrate that abolishing monocytes has no impact on survival of HCC-bearing mice but leads to an increase in recruitment of neutrophils. Decreased neutrophil recruitment resulted in extended survival of HCC tumor-bearing mice.

Collectively, our results suggest there is a compensatory interplay between monocyte and neutrophil recruitment in tumors. When we targeted each pro-tumorigenic population separately, we observed compensatory recruitment of the other. Therefore, novel therapeutic strategies should aim at simultaneously targeting both populations to overcome these compensatory recruitment mechanisms.

## RESULTS

### MCPs exhibit region-specific expression patterns, which inversely correlates with patient survival

We and others have previously demonstrated that decreased expression of *CCL2* correlates with extended survival of patients with GBM (human GBM; hGBM) (Chen et al., 2017). Similarly, mouse GBM (mGBM) models with decreased *Ccl2* expression exhibited prolonged survival relative to WT GBM-bearing mice (Chen et al., 2017). However, in these mice, no decrease in TAM recruitment was observed, suggesting other MCP family members likely compensate for *Ccl2* loss. To determine the basis for this compensatory adaptation, we sought to determine whether the expression of the other MCP members (*CCL7, CCL8, CCL13)* is elevated in hGBM and whether their increased expression serves as a predictor of patient prognosis. We stratified IDH-WT patients using TCGA datasets into high MCP expressers (+0.5 standard deviation (SD) from the Mean of all samples) or low MCP expressers (-0.5 SD from the Mean, Fig. 1A, insets) and compared their survival. Patients with elevated *CCL7* or *CCL8* expression had a shorter survival time compared to the low expressers (Fig. 1A, P<0.05). A similar trend was also observed with *CCL13*, although the survival difference was not statistically significant (Fig. 1A, P=0.18).

**Figure 1.**
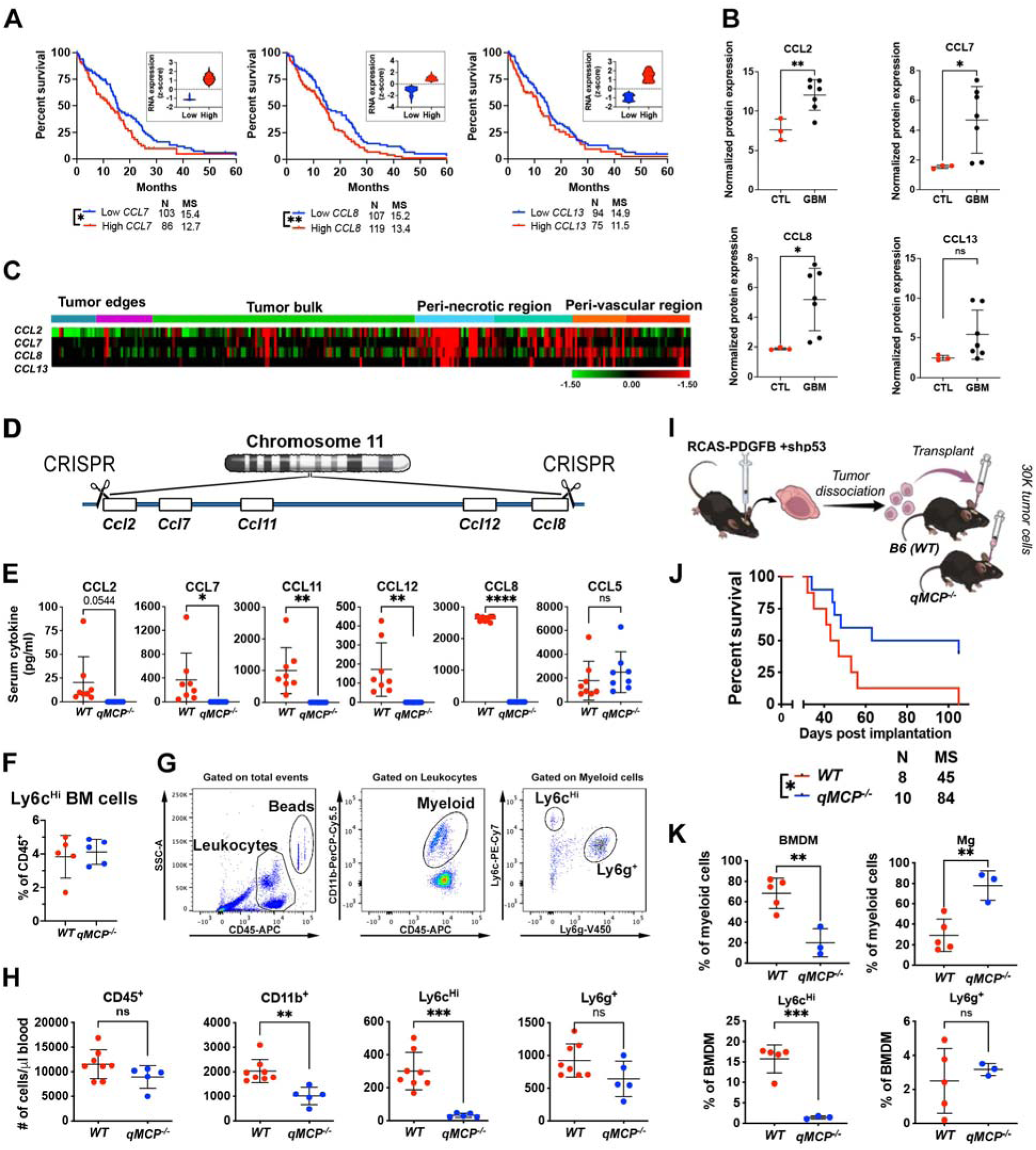
Generation and validation of *qMCP^-/-^* mouse. (**A**) Correlations between *CCL7, CCL8,* and *CCL13* expression levels and patient survival were analyzed using an IDH-WT cohort from TCGA. High and low expression were defined as +/- 0.5 STDEV from the mean of all samples (n = 260). Both Log-rank and Mantel Cox (MC) tests were applied. P = 0.1868 by MC for *CCL13*. (**B**) Normalized protein expressions of MCPs examined by Olink proteomic assay. Two-tailed Student’s *t*-test. **(C)** Expression distribution of *MCP* family members in human GBM tissue as determined in tandem by laser capture microdissection and RNA-seq queried from the IVY Gap database (n=34). **(D)** Schematic illustration of CRISPR/Cas9-mediated deletion of the *MCP* genes. **(E)** Serum MCP levels were measured by ELISA following LPS treatment. CCL5 was used as an internal control. Two-tailed Student’s *t*-test. **(F)** Flow cytometry quantification of Ly6c^Hi^ monocytes in the bone marrow of healthy adult mice. **(G)** Multiplex flow cytometry analysis was used to enumerate blood cells in the circulation. **(H)** Analysis of blood cells in healthy adult mice. Two-tailed Student’s *t*-test. **(I)** Schematic illustration of orthotopic transplantation of primary tumors. (**J**) Kaplan Meier-survival curves of *PDGFB*-driven tumors generated in *WT* and *qMCP^-/-^*mice. (**K**) flow cytometric quantification of myeloid cells in tumors at humane endpoint. Student’s *t*-test. *p<0.05, **p<0.01, ***p<0.001, ****p<0.0001, ns = not significant. MS = median survival.

To determine the protein concentration of all MCPs in hGBM samples, we used Olink multiplex proteomics to quantify a predefined group of immune-related proteins (Fig. S1). A total of 7 IDH-WT hGBM tissues along with three normal brain samples were analyzed (sample information in Table S1). When MCP expression was specifically assessed, all were increased in GBM samples, with CCL2, CCL7 and CCL8 exhibiting the highest levels (Fig. 1B). Using the IVYGap (IVY Glioblastoma Atlas Project) database (https://glioblastoma.alleninstitute.org/), we found that *MCPs* are predominantly transcribed in the peri-necrotic and peri-vascular regions, rather than in the tumor bulk or leading edge (Fig. 1C). This is consistent with the observation that the leading edge is mainly populated by microglia (Muller et al., 2017), which do not require MCPs for their infiltration.

### Decreased, but not abolished, *Ccl7, Ccl8, or Ccl12* expression leads to extended survival of GBM-bearing mice

To investigate biological significance of the reverse correlation between MCP expression and survival of GBM patients, we leveraged GEMMs deficient in the expression of individual MCPs. In these experiments, we orthotopically transplanted *PDGFB*-driven primary mGBM tumor cells into the brains of wild-type (*WT*) mice and mice deficient in *Ccl7* or *Ccl8/12* expression (Fig. S2A). It should be noted that *Ccl7* or *Ccl8/12* are depleted in the TME, but are retained in tumor cells. While we observed increased survival in these tumor-bearing mice relative to *WT* controls, there was no reduction in TAM content (Iba1^+^ cells; Fig. S2B). Based on this finding, we wondered whether complete genetic deletion of MCPs from both tumor cells *and* the TME could further extend survival. To address this question, we induced *de novo* tumors in *WT;Ntv-a, Ccl2^-/-^;Ntv-a*, *Ccl7^-/-^;Ntv-a,* and *Ccl8/12^-/-^;Ntv-a* mice by injecting a combination of RCAS-shp53 and RCAS-*PDGFB* in the frontal striatum. Unexpectedly, the survival benefits previously observed with the transplant model were abolished when these *MCPs* were deleted in both the tumor cells and the TME (Fig. S2C). When we examined the immune composition of the tumors by flow cytometry, using *Ccl8/12^-/-^;Ntv-a* mice to represent entire cohort, there was no difference observed in the proportion of infiltrating monocytes or microglia compared to *WT;Ntv-a* controls (Fig. S2D). These results suggest that partial loss of MCPs, which may not trigger a compensatory response, provides a survival advantage, but complete deletion of an individual MCP may cause other MCP members to compensate.

### Creating quintuple MCP-KO mice using CRISPR/Cas9

Because of the functional redundancy of the MCP members, we sought to generate a knockout (KO) strain devoid of all MCPs by interbreeding each individual MCP-KO lines (*Ccl2^-/-^* X *Ccl7^-/-^* X *Ccl8/12^-/^*^-^). However, this approach proved futile, due to the close linkage of *Ccl2, Ccl7, Ccl8,* and *Ccl12* on chromosome 11, making homology recombination unfeasible. To surmount this obstacle, we designed a strategy to collectively delete all the MCP genes using CRISPR/Cas9-based technology. Combined, these genes span only ∼80k base pairs on chromosome 11 (Fig. 1D). *Ccl11* (Eotaxin) was also deleted because it intercedes the MCPs (Fig. 1D). These *quintuple knockout* (*qMCP^-/-^*) mice were then validated by lipopolysaccharide (LPS, IP, 1 mg/kg) injection into adult mice. Each individual MCP was quantitated in the serum by ELISA, demonstrating an absence of all (Fig. 1E). We used CCL5 as a positive control, whose gene is also located on chromosome 11, but is further away from the MCPs (Fig. 1E). Next, we performed extensive characterization of the brain (Fig. S3), bone marrow (Fig. S4), and spleen (Fig. S5) of healthy non-tumor bearing adult *qMCP*^-/-^ mice by flow cytometry. We did not observe any differences in microglia (Fig. S3B) or bone marrow monocytes (Fig. 2C), but noted an increase in neutrophils in bone marrow (Fig. S4B) and a reduction in total monocytes and Ly6c^Hi^ monocytes in the spleen (Fig. S5B). When we analyzed the absolute count of leukocytes in the blood by flow cytometry (Fig. 1G), there were reduced CD11b^+^ myeloid cells (Fig 1H, P<0.01), likely attributable to the loss of Ly6c^Hi^ inflammatory monocytes (Fig 1H, 31±12 cells/μl in *qMCP^-/-^* vs. 300±113 cells/μl in *WT* mice, P<0.001). No difference in blood neutrophils or lymphocytes was observed (Fig. 1H).

**Figure 2.**
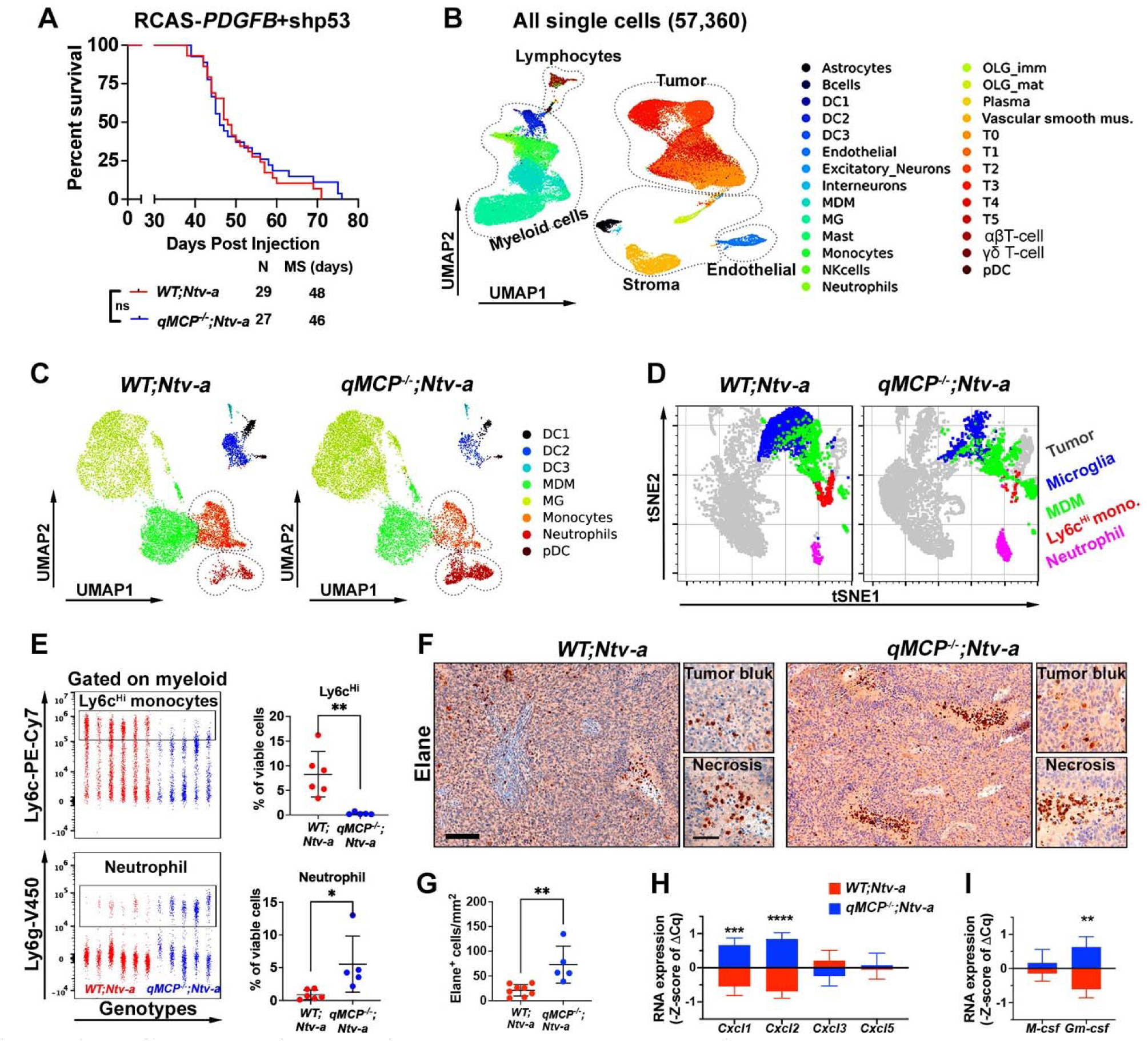
MCP chemokine deletion blocks monocyte recruitment and leads to a compensatory infiltration of neutrophils. (**A**) Survival curves were compared by log-rank test. Ns = not significant. (**B**) UMAP dimensionality reduction of scRNA-seq data from 57,360 cells isolated from three *WT;Ntv-a* and three *qMCP^-/-^;Ntv-a* tumors. Consistent expression of known markers was used to annotate cell clusters into 5 broad cell classes: Lymphoid (B-cells, NK-cells, Plasma, αβT-cell and γδT-cells), Myeloid (DC1, DC2, DC3, pDC, MDM, MG, Monocytes and Neutrophils), Stromal (Astrocytes, Excitatory Neurons, Interneurons, OLG_imm, OLG_mat and vascular smooth muscle cells), Endothelial, and Tumor (T0 to T5). (**C**) UMAP showing refined clustering of myeloid cells isolated from *WT;Ntv-a* (left) and *qMCP^-/-^;Ntv-a* (right) tumors. (**D**) UMAP plots showing results of spectral flow cytometry analysis of tumors. (**E**) Dot-plot and analysis of monocytes and neutrophils analyzed by spectral flow cytometry. Two-tailed Student’s *t*-test. (**F**) Immunohistochemistry staining of Elane in mGBM. (**G**) Quantification of Elane^+^ cells. (**H**) and (**I**) Quantitative analysis of *Cxcl* chemokines or *Csf* by qPCR. Two-tailed Student’s *t*-test. *p<0.05, **p<0.01, ***p<0.001, ****p<0.0001. Scale bar = 100 μm, scale bar in inset = 50 μm. Abbreviations: Mus = muscle; imm= immature; mat= mature.

### Genetic deletion of *qMCP* results in a compensatory influx of neutrophils

Leveraging this new mouse strain, we next sought to determine the role of stroma-derived MCPs in tumor monocyte recruitment. For these studies, we generated GBMs in *WT;Ntv-a* mice with RCAS-shp53 and RCAS-*PDGFB*. When tumors emerged, freshly dissociated tumor cells were orthotopically transplanted into the brains of *qMCP^-/-^* and *WT (B6)* mice (Fig. 1I). Kaplan-Meier analysis demonstrated that eliminating all MCPs from the stroma extended the survival time of tumor-bearing *qMCP^-/-^* mice (Fig. 1J). FACS analysis showed decreased bone marrow-derived myeloid (BMDM) cell infiltration, which likely resulted from decreased Ly6c^Hi^ monocytes (Fig. 1K). When compared to the results using single chemokine KO mice (Fig. S2B), where no reduction of infiltrating monocytes was observed, these results suggest that all MCP members contribute to monocyte recruitment, and that loss of one member can be compensated by other MCPs.

To determine whether survival is extended when all MCPs are genetically ablated in both TME and tumor cells, we induced *de novo* GBM in *WT;Ntv-a* and *qMCP^-/-^;Ntv-a* mice by co-injecting RCAS-*shp53* and RCAS-*PDGFB* in the frontal striatum. We hypothesized that abolishing MCPs would inhibit monocytes tumor infiltration, thereby extend the survival of GBM-bearing mice. Surprisingly, there was no difference in survival between *WT;Ntv-a* and *qMCP^-/-^;Ntv-a* mice (Fig. 2A). To understand the cellular and molecular mechanisms underlying this unexpected result, we analyzed the tumors by single-cell RNA sequencing (scRNA-seq, Fig. S6). After filtering out low quality cells and putative doublets (Methods, Fig. S6A-D), we performed unsupervised clustering on 57,360 cells and identified five major cell classes - lymphoid, myeloid, stromal, endothelial, and malignant (Fig. 2B). Within each class we further stratified cells into phenotypical or functional subsets according to their unique gene signatures (Fig. 2B and Fig. S6E). MCP transcripts can be detected in many cell types in *WT;Ntv-a* mice, particularly malignant cells, macrophages, and monocytes, but were undetectable in *qMCP^-/-^;Ntv-a* mice, reaffirming the efficacy of gene deletion (Fig. S7). Additionally, we found a decrease in monocytes in *qMCP^-/-^;Ntv-a* mice, consistent with this genotype and a corresponding increase of neutrophil infiltration in *qMCP^-/-^;Ntv-a* mice (Fig. 2C).

To complement and corroborate the scRNA-seq data, we used multi-parameter spectral flow cytometry to analyze the composition of myeloid cells in these tumors (Fig. 2D). Based on the combination of multiple surface markers (gating strategy shown in Fig. S8), we identified total myeloid cells (CD11b^+^CD45^+^), which comprise of both brain resident microglia (Mg, CD11b^+^CD45^Lo^Ly6c^Neg^Ly6g^Neg^ CD49d^Neg^) and infiltrating bone marrow-derived myeloid cells (BMDM, CD11b^+^CD45^+^CD49d^+^). These infiltrating myeloid cells were further stratified into inflammatory monocytes (CD11b^+^CD45^Hi^Ly6c^Hi^Ly6g^Neg^CD49d^+^), monocyte-derived macrophages (MOM, CD11b^+^CD45^Hi^Ly6c^Lo/Neg^Ly6g^Neg^CD49d^+^F4/80^+^), or neutrophils (CD11b^+^CD45^+^Ly6c^+^Ly6g^+^CD49d^+^). Quantitatively, we did not observe any difference in total myeloid cells, Mg, or BMDM between the two genotypes (Fig. S8). However, we found a reduction in the presence of Ly6c^Hi^ inflammatory monocytes and an increase in Ly6g^+^ neutrophils in tumors in *qMCP^-/-^;Ntv-a* mice (Fig. 2E). Moreover, in-depth analyses of the lymphocyte compartment (Foxp3^+^ Treg cells, exhausted CD8^+^ T cells, B cells, and NK cells, Fig. S9) and dendritic cells (DCs, DC1 and DC2, Fig. S10) did not reveal any differences between *WT;Ntv-a* and *qMCP^-/-^;Ntv-a* mice.

Next, to confirm this neutrophil infiltration and spatially resolve their presence in tumor tissue, we used immunohistochemistry staining of a neutrophil-specific elastase (Elane; Fig. 2F). We found increased neutrophils in *qMCP^-/-^;Ntv-a* mice (Fig. 2G), consistent with the scRNA-seq and spectral flow cytometry results. Interestingly, these neutrophils tended to cluster around or within the necrotic regions (Fig. 2F), similar to what was recently reported by others (Yee et al., 2020). To determine whether the increased neutrophil influx was associated with increased levels of neutrophil recruitment chemokines, we performed qPCR for *Cxcl1, Cxcl2, Cxcl3,* and *Cxcl5*. Z-score analysis demonstrates significant increases in *Cxcl1* and *Cxcl2* expressions in tumors from *qMCP^-/-^;Ntv-a* mice compared to *WT;Ntv-a* mice (Fig. 2H). Since *Cxcl1* is a major neutrophil recruitment chemokine in mice, these data suggest that increased *Cxcl1* levels may be responsible for the increased neutrophil content in *qMCP*-deficient tumors. In addition, we detected increased granulocyte-macrophage colony-stimulating factor (*Gm-csf*) expression in *qMCP^-/-^;Ntv-a* mice, but not macrophage colony-stimulating factor (*M-csf*) (Fig. 2I). Taken together, abrogating MCP expression results in a near-complete blockade of tumor monocyte recruitment and a compensatory influx of neutrophils, which is associated with increased expression of *Cxcl1* and *Gm-csf*.

While CSF1R inhibition in *PDGFB*-driven GBM-bearing mice (Pyonteck et al., 2013) is ineffective as a monotherapy, the CSF1R inhibitor BLZ945 showed a synergistic effect when combined with radiation (RT) (Akkari et al., 2020). To determine whether abolishing monocytes in combination with RT can also increase anti-tumor efficacy, *WT;Ntv-a* and *qMCP^-/-^;Ntv-a* tumor-bearing mice were treated with irradiation (Fig. S11). Interestingly, no survival advantage was observed with the *qMCP^-/-^;Ntv-a* mice. These different outcomes between CSF1R inhibition and MCP abolition can be partially attributed to the inefficacy of CSF1R-inhibitor in decreasing TAM numbers in GBM; therefore, no compensatory recruitment of neutrophils would be present. The results described here also suggest that compensatory neutrophil recruitment in monocyte-abolished tumors may reverse the synergizing effects of RT, similar to what was recently shown that locally activated neutrophils as a result of irradiation can create a tumor-supportive microenvironment in the lungs (Nolan et al., 2022).

### Human MES tumors show increased neutrophil presence

Increased neutrophil influx is a prominent feature in *qMCP^-/-^;Ntv-a* mice, reminiscent of the mesenchymal (MES) hGBM subtype, which have an abundance of neutrophils (Magod et al., 2021). Using NanoString Pan-Cancer Immune Pathways analyses of hGBM with known molecular subtypes (determined by a custom-made probes for 152 genes from the original GBM_2 design) (Kaffes et al., 2019; Kastenhuber et al., 2014; Omuro et al., 2014), we dissected the cellular landscape of IDH-Mut (G-CIMP) and IDH-WT samples (including PN, CL and MES, Fig. 3A). *In silico* deconvolution of these data showed an increased “neutrophil score” in MES hGBM relative to all other molecular subtypes (Fig. 3B). Consistent with this finding, the neutrophil recruiting chemokine *CXCL8* (P<0.001) and their characteristic surface marker *S100A9* were elevated in MES hGBM, although the latter was not statistically significant (Fig. 3C).

**Figure 3.**
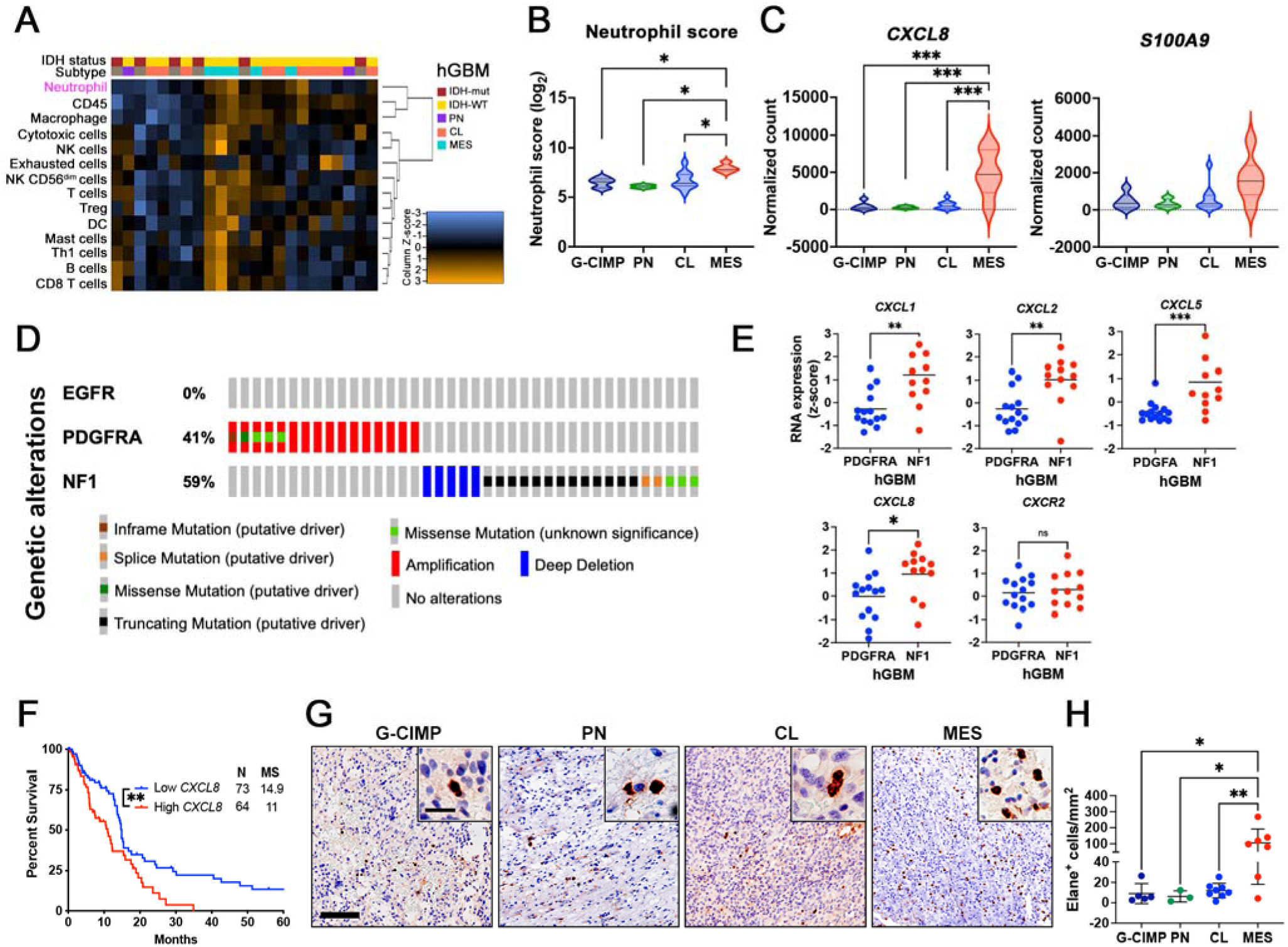
Human MES GBM tumors have increased expression of neutrophil recruitment chemokines and neutrophil content. **(A)**NanoString *in silico* analysis of cellular scores in human GBM tumor samples. **(B)** Neutrophil score in hGBM subtype samples. One-way ANOVA with Tukey’s multiple comparisons test. **(C)** Expression of neutrophil recruitment chemokines *IL8* and *S100A9* examined by NanoString. **(D)** Genetic alterations of GBM patient samples (cBioportal, TCGA, Firehose Legacy) selected based on mutual exclusivity of alterations in PDGFFRA, NF1, and EGFR. **(E)** Expression of neutrophil recruitment chemokines and their shared receptor CXCR2 examined by TCGA. Student’s *t*-test. **(F)** Survival curves of IDH-WT human GBM patients based on low and high expression levels of *IL8*. High and low are defined as +/- 1STDEV from Average of 373 IDH-WT GBM patient samples (cBioportal, TCGA, Firehose Legacy). Log-rank test. **(G)** Representative images of IHC for Elane. (**H**) Quantification of Elane^+^ neutrophils. One-way ANOVA with Tukey’s multiple comparisons test. *p<0.05, **p<0.01, ***p<0.001. Scale bar = 100μm, scale bar in inset = 25 μm. MS = median survival.

To extend our discovery to a larger cohort of GBM patients, we culled data from TCGA (cBioportal, Firehose Legacy set) and filtered the samples using criteria so that they only present alterations in one of the following three driver genes – PDGFRA (driver of the PN subtype), EGFR (CL subtype), or NF1 (MES subtype, Fig. 3D). Since no samples with the EGFR alteration meet these criteria, we performed comparisons between PDGFRA- and NF1-altered patient tumors (Fig. 3D). Similar to the observations above, NF1-altered tumors, which predominantly cluster in MES subtype, showed significantly higher expression of neutrophil recruitment chemokines (*Cxcl-1, -2, -5, -8*) compared to PDGFRA-altered tumors, which predominantly cluster in the PN expression signature group (Brennan et al., 2013; Wang et al., 2017). No difference was observed in chemokine receptor *Cxcr2* (Fig. 3E). Interestingly, when we stratified GBM patients based on their expression of *Cxcl8 (IL8),* a potent neutrophil recruitment chemokine in humans (same method as described in Fig. 1A), increased expression of *Cxcl8* was associated with reduced patient survival (Fig. 3F, P<0.01), analogous to prior reports (Magod et al., 2021).

Finally, to determine whether increased neutrophil recruitment chemokines exist at the protein level, we examined their concentrations in hGBM samples by Olink® proteomic assay (Supplementary Fig. S12). Using a total of 3 IDH-WT MES hGBM tissues and three normal brain samples, we found an increase in chemokines involved in neutrophil recruitment. Furthermore, when we analyzed subtype-defined hGBM samples (Kaffes et al., 2019) by immunohistochemistry for Elane expression (Fig. 3G), there was a similar increase in the number of neutrophils in MES tumors relative to IDH-Mut, PN, and CL GBM samples (Fig. 3H).

### Increased neutrophil influx leads to a tumor transition from PN to MES signature

When GBMs with a PN signature are treated with standard-of-care therapies, they often recur with a MES signature, referred to as PN to MES transition (PN-MES transition) (Fedele et al., 2019). This shift also occurs in *PDGFB*-driven mGBM models in response to anti-VEGFA or RT (Halliday et al., 2014; Pitter et al., 2016). It is interesting to note that neutrophils are highly enriched in *qMCP*^-/-^*;Ntv-a* mice bearing tumors generated with *PDGFB* overexpression, a potent driver mutation of PN GBM. Together, these observations prompted us to determine whether increased neutrophil infiltration can induce PN-MES transition, especially in light of a recent study revealing that reciprocal interactions between TAMs and tumor cells can drive the transition of GBM to a MES-like state (Hara et al., 2021). Based on a previously published algorithm classifying GBM cells into distinct and reproducible cellular states (MES-like, AC-like, NPC-like, and OPC-like) (Hara et al., 2021; Neftel et al., 2019), we analyzed the malignant cells identified in our scRNA-seq datasets for their MES-like state expression score (Fig. 4A). We found a significant increase in MES-like scores in *qMCP^-/-^;Ntv-a* mice relative to *WT;Ntv-a* mice (Fig. 4B, P<0.001), suggesting the tumor cells were undergoing PN-MES transition. To substantiate this result, we leveraged a previously published qPCR panel (Herting et al., 2017) that includes genes associated with either PN or MES subtypes, some of which (e.g., *SERPINE1, CHI3L1*) overlap with the Hara et al. dataset (Hara et al., 2021). As predicted by the observed PN-MES transition, we found increases in many of these MES-related genes (*MGMT, SERPINE1, TAZ, CASP1, TGFB*) in *qMCP^-/-^;Ntv-a* mice (Fig. 4C).

**Figure 4.**
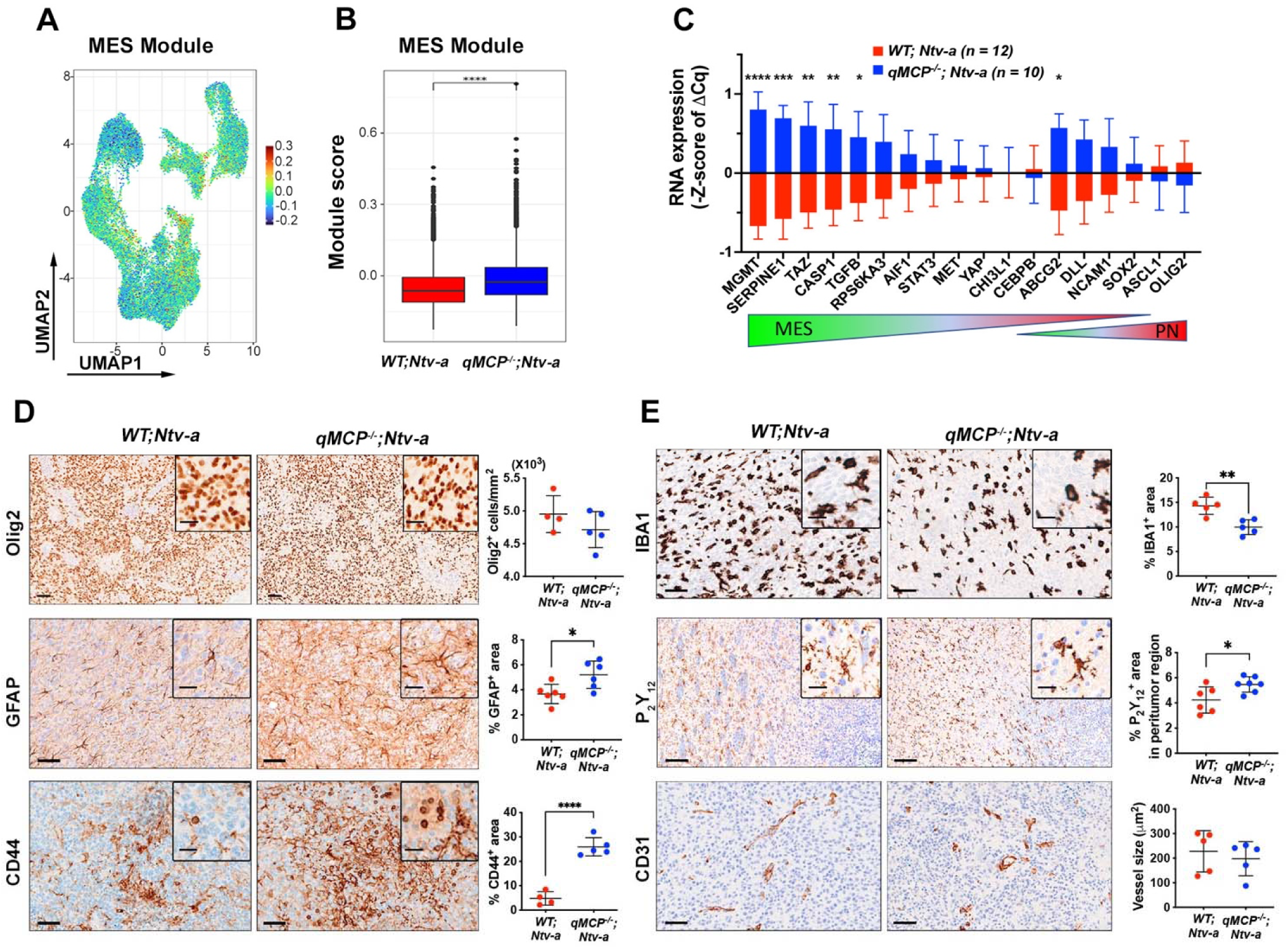
MCP chemokine deletion results in PDGFB-driven GBM PN to MES shift. **(A)** UMAP dimensionality reduction of MES-like cell state module score in all malignant cells examined by scRNA-seq. (**B**) Quantification of MES score between *WT;Ntv-a* (red) and *qMCP^-/-^;Ntv-a* (blue) malignant cells. Student’s *t*-test. (**C**) Real time qPCR panel of signature genes that are differentially expressed in PN and MES mGBM. *P* value was calculated using the Wilcoxon signed-rank test. **(D)** Representative images and quantification of immunohistochemistry for signature molecules of neoplastic cells. Student’s *t*-test. **(E)** Representative images and quantification of immunohistochemistry for molecules in TME. Student’s *t*-test. *p<0.05, **p<0.01, ***p<0.001, ****p<0.0001. Scale bar = 50 μm, scale bar in inset = 20 μm.

Next, we analyzed OLIG2, GFAP, and CD44 expression by immunohistochemistry (Fig. 4D), and found no difference in OLIG2 expression between these two genotypes; however, both GFAP and CD44, canonical markers of MES GBM were increased in *qMCP^-/-^;Ntv-a* mice (Fig. 4D). Taken together, we establish that neutrophil tumor infiltration following monocyte abolition induced PN-MES transition.

In addition to the molecular changes observed within the tumor tissue, we sought to determine whether *MCP* loss changes the cellular composition in the tumor microenvironment (Fig. 4E). IBA1, a pan-macrophage marker, labels TAMs regardless of their origin (microglia, monocytes, and monocyte-derived macrophages). The IBA1-positive areas within the core of the tumors were decreased in tumors generated in *qMCP*^-/-^*;Ntv-a* mice compared to *WT;Ntv-a* mice, as expected for this genotype (Fig. 2E). We also used the microglia-specific marker P_2_Y_12_ to demonstrate increased microglia content at the tumor margins in *qMCP*^-/-^*;Ntv-a* mice (Fig. 4E). No differences in blood vessel sizes (CD31 reactivity; Fig. 4E) were observed.

### Defining the molecular mechanisms underlying the pro-tumor neutrophil effects in GBM

To study the molecular mechanism(s) underlying the tumor-promoting effects of neutrophils in *qMCP^-/-^;Ntv-a* mice, we performed weighted gene co-expression network analysis (WGCNA) on all the malignant cells detected in our scRNA-seq data (Fig. S13). This analysis revealed gene regulatory network, or “modules”, based on gene co-expression patterns (Langfelder and Horvath, 2008), enabling us to identify co-regulated genes shared across multiple cell clusters, which would not be apparent using standard, hard clustering methods implemented in Seurat. Among all the modules examined, the “Greenyellow” module showed prominent increase in *qMCP^-/-^;Ntv-a* mice as quantified by its module score (Fig. 5A). The most co-expressed genes in this module form an interconnected graph (Fig. 5B) and consist of genes implicated in glycolysis (*Gapdh, Pgk1, Pgam1*) and hypoxia (*Aldoa, Mif, Ldha*). To determine the biological functions of WGCNA modules, we performed pathway enrichment analysis using the Hallmark gene sets (Fig. 5C). Among others, we found significant enrichment in glycolysis, hypoxia, and MTOR signaling in the “Greenyellow” module (arrowheads, Fig. 5C), indicating that the tumor cells in *qMCP^-/-^;Ntv-a* mice are likely experiencing a higher metabolic stress. These findings were also recapitulated using GO (gene ontology) pathways (Fig. S14).

**Figure 5.**
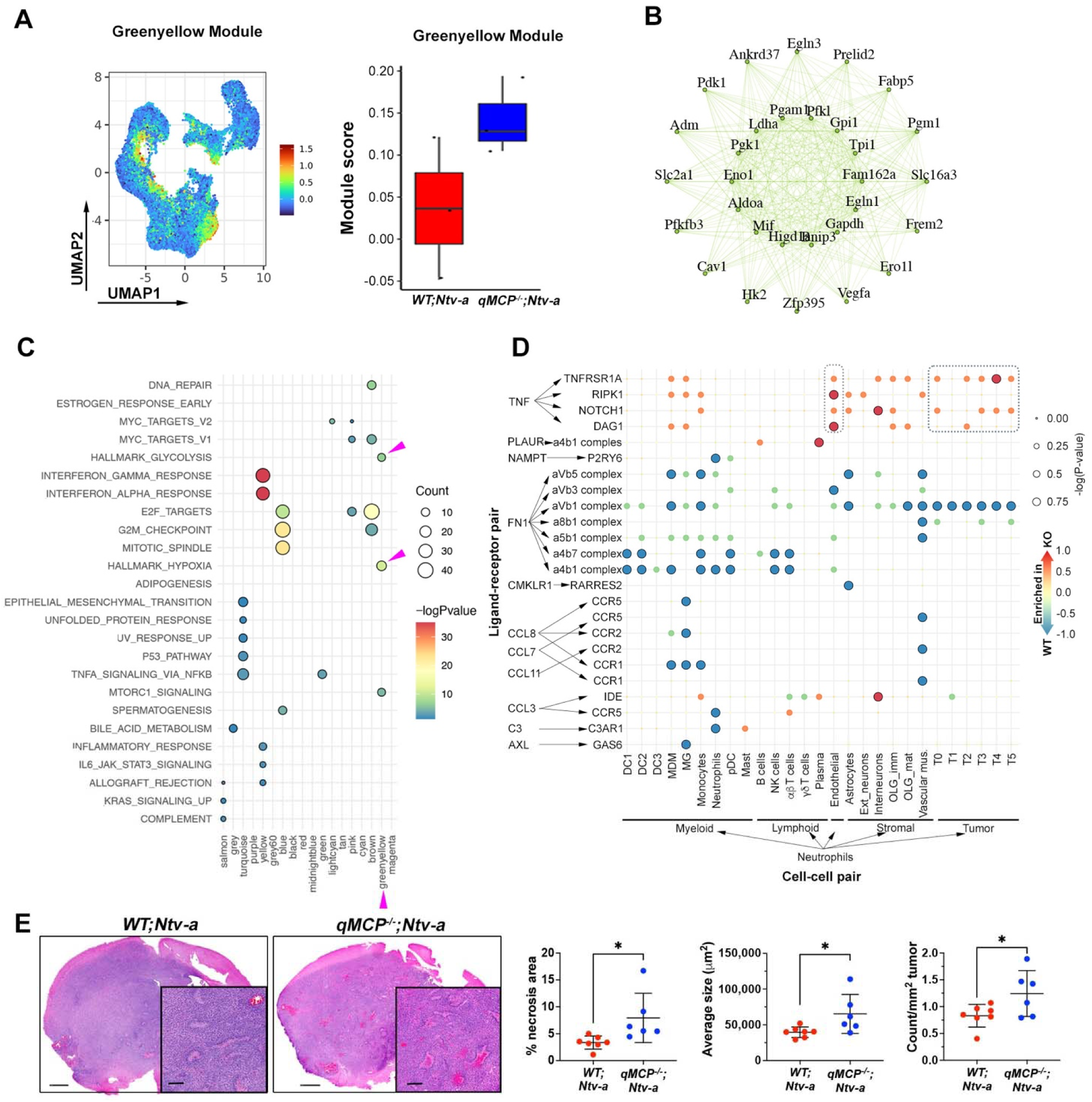
Neutrophils promote tumor progression by inducing a hypoxic response and necrosis. **(A)**UMAP dimensionality reduction of the “Greenyellow” module score identified by scWGCNA analysis (left). Distribution of the average “Greenyellow” module score in malignant cells (right). (**B**) Network graph of the top 30 co-expressed genes in the Greenyellow module. (**C**) Hallmark pathway gene set enrichment analysis of each WGCNA module. Dot colors represent -log(p-value) and dot sizes represent the number of genes in each Hallmark pathway. Arrowheads indicate biological functions related to the Greenyellow module. (**D**) CellphoneDB dot plot showing differentially enriched interactions between ligands (expressed by neutrophil) and receptors (expressed by recipient cells). Dot colors represent the proportion of *WT;Ntv-a* vs. *qMCP^-/-^;Ntv-a* enrichment and dot sizes represent the - log(p-value) of the differential enrichment. Dark circles represent significant interaction (Fisher’s exact test *P* < 0.05). (**E**) Representative images and quantification of H&E staining for necrosis. Student’s *t*-test. *p<0.05. Scale bar = 1 mm, scale bar in inset = 250 μm.

To determine how neutrophils contribute to the metabolic changes in tumors, we performed ligand-receptor interaction inference using CellPhoneDB (Efremova et al., 2020). CellPhoneDB predicts interactions between two cell types based on coordinated expression of ligand-receptor pairs in the respective cell types. We performed this analysis on all cell annotations (Fig. S15), and focused on interactions between neutrophils and all other cell types that are enriched in *qMCP^-/-^*vs. *WT* mice (Fig. 5D). We noted that the neutrophils from tumors generated in *qMCP^-/-^;Ntv-a* mice appeared to be enriched in interactions with many tumor clusters (T0, and T2 to T5) through secretion of tumor necrosis factor α (TNF-α) and signaling via TNF-α receptor-I (TNFR-I: p55) and DAG1 on tumor cells (Fig. 5D). Prior work revealed that TNF-α induces human glioma cell death (Sawada et al., 2004) consistent with our observation that increased necrotic regions exist in tumors from *qMCP^-/-^;Ntv-a* mice. A similar interaction of TNF-α/TNFR-I/II was apparent between neutrophil and endothelial cells (Fig. 5D), which has been shown to be essential for neutrophil transmigration through the endothelial layer (Chandrasekharan et al., 2007), and could potentially explain their increased presence in tumors from *qMCP^-/-^;Ntv-a* mice.

Along with the observations that neutrophils aggregate around necrosis in the tumor tissues (Fig. 2F), the scRNA-seq analysis suggested that neutrophils contribute to tumor progression by facilitating pseudopalisading necrosis formation and increased hypoxic responses. In agreement, we observed increased necrotic areas in tumors from *qMCP^-/-^;Ntv-a* mice (Fig. 5E). In addition, the average size and total occurrence of the necrotic cores were also increased in *qMCP^-/-^;Ntv-a* mice (Fig. 5E).

### Genetic *Cxcl1* loss extends the survival of *qMCP*^-/-^*;Ntv-a* mice

In light of the prominent neutrophil infiltration seen in *qMCP*^-/-^*;Ntv-a* mice, we hypothesized that genetic deletion of *Cxcl1* might reduce neutrophil infiltration and extend survival (Tani et al., 1996). To this end, we first generated GBM in *WT;Ntv-a* and *Cxcl1^-/-^;Ntv-a* mice by co-injecting RCAS-*shp53* and RCAS-*PDGFB* (Fig. 6A, top). However, Kaplan-Meier analysis demonstrated no survival differences between these two genotypes (Fig. 6A, bottom). When we analyzed the myeloid compartment of the TME by spectral flow cytometry (Fig. 6B), no significant changes in infiltrating myeloid cells were observed, although decreased microglia and neutrophil abundance were noted (Fig. 6C). Since microglia and neutrophils account for only a small portion of the myeloid cells in *PDGFB*-driven mGBM, it is not surprising that further reduction of either had no impact on the survival of GBM-bearing mice.

**Figure 6.**
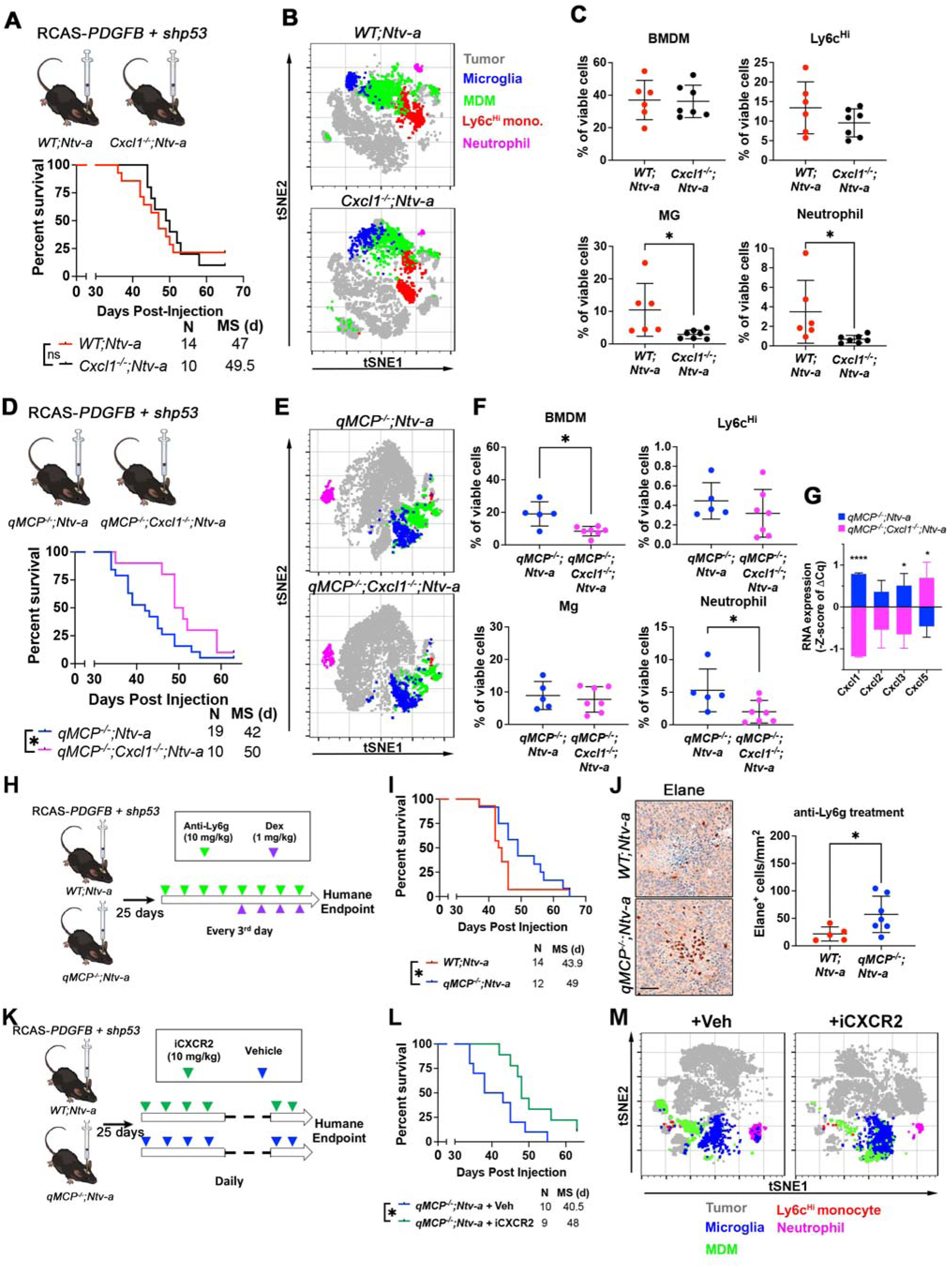
Genetic deletion of *Cxcl1* or pharmacological inhibition of neutrophil extends the survival of *qMCP^-/-^;Ntv-a* mice. **(A)**Schematic illustration and Kaplan Meier-survival curves of *PDGFB*-driven tumors generated in *WT;Ntv-a* and *Cxcl1^-/-^;Ntv-a* mice (*Cxcl1* is lost in both tumor cells and TME). **(B)** tSNE plots illustrating myeloid composition in tumors. **(C)** FACS quantification of myeloid subtypes. Student’s *t*-test. **(D)** Schematic illustration and Kaplan Meier-survival curves of *PDGFB*-driven tumors generated in *qMCP^-/-^;Ntv-a* and *qMCP^-/-^;Cxcl1^-/-^;Ntv-a* mice. **(E)** tSNE plots illustrating myeloid composition in tumors. **(F)** FACS quantification of myeloid subtypes. Student’s *;Ntv-a* -test. **(G)** Real-time qPCR on tumors from **F** at endpoint of survival from. **(H)** Schematic illustration of treatment paradigm using anti-Ly6g antibodies. **(I)** Kaplan-Meier survival curves of *WT;Ntv-a* and *qMCP^-/-^;Ntv-a* mice treated with anti-Ly6g antibodies. Log-rank test. **(J)** Representative images and quantification of Elane in terminal tumors. Student’s *t*-test. **(K)** Schematic illustration of treatment paradigm using CXCR2 antagonist. (**L**) Kaplan-Meier survival curves of *qMCP^-/-^;Ntv-a* mice treated with or without iCXCR2. Log-rank test. (**M**) tSNE plots illustrating myeloid composition in tumors. *p<0.05. BMDM = bone marrow derived myeloid cells, Mg = microglia. iCXCR2 = CXCR2 inhibitor. Dex = dexamethasone. MS = median survival. Scale bar = 50 μm.

Given that neutrophil influx and *Cxcl1* expression were increased in tumors generated in *qMCP*^-/-^*;Ntv-a* mice, we reasoned that genetic deletion of *Cxcl1* in *qMCP*^-/-^*;Ntv-a* mice might reverse these phenotypes and prolong the survival of tumor-bearing mice. *De novo PDGFB*-driven tumors were thus generated in *qMCP*^-/-^*;Cxcl1^-/-^;Ntv-a* mice, resulting in extended survival (Fig. 6D). Spectral flow cytometry analysis of the myeloid cells (Fig. 6E) revealed a reduction in total BMDM infiltrates (Fig. 6F), but Ly6c^Hi^ inflammatory monocytes remained low, comparable to that seen in *qMCP*^-/-^*;Ntv-a* mice (Fig. 6F). Additionally, the infiltrating neutrophils were reduced by *Cxcl1* deletion to less than 50% of that seen in *qMCP*^-/-^*;Ntv-a* mice (Fig. 6F), establishing a critical role for *Cxcl1* in recruiting neutrophil GBM chemoattraction. Next, we examined neutrophil recruitment chemokines by qPCR (Fig. 6G) and observed a reduction in *Cxcl2* and *-3* expression, but a compensatory increase in *Cxcl5* (Fig. 6G). It is interesting to note that neutrophil recruitment chemokines are localized adjacent to each other on chromosome 5, reminiscent of the MCPs, suggesting a similar compensatory mechanism might also exist.

### Pharmacological targeting of neutrophils prolongs the survival of *qMCP*^-/-^*;Ntv-a* mice

In light of the finding that *Cxcl1* deletion reduces GBM neutrophil infiltration and extends survival, we sought to determine whether pharmacological inhibition of neutrophils would produce the same effects. For these experiments, we employed two strategies to reduce neutrophils. First, we adopted the widely used anti-Ly6g antibody neutrophil depletion paradigm. Starting day 25 after tumor initiation until the humane endpoint, we injected anti-Ly6g antibodies (IP, 200 µg/mouse, once every third day) to deplete neutrophils from the circulation and from the tumors (Fig. 6H). Following anti-Ly6g treatment, the survival time of tumor-bearing *qMCP*^-/-^*;Ntv-a* mice was prolonged relative to *WT;Ntv-a* controls (Fig. 6I). Surprisingly, when we quantified Elane^+^ neutrophils at the endpoint of the survival experiment, neutrophils remained elevated in *qMCP*^-/-^*;Ntv-a* mice (Fig. 6J). As recently documented, anti-Ly6g neutrophil depletion effects are transient, occurring only at the initial stage of treatment, followed by an effective rebound (Boivin et al., 2020; Yee et al., 2020). To better understand the dynamics of neutrophil response to Ly6g depletion, we collected blood from naïve animals treated with anti-Ly6g antibody at various time points and analyzed blood neutrophil counts using a Cytospin assay (Fig. S16A). In keeping with a prior report (Yee et al., 2020), our results showed a transient depletion and a gradual rebound of neutrophils in the blood (Fig. S16B), accounting for our findings in the tumor tissues (Fig. 6J). It is important to note that, independent of genotype, mice treated with anti-Ly6g antibody exhibit treatment-induced seizures after 4 doses, which we attributed to increased systemic inflammation. To counteract this adverse effect, we administered dexamethasone at 1 mg/kg (IP, every third day) (Fig. 6H).

Second, we inhibited neutrophil recruitment using a CXCR2 antagonist – a potent and selective small molecule inhibitor (iCXCR2) – SB225002 (White et al., 1998). CXCR2 is the only functional receptor for Cxcl-1, -2, -5, and -15 in mice, where it is crucial for neutrophil recruitment and the regulation of vascular permeability (Belperio et al., 2002; Cao et al., 2018; Mei et al., 2012). The potency of SB225002 in inhibiting neutrophil chemotaxis *in vitro* (White et al., 1998) and *in vivo* (Cao et al., 2018; Yellowhair et al., 2019) had been demonstrated in various disease contexts, including cancer (Kumar et al., 2017). GBM-bearing *qMCP*^-/-^*;Ntv-a* mice were treated with either vehicle or iCXCR2 twenty-five days after tumor initiation (Fig. 6K), resulting in increased survival (Fig. 6L). We performed a comprehensive analysis of myeloid composition by spectral flow cytometry (Fig. 6M). Similar to anti-Ly6g treatment, no difference in neutrophil numbers was observed in iCXCR2 treated tumors at the terminal stage of cancer (Fig. 17). Collectively, these results demonstrate that strategies aiming at reducing neutrophils in *qMCP*^-/-^*;Ntv-a* mice only transiently prolong survival as a result of rebound neutrophil infiltration.

### Abrogation of neutrophil, but not monocyte, infiltration increases the survival of HCC-bearing mice

To determine whether the compensatory recruitment of neutrophils is a CNS- and/or GBM-specific phenomena, we employed a GEMM model of hepatocellular carcinoma (HCC). In contrast to GBM, HCC tumors are mainly populated by neutrophils, mirroring the monocyte:neutrophil ratio (1:3) seen in the blood of WT animals (Fig. 1H). Two of the most frequently altered genes in human HCC are *MYC* (amplified in 17% of HCCs) and *TP53* (deleted or mutated in 33% of HCCs) (Cancer Genome Atlas Research Network. Electronic address and Cancer Genome Atlas Research, 2017), where they tend to co-occur (Cancer Genome Atlas Research Network. Electronic address and Cancer Genome Atlas Research, 2017; Ruiz de Galarreta et al., 2019). For this reason, we generated murine HCCs with MYC overexpression and TP53 loss by hydrodynamic tail vein injections of a transposon vector co-expressing MYC and luciferase (*MYC-Luc*) and a CRISPR vector expressing a sgRNA targeting *p53 (sg-p53)* (Bollard et al., 2016; Ruiz de Galarreta et al., 2019) in *qMCP*^-/-^ (abrogated monocyte infiltration), *Cxcl1^-/-^* (decreased neutrophil infiltration), and *WT* C57BL/6 mice (Fig. 7). Because of the known difference in median survival between male and female mice in this HCC model, we stratified the mice by gender and analyzed each sex separately. Liver luciferase expression measured by bioluminescence imaging (BLI) at day 7 demonstrates equal intensity between all genotypes, revealing similar *in vivo* transfection efficacy of the plasmids (Fig. 7A, B).

**Figure 7.**
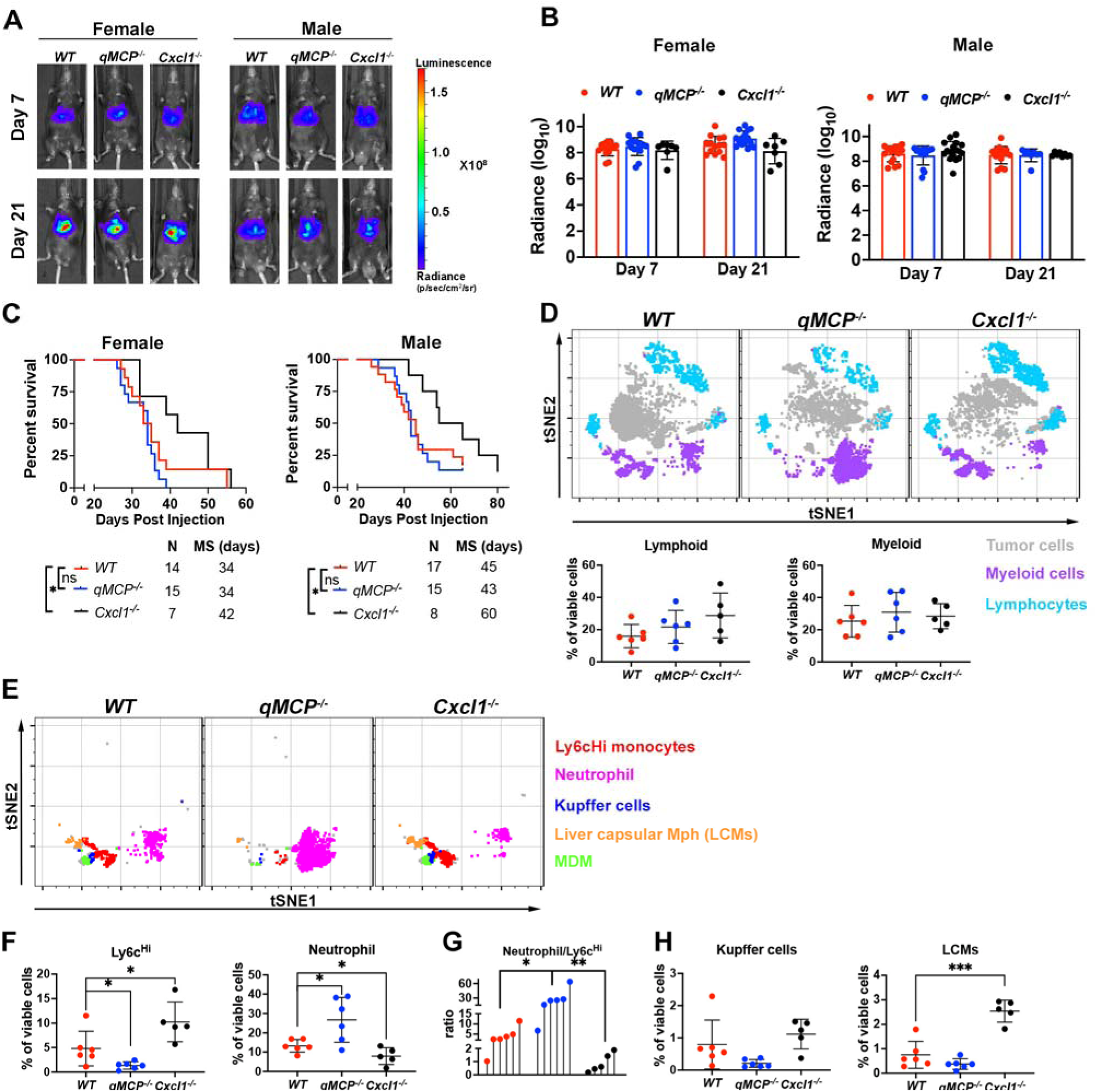
Blocking HCC neutrophil, but not monocyte, recruitment decreases tumor growth and mouse survival. **(A)** Representative images and **(B)** corresponding quantification graphs of bioluminescence imaging at 7 and 21 days after tumor initiation. **(C)** Kaplan-Meier survival curves for HCC-bearing female (left) and male (right) WT, *qMCP^-/-^, and Cxcl1^-/-^* mice. Log-rank test. **(D)** tSNE plots and flow cytometry quantification of lymphoid and myeloid cells in HCC-bearing mice. **(E)** tSNE plots illustrating myeloid cells examined by spectral flow cytometry. (**F**) Quantification of monocytes and neutrophils by spectral flow cytometry. One-way ANOVA with Tukey’s *post-hoc* test. (**G**) Lollipop plot showing neutrophil to Ly6c^Hi^ monocytes ratio. One-way ANOVA with Tukey’s post-hoc test. (**H**) Quantification of Kupffer cells and LCMs by spectral flow cytometry. One-way ANOVA with Tukey’s *post-hoc* test. *p<0.05, **p<0.01, ***p<0.001. Ns = not significant. MS = median survival.

No differences in BLI signal were observed 21 days post-injection (Fig. 7A, B). While Kaplan-Meier survival analysis demonstrated that deleting all *MCP* genes had no effect on survival time of HCC mice, knocking out *Cxcl1* extended survival, regardless of gender (Fig. 7C). Because of similar outcomes between males and females, subsequent studies were performed only using male mice. No significant differences in total myeloid and lymphoid cell populations were observed between genotypes (Fig. 7D). However, when we specifically evaluated the myeloid compartment of HCC by spectral flow cytometry (Fig. 7E, gating strategies in Supplementary Fig. S18), there was a reduction of monocytes and an increase in neutrophil infiltration in *qMCP^-/-^*tumors (Fig. 7F). These results suggest that, similar to *PDGFB*-driven-GBM, abolishing monocyte recruitment leads to compensatory neutrophil infiltration in HCC, but confers no effect on survival. We also observed decreased neutrophil infiltration which was associated with a compensatory increase in monocyte infiltration in *Cxcl1*-deficent tumors (Fig. 7F) and a decreased neutrophil to monocyte ratio (Fig. 7G).

Whereas no difference in Kupffer cells (KCs) was observed (Fig. 7H), *Cxcl1* deletion resulted in increased liver capsular macrophages (LCMs) (Fig. 7H), consistent with prior reports demonstrating that LCMs are replenished from blood monocytes, while KCs are embryonically derived and capable of self-renewal *in situ* (Sierro et al., 2017). In-depth analysis of the lymphoid compartment (Fig. S19) showed no changes between the three genotypes (Fig. S20). Taken together, these observations establish that compensatory infiltration of monocytes and neutrophils is not specific to the CNS, but rather a generalized phenomenon in cancer.

## DISSCUSION

Despite intensive efforts over the last decade, modulating tumor-associated myeloid cells to treat solid tumors, including GBM, has proven exceptionally difficult. This is in large part due to an incomplete understanding of the immune cell heterogeneity during tumor progression and treatment. A classic example is CSF1R antagonism, where its variable therapeutic efficacy is heavily impacted by tumor type and/or models studied (O’Brien et al., 2021; Pyonteck et al., 2013) (Butowski et al., 2016) (Maximov et al., 2019) (Tan et al., 2021). These studies suggest that TAMs evolve and attain independence from CSF1R inhibition as diseases progress, thereby become elusive to therapy. To gain insights into the complexity of immune cell responses following myelo-inhibition, we utilized numerous complementary approaches in this study, leveraging unique chemokine knockout GEMMs, scRNA-seq, human transcriptomics and proteomics data, and two different murine tumor models (GBM and HCC).

First, we demonstrate that genetic loss of individual MCP genes in stromal cells results in improved survival of tumor-bearing mice with no change in TAM content, while loss of individual MCPs from both stromal *and* tumor cells abolishes survival advantage (Fig. S2). These results suggest a redundancy in function of MCP members and compensatory changes in monocyte populations following their elimination. It also motivated us to create a mouse model that is deficient of all MCP members. As expected, depletion of all MCPs from both the stroma and the tumor compartments abolishes monocyte recruitment. To our surprise a neutrophil influx and a concomitant rise in neutrophil chemotactic cytokines Cxcl-1 and -2 accompanied monocyte depletion (Fig. 2). In contrast to monocytes, due to their lower abundance in GBM, the role of neutrophils has remained less known (Lin et al., 2021). Recent studies have demonstrated that neutrophil infiltration and activation are markers of poor glioma prognosis (Rahbar et al., 2016). Increased neutrophil degranulation, elevated levels of ARG1 that suppress T-cell functions, upregulation of S100A4 expression, and increased IL-12 levels have been shown to be associated with glioma malignancy (Liang et al., 2014; Rahbar et al., 2016). Together, these observations elucidate the lack of difference in survival duration between WT and *qMCP*-deficient mice in this study.

Functional investigation of neutrophil in GBM, or in cancer in general, is a nascent field (Quail et al., 2022). It has been shown at the onset of GBM formation, neutrophils have an anti-tumoral effect, but adopt a tumor-supportive phenotype as tumor progression occurs (Magod et al., 2021). One of the mechanisms by which neutrophils exert their pro-tumorigenic function in GBM is via induction of ferroptosis and tumor necrosis (Yee et al., 2020). Similarly, spatial analysis of tumor-associated neutrophil by IHC in this study reveals that the majority of these cells gather in or around necrotic areas

(Fig. 2). Inference of ligand-receptor pairs from scRNA-seq data suggest that neutrophils release TNFα in a paracrine fashion, which likely facilitates tumor necrosis and a hypoxic response (Fig. 5). Future studies should address whether inhibiting TNF in the TME or ablating TNF receptors on tumor cells can hamper neutrophil pro-tumorigenic functions.

We and others have demonstrated that murine GBM models closely resembling the human MES GBM subtype have increased numbers of neutrophils relative to PN- and CL-like GBM models (Chen et al., 2020; Magod et al., 2021). Our findings that *PDGFB*-driven tumors in *qMCP*-deficient mice have increased expression of MES genes, increased expression of MES marker CD44, and increased hypoxia upon IHC staining are consistent with a transition from a PN to a MES GBM subtype (Fig. 4) (Hara et al., 2021). To evaluate clinical relevance of our observations, we analyzed NanoString data of IDH-WT and IDH-MUT tumors for neutrophil and their recruitment chemokines. In agreement, we find an increased presence of neutrophil and neutro-attractant chemokines in human MES subtype samples (Fig. 3). When a larger patient cohort from TCGA was stratified based on their IL-8 (*CXCL8*) expression, higher expression is found to be associated with inferior survival. Overall, our results indicate that monocyte:neutrophil ratio can define tumor signature, highlighting the essential roles these two cell types play in shaping tumor cell expression profiles and crafting the evolving TME.

It is interesting to note that depleting one pro-tumorigenic myeloid subset resulted in an influx of an equally pro-tumorigenic subset. Similar observations were documented following CSF1R inhibition in transplantable solid tumor models and patient samples outside of the CNS (Kumar et al., 2017). In contrast, CSF1R inhibition in a GBM mouse model did not reduce the number of TAMs, and thereby did not induce compensatory neutrophil influx (Coniglio et al., 2012). While CSF1R inhibitors target TAMs independent of their ontogeny, we show in this study that the compensatory neutrophil recruitment in GBM is driven by abolishing monocyte infiltration without affecting resident brain microglia.

We were curious whether decreasing neutrophil infiltration in neutrophil-enriched tumors would lead to increased monocyte infiltration. Therefore, as a proof-of-principle we selected a well-documented neutrophil-enriched tumor outside of the CNS – HCC. It is well established that neutrophil numbers can serve as powerful predictors of poor outcome in HCC patients, but mechanisms of their infiltration remain elusive (Kuang et al., 2011; Margetts et al., 2018). By using a genetic mouse model, we demonstrate that similar to human HCC, murine HCC are enriched with neutrophils and their ratio to monocytes (∼3:1) mirror that of the blood of healthy WT mice, contrasting to GBM. Abolishing monocytes had no impact on survival of neutrophil-enriched HCC tumor-bearing mice, similar to what we had shown with abolishing neutrophils in monocyte-enriched *PDGFB*-GBM. Decreased neutrophil infiltration resulted in extended survival and was associated with increased monocyte recruitment, suggesting this compensatory mechanism exists both ways.

An intriguing question remains: why are some tumors enriched in monocytes while others neutrophils? Monocytes are a minority in blood circulation, yet they give rise to the dominant infiltrates in GBM; whereas neutrophils, the most abundant in the blood, rarely invade GBM (except for MES tumors in both human and mice)? It is speculated that *Tp53* loss, either alone or in combination with *Kras* or *Pten*, create microenvironments that preferentially favor neutrophil infiltration in various solid tumors (Quail et al., 2022). However, *Tp53* mutation occurs at a very high prevalence across many cancer types, ranging from 30% to 47% in brain, liver, lung, skin, ovarian and many other cancers (Olivier et al., 2010). Therefore, this widespread presence cannot explain the highly diverse TME across all the tumor types. For instance, both GBM and HCC models used in this study involved *Tp53* silencing, however their TMEs are contrasting in terms of monocytes/neutrophil compositions. We posit that unique combinations of driver mutations in different tumor types, rather than loss of a universal tumor suppressor gene, play a decisive role in this phenomenon. We have shown that *Nf1*-silenced murine and human GBMs have increased expression of neutrophil recruitment chemokines compared to *PDGFRA* amplified hGBM and *PDGFB*-driven mGBMs, which favor monocyte infiltration (Chen et al., 2020; Kaffes et al., 2019). Therefore, in-depth analysis of tumor samples created by different driver mutations will uncover potential mechanisms each tumor uses to construct their distinctive TMEs.

In conclusion, we demonstrated that when all MCPs were genetically deleted and monocyte recruitment abolished, GBM adapts to mobilize an influx of neutrophils. Similar compensatory effects exist in HCC. These observations explain the failure of current treatment attempts that pursue single chemokines. It is therefore imperative to develop combinatory therapies that are simultaneously directed at both monocytes and neutrophils. Effective treatment can also be confounded by the complexity where targeting neutrophil influx are often challenged by a rebound effect. Fundamental understanding of the interplay between driver mutations, monocytes, neutrophils, and other TME cell types using state-of-the-art GEMMs, single-cell resolution measurements, and integrative analysis as utilized in our study will be critical to future pharmacological development aiming at creating long lasting, dual function compounds.

## METHODS

### Generating qMCP knock-out mice

The qMCP knock-out mice were generated at the Mouse Transgenic and Gene Targeting Core at Emory University. A pair of guide RNAs (upstream: CCCTGGCTTACAATAAAAGGCT, and downstream: CAGCAGGCCAAATGAGGGGAGG) were designed to recognize the 81k base DNA segments flanking the genes between *Ccl2* and *Ccl8* (inclusive) on chromosome 11. The guide RNAs were synthesized and validated by Sigma (MilliporeSigma). The guide RNAs, CRSPR/cas9 mRNA, and a donor repair oligonucleotide (5’-TCACTTATCCAGGGTGATGCTACTCCTTGGCACCAAGCACCCTGCCTGACTCCACCCCCCAGGTGTTCAAGGGTTCCTGTGTATTATTTGGGTTTCATTTTATGGGGTTCAAGTGAAGGA-3’) were co-injected into C57BL/6N (RRID:MG:6198353) zygotes and transferred to surrogate dames. Two founder female mice were identified by PCR, and verified by DNA sequencing. We back crossed founder #5 to the C57BL/6J strain for over 10 generations and the progeny’s genetic background was confirmed to be C57BL/6J via genetic monitoring service provided by Transnetyx. All the mice were viable and fertile. All subsequent genotyping was done at Transnetyx with the probe set named Gm17268-1.

### Mice used in this study

Mice of both sexes (equal distribution) in the age range of 6-12 weeks were used for experiments (Chen et al., 2020; Herting et al., 2017). Previously-described *Ccl2^-/-^* (Jackson laboratory, #004434), *Ccl7^-/-^* (Jackson laboratory, #017638), *Ccl8/12^-/-^* (gifted by Dr. Sabina Islam), *Cxcl1^-/-^* (Shea-Donohue et al., 2008), and *Cxcr2^-/-^* (Jackson laboratory, #006848) mice were either maintained as single knock-out strains, or cross-bred to the *Ntv-a* mice to generate double or triple knock-out strains. All these mice are in a C57BL/6 background. C57BL/6J mice (#000664) at 6 weeks old were purchased from the Jackson labs. All animals were housed in a climate-controlled, pathogen-free facility with access to food and water *ad libitum* under a 12-hour light/dark cycle. All experimental procedures were approved by the Institutional Animal Care and Use Committee (IACUC) of Emory University (Protocol #2017-00633) and the Icahn School of Medicine at Mount Sinai (Protocol #2019-00619 and #2014-0229).

### RCAS virus propagation to generate *de novo* GBM

DF-1 cells (ATCC, CRL-12203, RRID:CVCL_0570) were purchased and grown at 39°C according to the supplier’s instructions. Cells were transfected with RCAS-*hPDGFB*-*HA*, RCAS-*hPDGFA*-myc, RCAS-shRNA-*p53*-RFP, RCAS-shRNA-*Nf1*, RCAS-shRNA-*Pten*-RFP, and RCAS-*Cre* using a Fugene 6 transfection kit (Roche, 11814443001) according to the manufacturer’s instructions. DF-1 cells (4x10^4^) in 1 µl neurobasal medium were stereotactically delivered with a Hamilton syringe equipped with a 30-gauge needle for tumor generation (Franklin and Paxinos, 1997). The target coordinates were in the right-frontal striatum for PDGFB-overexpressing tumors at AP -1.5 mm and right -0.5 mm from bregma; depth -1.5 mm from the dura surface. NF1-silenced tumors were generated via injection into the subventricular zone at coordinates AP -0.0 mm and right -0.5 mm from bregma; depth -1.5mm from the dura surface (Chen et al., 2020; Franklin and Paxinos, 1997; Herting et al., 2017). Mice were continually monitored for signs of tumor burden and were sacrificed upon observation of endpoint symptoms including head tilt, lethargy, seizures, and excessive weight loss.

### Orthotopic glioma generation

The same procedure was used as described above, except 3X10^4^ of freshly-dissociated tumor cells were injected in the right-frontal striatum AP -1.7mm and right -0.5mm from bregma; depth -1.5mm from the dural surface of recipient animals. Two or three donor tumors of either sex were used for obtaining single cells for orthotopic glioma generation in male and female recipient animals.

### Hydrodynamic tail-vein injection to generate HCC

A sterile 0.9% NaCl solution/plasmid mix was prepared containing DNA. We prepared 11.4 μg of pT3-EF1a-MYC-IRES-luciferase (MYCluc), 10 μg of px330-sg-p53 (sg-p53), and a 4:1 ratio of transposon to SB13 transposase–encoding plasmid dissolved in 2 mL of 0.9% NaCl solution and injected 10% of the weight of each mouse in volume as previously described (Ruiz de Galarreta et al., 2019). Because two independent “hits” are required for tumor formation in C57BL/6 mice (Chen and Calvisi, 2014), only those hepatocytes that receive the three plasmids (transposon-based, transposase, and CRISPR-based) will have the potential to form tumors. Mice were injected with the 0.9% NaCl solution/ plasmid mix into the lateral tail vein with a total volume corresponding to 10% of body weight in 5 to 7 seconds. Vectors for hydrodynamic delivery were produced using the QIAGEN plasmid PlusMega kit (QIAGEN). Equivalent DNA concentration between different batches of DNA was confirmed to ensure reproducibility among experiments.

### Luciferase Detection

In vivo bioluminescence imaging was performed using an IVIS Spectrum system (Caliper LifeSciences, purchased with the support of NCRR S10-RR026561-01) to quantify liver tumor burden. Mice were imaged 5 minutes after intraperitoneal injection with fresh d-luciferin (150 mg/kg; Thermo Fisher Scientific). Luciferase signal was quantified using Living Image software (Caliper LifeSciences, RRID:SCR_014247). Normalized luciferase signal was calculated by subtracting the background signal. Those mice with a luciferase signal a log of magnitude lower than the average signal were excluded from the study.

### Tumor and cultured cell RNA isolation and analysis

At humane endpoint, mice were sacrificed with an overdose of ketamine and immediately perfused with ice cold Ringer’s solution (Sigma-Aldrich, 96724-100TAB). The brain was extracted, and a piece of tumor was immediately snap-frozen in liquid nitrogen for storage at -80°C. Alternatively, cultured cells were harvested from plates using TRIzol (ThermoFisher, 15596026). RNA was isolated from the frozen tumor pieces or cells with the RNeasy Lipid Tissue Mini Kit (Qiagen, 74804) according to the manufacturer’s instructions. RNA quantity was assessed with a NanoDrop 2000 spectrometer, while quality was confirmed via electrophoresis of samples in a 1% bleach gel as previously described (Aranda et al., 2012). RNA was used to generate cDNA with a First Strand Superscript III cDNA synthesis kit (ThermoFisher, 18080051) according to the manufacturer’s instructions and with equal amounts of starting RNA. Quantitative-PCR was performed with the validated BioRad PCR primers (Table S2, except *Cxcl1* and *Cxcl2* whose sequence were obtained from Girbl et al (Girbl et al., 2018)). using SsoAdvanced Universal green Supermix (BioRad, 1725271). Fold changes in gene expression were determined relative to a defined control group using the 2^-ΔΔCt^ method or by z-score, with β-Actin or HPRT used as housekeeping genes.

#### NanoString analysis

Human formalin-fixed, paraffin-embedded (FFPE) tissue scrolls were cut in 10µm sections for RNA extraction using the RNeasy Lipid Tissue Mini Kit (Qiagen #74804) according to the manufacturer’s instructions. RNA integrity was confirmed using a bioanalyzer and samples possessing a DV300% greater than 30 were used. 50 ng of RNA was used for NanoString analysis with the pan-cancer pathways immune panel (NanoString, XT-CSO-HIP1-12). All data analysis was processed and normalized using nSolver Analysis Software version 4.0 (NanoString) and GraphPad Prism 9 (GraphPad Software, RRID: SCR_002798).

#### Neutrophil morphology analysis by Cytospin

Whole blood was collected with a heparinized capillary tube (Fisher, 22-362-566) via mandibular vein puncture. 10 μl of whole blood was suspended in 1 ml RBS lysis buffer in a 1.5 ml Eppendorf tube, which was kept at 37 C for 1 min. The tube was centrifuged at 450g for 4 min at RT and the pellet was resuspended in 100 μl PBS containing 1% BSA. Cytospin slides (ThermoFisher, 5991056) and filter paper (ThermoFisher, 5991022) were assembled according to manufacturer’s instructions. The cell suspension was then transferred to the funnel that was attached to the slides. The assembly was loaded into the Cytospin centrifuge and run at 800 rpm for 5 min. The slides were air-dried at RT for 20 min before they were plunged into ice cold methanol for 5 min. The slides were then stained with DAPI (Sigma, 1 ug/ml) for 10 min at RT in the dark. Images of the nuclei were taken with a Leica confocal microscope (Leica, SP8) with a 10X objective, with multiple areas of view were acquired for each sample. Number of nuclei with typical neutrophil nuclear morphology (Fig. 7D) was counted.

#### Human tissue samples and pathological appraisal

Human FFPE GBM samples, post-mortem brain specimens, and de-identified clinical information were provided by Emory University. Board-certified neuropathologists graded and diagnosed both the human tumor tissues and murine samples according to the 2016 World Health Organization Classification of Tumors of the Central Nervous System (Louis et al., 2016). Gene expression profiling to determine transcriptional subtypes was performed using NanoString nCounter Technology using custom-made probes for 152 genes from the original GBM_2 design (Kaffes et al., 2019). Flash frozen, de-identified GBM samples and adjacent non-malignant tissues acquired during tumor resection were obtained from Mount Sinai Hospital through the biorepository, under IRB-approved protocols (18-00983). Whole genome sequencing was performed to determine patients’ IDH mutation status and molecular genotype of the tumors.

#### TCGA analysis

U133 Microarray data for the GBM (TCGA, provisional) dataset were downloaded from cBioPortal (https://www.cbioportal.org, RRID:SCR_014555) in August 2019 and sorted into subtypes based upon a proprietary key. G-CIMP-positive tumors were excluded from analysis. We included 372 patient samples for which covariate information (survival information, age, and gender) was available. Cox Proportional Hazard Models were fitted in R using age and gene expression as continuous covariates, and gender as a binary variable.

#### Tissue processing and immunohistochemistry

Archived FFPE human GBM samples and de-identified clinical information were provided by Emory University. Murine FFPE sapeles were generated as previously described (Chen et al., 2017). The specimens were sectioned at 5 μm thickness, slide-mounted, and stored at -80°C until use. To process mouse tumor tissues, animals were anesthetized with an overdose of ketamine/xylazine mix and transcardially perfused with ice-cold Ringer’s solution. Brains were removed and processed according to the different applications. For H&E tumor validation and immunohistochemistry staining, brains were fixed in 10% neutral buffered formalin for 72 hours at room temperature (RT), processed in a tissue processor (Leica, TP1050), embedded in paraffin, sectioned at 5 μm with a microtome (Leica), and mounted on superfrost glass slides (ThermoFisher 3039-002). Slides were rehydrated with tap water and dipped in hematoxylin (ThermoFisher, 7231), bluing agent (ThermoFisher, 22-220-106) and eosin (ThermoFisher, M1098442500) for 1 min each with thorough washes with tap water in-between. Slides were dehydrated with series washes in ethanol and Neo-clear (ThermoFisher, M1098435000) before mounted in Permount medium (ThermoFisher, SP15-100).

All immunohistochemistry staining was performed on a Leica Bond Rx platform (Leica). Primary antibodies (a full list of primary antibodies used in this study is shown in Table S3) used in this study include: anti-IBA-1 (1:1,500, FUJIFILM Wako, 019-19741, RRID:AB_839504), anti-CD31 (1:50, Dianova, DIA-310), anti-CD44 (1:100, BD Pharmingen, 550538, RRID:AB_393732), and anti-OLIG2 (1:500, Millipore, AB9610, RRID:AB_570666). Anti-GFAP (1:10,000, CST, 3670, RRID:AB_561049), anti-Elastase (1:400, Bioss, bs6982R or 1:400, AbCam, ab68672), anti-P2Y12 (1: 500, AnaSpec, SQ-ANAB-78839, discontinued). Appropriate secondary antibodies were purchased from Leica or Vectorlab. Digital images of the slides were acquired by using a Nanozoomer 2.0HT whole-slide scanner (Hamamatsu Photonic K.K) and observed offline with NDP.view2 software (Hamamatsu). Image analysis was performed using Fiji (NIH, RRID:SCR_002285).

#### Tumor dissociation and primary cell culturing

Tumor dissociation was performed as previously described. Briefly, tumors were dissected from the brain, minced into pieces < 1 mm^3^, and digested with an enzymatic mixture that includes papain (0.94 mg/ml, Worthington, LS003120), EDTA (0.18 mg/ml, Sigma, E6758), cystine (0.18 mg/ml, Sigma, A8199), and DNase (60 μg/ml, Roche, 11284932001) in 2 ml HBSS (Gibco, 14175-095). Tumor tissues were kept at 37°C for 30 minutes with occasional agitation. The digestion was terminated with the addition of 2 ml Ovomucoid (0.7 mg/ml, Worthington, LS003086). Following digestion, single cells were pelleted, resuspended in HBSS, and centrifuged at low speed (84 RCF) for 5 min, before passing through a 70 μm cell strainer.

#### Anti-Ly6g antibody or iCXCR2 treatment

Tumor-bearing mice were randomly assigned to different experimental groups on the first day of treatment. For neutrophil depletion, tumor-bearing mice received intraperitoneal injections of 200 µg of 1A8 (Ly6g depletion, BE0075-1) or 2A3 (control; BE0089, both from Bio X Cell) antibody per mouse starting from day 25 after DF1 cell injection. Injections were given every day until mice succumb to disease and were sacrificed at humane endpoints. Mice were monitored for signs of disease progression as described above. To decrease treatment induced seizures, mice were given 1mg/kg Dexamethasone (West-ward) every third day starting day 37.

CXCR2 inhibitor SB 225002 (iCXCR2) was purchased from Tocris (#2725) and dissolved in DMSO to make 10 mg/ml solution. This solution is diluted 10 X with 0.33% Tween80 (v/v) in saline on the day of treatment. Starting on day 20 after tumor initiation, each mouse assigned to the treatment group received an IP injection of iCXCR2 at 10 mg/Kg daily until humane endpoint.

#### Olink multiplex proteomic analysis

Flash frozen human GBM samples and adjacent non-malignant tissues were weighed and ∼ 30 mg of tissues from each sample were transferred to a 1.5 ml Eppendorf tube. T-PER Tissue Protein Extraction Reagent (ThermoFisher, 78510) containing phosphoSTOP and protein inhibitors cocktail (100 mg/ml, Roche 11836153001) was added to the tube at the ratio of 1 ml buffer per 100 mg tissue. The tissues were homogenized on a sonicator till no chunk of tissue visible, for about 30 sec. The extractions were kept in cold room for 1 hours with rotation. They were centrifuged at 10,000 g for 5 min at 4 C and the supernatants were carefully collected. Protein concentration was determined by Bradford assay (BioRad, 5000001) following manufacturer’s instructions. The final concentration of the samples was standardized to 0.5 mg/ml. samples were shipped to Olink Proteomics (Watertown, MA) on dry ice overnight. The Olink Immuno-oncology panel that analyses 96 immuno-oncology related human proteins were utilized. Normalized protein expression (NPX) values were generated and reported by Olink, and subsequently analyzed in-house using Prism (Graphpad) or Morpheus (Broad institute) online tool. PCA analysis and graph were performed with MATLAB (MathWorks, RRID:SCR_001622).

#### Hematoxylin and eosin staining to identify necrosis in GBM

Mice were sacrificed at humane endpoint with an overdose of ketamine and xylazine and perfused with 10 ml cold Ringer’s solution. The brain was carefully extracted and incubated in 10% formalin for 72 hours. The brains were dissected through the middle of the tumor and embedded in paraffin. The paraffin block was trimmed and the brains were sectioned on a microtome (Leica) to cut 5 µm sections. The sections were collected and mounted on a slide for automated hematoxylin and eosin staining as described above. The slides were scanned at 20x magnification with a whole-slide scanner (Hamamatsu). Tumor area in each section was determined in a blinded fashion in NDP.view2.

#### Enzyme-linked immunosorbent assay

Whole blood was collected from anesthetized mice via cardiac puncture. Blood Cell lysates for enzyme-linked immunosorbent assay (ELISA) were collected via sonication of cells in lysis buffer supplemented with protease and phosphatase inhibitors. ELISAs were performed for CCL2 (R&D, DY479), CCL7 (Boster Bio, EK0683), CCL8 (R&D, DY790), CCL11 (R&D, MME00), CCL12 (R&D, MCC120) and CCL5 (R&D, DY478-05) on cell lysates and cell supernatants according to the manufacturer’s instructions.

#### Flow Cytometry and spectral flow cytometry

Initial steps of the enzymatic dissociation of the tumors are the same as described above, except 0.5% collagenase D (Sigma, 11088858001) and DNase I (Roche, 11284932001) were used in place of papain. Single-cell suspensions were passed through 70 μm cell strainers, centrifuged, and resuspended in 30% Percoll (GE Healthcare, 17-0891-01) solution containing 10% FBS (Hyclone SH30396.03). Cells were separated by centrifugation at 800*g* for 15 minutes at 4°C. The supernatant was carefully removed to discard debris and lipids. The cells were then washed in cold PBS and resuspended in RBC lysis buffer (BioLegend, 420301) for 1 min at 37°C. Cells were transferred to an Eppendorf tube and washed once with FACS buffer (DPBS with 0.5% BSA) and blocked with 100 μl of 2x blocking solution (2% FBS, 5% normal rat serum, 5% normal mouse serum, 5% normal rabbit serum, 10 μg/ml anti-FcR (BioLegend, 101319) and 0.2% NaN_3_ in DPBS) on ice for 30 minutes. Cells were then stained with primary antibodies (Table S2) on ice for 30 minutes and washed with PBS. The cells were subsequently incubated in 100 μl viability dye (Zombie UV, BioLegend, 1:800) at room temperature for 20 min. The cells were washed and fixed with fixation buffer (eBioscience, 00-5123-43, 00-5223-56) for 30 min at 4 °C. Cells were washed and stained with the cocktail of antibodies examined myeloid lineage are set aside in the fridge until loading to the cytometer. Cells stained for the lymphoid panel were then permeabilized with a permeabilization buffer (eBioscience, 00-8333-56) before the intracellular markers were stained. The cells were washed and stored in fridge till analysis. Antibodies used in this study include are listed in table S2. All data were collected on a BD LSR II flow cytometer or Cytek Aurora spectral flow cytometer. Data were analyzed off line using FlowJo 10 software (Tree Star Inc., RRID:SCR_008520).

#### Single-cell RNA-seq and data analysis

Single cell suspensions of the tumors were obtained by papain dissociation as described above. Viability of single cells was assessed using Trypan Blue staining, and debris-free suspensions of >80% viability were deemed suitable for single-cell RNA Sequencing (scRNA-seq). Samples with lower viability were run with caution. Single cell RNA Seq was performed on these samples using the Chromium platform (10X Genomics) with the 3’ gene expression (3’ GEX) V3 kit, using an input of ∼10,000 cells. Briefly, Gel-Bead in Emulsions (GEMs) were generated on the sample chip in the Chromium controller. Barcoded cDNA was extracted from the GEMs by Post-GEM RT-cleanup and amplified for 12 cycles. Amplified cDNA was fragmented and subjected to end-repair, poly A-tailing, adapter ligation, and 10X-specific sample indexing following the manufacturer’s protocol. Libraries were quantified using Bioanalyzer (Agilent) and QuBit (ThermoFisher) analyses and were sequenced in paired end mode on a NovaSeq instrument (Illumina) targeting a depth of 50,000-100,000 reads per cell.

Raw fastq files were aligned to mouse genome reference mm10 customized to include the Rfp sequence, using CellRanger v5.0.0 (10X Genomics). Count matrices filtered by CellRanger algorithm were further filtered by discarding cells with either < 200 genes, < 1000 UMI (unique molecular identifier), or > 25% mitochondrial genes expressed. Data was processed and analyzed using R package Seurat v4.0.5. Normalization was performed using NormalizeData function with normalization.method = ‘LogNormalize’. Dimensionality reduction was computed on the top 2,000 variable features using FindVariableFeatures, ScaleData and RunPCA functions. UMAPs were generated using the top 15 PCs. For subclustering the immune compartment, we used R package Harmony to mitigate for batch effects driven by technical variation between replicates. *De novo* clustering using the Louvain algorithm was applied at different resolutions (0.2; 0.8; 2; 5; 8) on the SNN graph space. For high-level annotation, cell classes were identified in an iterative and semi-supervised fashion by assigning *de novo* discovered clusters to cell classes based on expression of known marker genes that define each cluster. Annotation of cell subtypes at a lower-level was performed in a similar manner as for the high-level and further aided by *de novo* marker discovery using the Seurat FindMarkers function and Wilcoxon Rank Sum test for differential expression analysis. To identify doublet-enriched clusters we looked for clusters of cells displaying expression of canonical markers for two or more different cell types and higher number of genes/UMI; such clusters were removed from further analysis.

Identification of modules of co-expressed genes was carried out using the R package scWGCNA (https://github.com/smorabit/scWGCNA) by first computing meta cells of 100 neighboring cells (k=100) using the function construct_metacells. To identify modules, function blockwiseConsensusModules was called with following parameters: softPower=12, deepSplit=3, mergeCutHeight = 0.25. Only the top 2,000 variable genes were used. Module scores, representing a normalized average expression of all genes in the WGCNA module, were computed using Seurat function AddModuleScore. Pathway enrichment analysis of gene modules identified using WGCNA was carried out using R package clusterProfiler (Yu et al., 2012).

#### Statistical analyses

Graphs were created using GraphPad Prism 9 (GraphPad Software Inc.) or R. Variables from two experimental groups were analyzed using unpaired or paired parametric two-tailed *t*-tests as appropriate, assuming equal standard deviations. One-way ANOVA was used to compare variables from more than two groups. Kaplan–Meier survival analysis was performed using the log-rank (Mantel-Cox) test and Gehan-Breslow-Wilcoxon test. Further details are included in the figure legends. Power analysis was performed based on prior experimental results obtained in the lab, with consideration of 10% attrition rate due to unexpected events such as spontaneous sarcoma, dermatitis or fight wound. (*) *P* < 0.05; (**) *P* < 0.01; (***) *P* < 0.001; (****) *P* < 0.0001; (ns) not significant. Final figures were assembled in Creative Cloud Photoshop (Adobe, RRID:SCR_014199).

## ACKNOWLEDGEMENTS

We would like to acknowledge the Mouse Transgenic and Gene Targeting Core, Flow Cytometry Core and the Integrated Cellular Imaging Cores at Emory University for their services. We would also like to acknowledge the Mount Sinai Dean’s Flow Cytometry Core and Genomics Core for scRNA-seq services. The Tisch Cancer Institute and related research facilities are supported by P30 CA196521. We extend our thanks to Mr. David R. Schumick for generating illustrations and Dr. Christopher Nelson for scientific editing. This work was supported by NIH/NINDS R01 NS100864 and start-up funds to DH from Departments of Oncological Sciences and Neurosurgery, Icahn School of Medicine, Mount Sinai. A. Lujambio was supported by Damon Runyon-Rachleff Innovation Award (DR52-18) and R37 Merit Award (R37CA230636), and Icahn School of Medicine at Mount Sinai. E.E.B. Ghosn was supported by NIH/NIAID R01AI123126.

## AUTHOR CONTRIBUTIONS

### Concept and design

Z. Chen, D. Hambardzumyan

### Development of Methodology

Z. Chen, A. Lujambio, E.E.B, Gohsn, A.M. Tsankov, D. Hambardzumyan

### Acquisition of data (provided animals, acquired and managed patients, provided facilities, etc.)

Z. Chen, G. Pinero, D.J. Eddins, K.E. Lindblad, J.L. Ross, N. Tsankova, D.H. Gutmann, E.E.B. Ghosn, S.A. Lira, D. Hambardzumyan

### Analysis and interpretation of data (e.g., statistical analysis, biostatistics, computational analysis)

Z. Chen, N. Soni, G. Pinero, B. Giotti, D.J. Eddins, J.L. Ross, E.E.B. Ghosn, A.M. Tsankov, D Hambardzumyan

### Writing, review, and/or revision of the manuscript

Z. Chen and D. Hambardzumyan created the original draft. All authors participated in review and editing.

### Administrative, technical, or material support (i.e. reporting or organizing data, constructing databases)

Z. Chen, N. Soni, B. Giotti

### Study supervision

Z. Chen, D. Hambardzumyan

## DECLARATION OF INTERESTS

The authors have no relevant competing interests to disclose.

### Data and materials availability

The data that support the findings of this study are available from the corresponding author upon reasonable request. ScRNA-Seq data were deposited at GEO with accession number GSE203154. Newly created qMCP-KO mice will be distributed to interested colleagues upon mutually satisfactory materials transfer agreements.

## REFERENCES

Akkari, L., Bowman, R.L., Tessier, J., Klemm, F., Handgraaf, S.M., de Groot, M., Quail, D.F., Tillard, L., Gadiot, J., Huse, J.T., et al. (2020). Dynamic changes in glioma macrophage populations after radiotherapy reveal CSF-1R inhibition as a strategy to overcome resistance. Sci Transl Med 12.

Aranda, P.S., LaJoie, D.M., and Jorcyk, C.L. (2012). Bleach gel: a simple agarose gel for analyzing RNA quality. Electrophoresis 33, 366–369.

Becher, O.J., Hambardzumyan, D., Fomchenko, E.I., Momota, H., Mainwaring, L., Bleau, A.M., Katz, A.M., Edgar, M., Kenney, A.M., Cordon-Cardo, C., et al. (2008). Gli activity correlates with tumor grade in platelet-derived growth factor-induced gliomas. Cancer research 68, 2241–2249.

Becher, O.J., Hambardzumyan, D., Walker, T.R., Helmy, K., Nazarian, J., Albrecht, S., Hiner, R.L., Gall, S., Huse, J.T., Jabado, N., et al. (2010). Preclinical evaluation of radiation and perifosine in a genetically and histologically accurate model of brainstem glioma. Cancer research 70, 2548–2557.

Belperio, J.A., Keane, M.P., Burdick, M.D., Londhe, V., Xue, Y.Y., Li, K., Phillips, R.J., and Strieter, R.M. (2002). Critical role for CXCR2 and CXCR2 ligands during the pathogenesis of ventilator-induced lung injury. J Clin Invest 110, 1703–1716.

Boivin, G., Faget, J., Ancey, P.B., Gkasti, A., Mussard, J., Engblom, C., Pfirschke, C., Contat, C., Pascual, J., Vazquez, J., et al. (2020). Durable and controlled depletion of neutrophils in mice. Nat Commun 11, 2762.

Bollard, J., Miguela, V., Ruiz de Galarreta, M., Venkatesh, A., Bian, C.B., Roberto, M.P., Tovar, V., Sia, D., Molina-Sanchez, P., Nguyen, C.B., et al. (2016). Palbociclib (PD-0332991), a selective CDK4/6 inhibitor, restricts tumour growth in preclinical models of hepatocellular carcinoma. Gut.

Brennan, C.W., Verhaak, R.G., McKenna, A., Campos, B., Noushmehr, H., Salama, S.R., Zheng, S., Chakravarty, D., Sanborn, J.Z., Berman, S.H., et al. (2013). The somatic genomic landscape of glioblastoma. Cell 155, 462–477.

Buonfiglioli, A., and Hambardzumyan, D. (2021). Macrophages and microglia: the cerberus of glioblastoma. Acta Neuropathol Commun 9, 54.

Butowski, N., Colman, H., De Groot, J.F., Omuro, A.M., Nayak, L., Wen, P.Y., Cloughesy, T.F., Marimuthu, A., Haidar, S., Perry, A., et al. (2016). Orally administered colony stimulating factor 1 receptor inhibitor PLX3397 in recurrent glioblastoma: an Ivy Foundation Early Phase Clinical Trials Consortium phase II study. Neuro-oncology 18, 557–564.

Cancer Genome Atlas Research Network. Electronic address, w.b.e., and Cancer Genome Atlas Research, N. (2017). Comprehensive and Integrative Genomic Characterization of Hepatocellular Carcinoma. Cell 169, 1327–1341 e1323.

Cao, Q., Li, B., Wang, X., Sun, K., and Guo, Y. (2018). Therapeutic inhibition of CXC chemokine receptor 2 by SB225002 attenuates LPS-induced acute lung injury in mice. Arch Med Sci 14, 635–644.

Chandrasekharan, U.M., Siemionow, M., Unsal, M., Yang, L., Poptic, E., Bohn, J., Ozer, K., Zhou, Z., Howe, P.H., Penn, M., et al. (2007). Tumor necrosis factor alpha (TNF-alpha) receptor-II is required for TNF-alpha-induced leukocyte-endothelial interaction in vivo. Blood 109, 1938–1944.

Chen, X., and Calvisi, D.F. (2014). Hydrodynamic transfection for generation of novel mouse models for liver cancer research. Am J Pathol 184, 912–923.

Chen, Z., Feng, X., Herting, C.J., Garcia, V.A., Nie, K., Pong, W.W., Rasmussen, R., Dwivedi, B., Seby, S., Wolf, S.A., et al. (2017). Cellular and Molecular Identity of Tumor-Associated Macrophages in Glioblastoma. Cancer research 77, 2266–2278.

Chen, Z., Herting, C.J., Ross, J.L., Gabanic, B., Puigdelloses Vallcorba, M., Szulzewsky, F., Wojciechowicz, M.L., Cimino, P.J., Ezhilarasan, R., Sulman, E.P., et al. (2020). Genetic driver mutations introduced in identical cell-of-origin in murine glioblastoma reveal distinct immune landscapes but similar response to checkpoint blockade. Glia 68, 2148–2166.

Coniglio, S.J., Eugenin, E., Dobrenis, K., Stanley, E.R., West, B.L., Symons, M.H., and Segall, J.E. (2012). Microglial stimulation of glioblastoma invasion involves epidermal growth factor receptor (EGFR) and colony stimulating factor 1 receptor (CSF-1R) signaling. Mol Med 18, 519–527.

Efremova, M., Vento-Tormo, M., Teichmann, S.A., and Vento-Tormo, R. (2020). CellPhoneDB: inferring cell-cell communication from combined expression of multi-subunit ligand-receptor complexes. Nat Protoc 15, 1484–1506.

Fedele, M., Cerchia, L., Pegoraro, S., Sgarra, R., and Manfioletti, G. (2019). Proneural-Mesenchymal Transition: Phenotypic Plasticity to Acquire Multitherapy Resistance in Glioblastoma. Int J Mol Sci 20. Franklin, K.B.J., and Paxinos, G. (1997). The mouse brain in stereotaxic coordinates (San Diego: Academic Press).

Fuentes, M.E., Durham, S.K., Swerdel, M.R., Lewin, A.C., Barton, D.S., Megill, J.R., Bravo, R., and Lira, S.A. (1995). Controlled recruitment of monocytes and macrophages to specific organs through transgenic expression of monocyte chemoattractant protein-1. Journal of immunology 155, 5769–5776.

Gabrusiewicz, K., Rodriguez, B., Wei, J., Hashimoto, Y., Healy, L.M., Maiti, S.N., Thomas, G., Zhou, S., Wang, Q., Elakkad, A., et al. (2016). Glioblastoma-infiltrated innate immune cells resemble M0 macrophage phenotype. JCI Insight 1.

Girbl, T., Lenn, T., Perez, L., Rolas, L., Barkaway, A., Thiriot, A., Del Fresno, C., Lynam, E., Hub, E., Thelen, M., et al. (2018). Distinct Compartmentalization of the Chemokines CXCL1 and CXCL2 and the Atypical Receptor ACKR1 Determine Discrete Stages of Neutrophil Diapedesis. Immunity 49, 1062–1076 e1066.

Halliday, J., Helmy, K., Pattwell, S.S., Pitter, K.L., LaPlant, Q., Ozawa, T., and Holland, E.C. (2014). In vivo radiation response of proneural glioma characterized by protective p53 transcriptional program and proneural-mesenchymal shift. Proc Natl Acad Sci U S A 111, 5248–5253.

Hara, T., Chanoch-Myers, R., Mathewson, N.D., Myskiw, C., Atta, L., Bussema, L., Eichhorn, S.W., Greenwald, A.C., Kinker, G.S., Rodman, C., et al. (2021). Interactions between cancer cells and immune cells drive transitions to mesenchymal-like states in glioblastoma. Cancer Cell 39, 779–792 e711.

Herting, C.J., Chen, Z., Pitter, K.L., Szulzewsky, F., Kaffes, I., Kaluzova, M., Park, J.C., Cimino, P.J., Brennan, C., Wang, B., et al. (2017). Genetic driver mutations define the expression signature and microenvironmental composition of high-grade gliomas. Glia 65, 1914–1926.

Jones, C., Karajannis, M.A., Jones, D.T.W., Kieran, M.W., Monje, M., Baker, S.J., Becher, O.J., Cho, Y.J., Gupta, N., Hawkins, C., et al. (2017). Pediatric high-grade glioma: biologically and clinically in need of new thinking. Neuro Oncol 19, 153–161.

Kaffes, I., Szulzewsky, F., Chen, Z., Herting, C.J., Gabanic, B., Velazquez Vega, J.E., Shelton, J., Switchenko, J.M., Ross, J.L., McSwain, L.F., et al. (2019). Human Mesenchymal glioblastomas are characterized by an increased immune cell presence compared to Proneural and Classical tumors. Oncoimmunology 8, e1655360.

Kastenhuber, E.R., Huse, J.T., Berman, S.H., Pedraza, A., Zhang, J., Suehara, Y., Viale, A., Cavatore, M., Heguy, A., Szerlip, N., et al. (2014). Quantitative assessment of intragenic receptor tyrosine kinase deletions in primary glioblastomas: their prevalence and molecular correlates. Acta Neuropathol 127, 747–759.

Kuang, D.M., Zhao, Q., Wu, Y., Peng, C., Wang, J., Xu, Z., Yin, X.Y., and Zheng, L. (2011). Peritumoral neutrophils link inflammatory response to disease progression by fostering angiogenesis in hepatocellular carcinoma. J Hepatol 54, 948–955.

Kumar, V., Donthireddy, L., Marvel, D., Condamine, T., Wang, F., Lavilla-Alonso, S., Hashimoto, A., Vonteddu, P., Behera, R., Goins, M.A., et al. (2017). Cancer-Associated Fibroblasts Neutralize the Anti-tumor Effect of CSF1 Receptor Blockade by Inducing PMN-MDSC Infiltration of Tumors. Cancer Cell 32, 654–668 e655.

Langfelder, P., and Horvath, S. (2008). WGCNA: an R package for weighted correlation network analysis. BMC Bioinformatics 9, 559.

Liang, J., Piao, Y., Holmes, L., Fuller, G.N., Henry, V., Tiao, N., and De Groot, J.F. (2014). Neutrophils promote the malignant glioma phenotype through S100A4. Clinical Cancer Research 20, 187–198.

Lim, S.Y., Yuzhalin, A.E., Gordon-Weeks, A.N., and Muschel, R.J. (2016). Targeting the CCL2-CCR2 signaling axis in cancer metastasis. Oncotarget.

Lin, Y.J., Wei, K.C., Chen, P.Y., Lim, M., and Hwang, T.L. (2021). Roles of Neutrophils in Glioma and Brain Metastases. Front Immunol 12, 701383.

Louis, D.N., Perry, A., Reifenberger, G., von Deimling, A., Figarella-Branger, D., Cavenee, W.K., Ohgaki, H., Wiestler, O.D., Kleihues, P., and Ellison, D.W. (2016). The 2016 World Health Organization Classification of Tumors of the Central Nervous System: a summary. Acta Neuropathol 131, 803–820.

Magod, P., Mastandrea, I., Rousso-Noori, L., Agemy, L., Shapira, G., Shomron, N., and Friedmann-Morvinski, D. (2021). Exploring the longitudinal glioma microenvironment landscape uncovers reprogrammed pro-tumorigenic neutrophils in the bone marrow. Cell reports 36, 109480.

Margetts, J., Ogle, L.F., Chan, S.L., Chan, A.W.H., Chan, K.C.A., Jamieson, D., Willoughby, C.E., Mann, D.A., Wilson, C.L., Manas, D.M., et al. (2018). Neutrophils: driving progression and poor prognosis in hepatocellular carcinoma? British journal of cancer 118, 248–257.

Maximov, V., Chen, Z., Wei, Y., Robinson, M.H., Herting, C.J., Shanmugam, N.S., Rudneva, V.A., Goldsmith, K.C., MacDonald, T.J., Northcott, P.A., et al. (2019). Tumour-associated macrophages exhibit anti-tumoural properties in Sonic Hedgehog medulloblastoma. Nature communications 10, 2410–2410.

McLendon, R., Friedman, A., Bigner, D., Van Meir, E.G., Brat, D.J., Mastrogianakis, G.M., Olson, J.J., Mikkelsen, T., Lehman, N., Aldape, K., et al. (2008). Comprehensive genomic characterization defines human glioblastoma genes and core pathways. Nature 455, 1061–1068.

Mei, J., Liu, Y., Dai, N., Hoffmann, C., Hudock, K.M., Zhang, P., Guttentag, S.H., Kolls, J.K., Oliver, P.M., Bushman, F.D., et al. (2012). Cxcr2 and Cxcl5 regulate the IL-17/G-CSF axis and neutrophil homeostasis in mice. J Clin Invest 122, 974–986.

Muller, S., Kohanbash, G., Liu, S.J., Alvarado, B., Carrera, D., Bhaduri, A., Watchmaker, P.B., Yagnik, G., Di Lullo, E., Malatesta, M., et al. (2017). Single-cell profiling of human gliomas reveals macrophage ontogeny as a basis for regional differences in macrophage activation in the tumor microenvironment. Genome Biol 18, 234.

Neftel, C., Laffy, J., Filbin, M.G., Hara, T., Shore, M.E., Rahme, G.J., Richman, A.R., Silverbush, D., Shaw, M.L., Hebert, C.M., et al. (2019). An Integrative Model of Cellular States, Plasticity, and Genetics for Glioblastoma. Cell 178, 835–849 e821.

Nolan, E., Bridgeman, V.L., Ombrato, L., Karoutas, A., Rabas, N., Sewnath, C.A.N., Vasquez, M., Rodrigues, F.S., Horswell, S., Faull, P., et al. (2022). Radiation exposure elicits a neutrophil-driven response in healthy lung tissue that enhances metastatic colonization. Nat Cancer 3, 173–187.

O’Brien, S.A., Orf, J., Skrzypczynska, K.M., Tan, H., Kim, J., DeVoss, J., Belmontes, B., and Egen, J.G. (2021). Activity of tumor-associated macrophage depletion by CSF1R blockade is highly dependent on the tumor model and timing of treatment. Cancer Immunol Immunother 70, 2401–2410.

Olivier, M., Hollstein, M., and Hainaut, P. (2010). TP53 mutations in human cancers: origins, consequences, and clinical use. Cold Spring Harb Perspect Biol 2, a001008.

Omuro, A., Beal, K., Gutin, P., Karimi, S., Correa, D.D., Kaley, T.J., DeAngelis, L.M., Chan, T.A., Gavrilovic, I.T., Nolan, C., et al. (2014). Phase II study of bevacizumab, temozolomide, and hypofractionated stereotactic radiotherapy for newly diagnosed glioblastoma. Clin Cancer Res 20, 5023–5031.

Patel, A.P., Tirosh, I., Trombetta, J.J., Shalek, A.K., Gillespie, S.M., Wakimoto, H., Cahill, D.P., Nahed, B.V., Curry, W.T., Martuza, R.L., et al. (2014). Single-cell RNA-seq highlights intratumoral heterogeneity in primary glioblastoma. Science (New York, NY) 344, 1396–1401.

Pitter, K.L., Tamagno, I., Alikhanyan, K., Hosni-Ahmed, A., Pattwell, S.S., Donnola, S., Dai, C., Ozawa, T., Chang, M., Chan, T.A., et al. (2016). Corticosteroids compromise survival in glioblastoma. Brain 139, 1458–1471.

Proudfoot, A.E. (2002). Chemokine receptors: multifaceted therapeutic targets. Nature reviews Immunology 2, 106–115.

Pyonteck, S.M., Akkari, L., Schuhmacher, A.J., Bowman, R.L., Sevenich, L., Quail, D.F., Olson, O.C., Quick, M.L., Huse, J.T., Teijeiro, V., et al. (2013). CSF-1R inhibition alters macrophage polarization and blocks glioma progression. Nat Med 19, 1264–1272.

Quail, D.F., Amulic, B., Aziz, M., Barnes, B.J., Eruslanov, E., Fridlender, Z.G., Goodridge, H.S., Granot, Z., Hidalgo, A., Huttenlocher, A., et al. (2022). Neutrophil phenotypes and functions in cancer: A consensus statement. The Journal of experimental medicine 219.

Rahbar, A., Cederarv, M., Wolmer-Solberg, N., Tammik, C., Stragliotto, G., Peredo, I., Fornara, O., Xu, X., Dzabic, M., Taher, C., et al. (2016). Enhanced neutrophil activity is associated with shorter time to tumor progression in glioblastoma patients. OncoImmunology 5.

Ruiz de Galarreta, M., Bresnahan, E., Molina-Sanchez, P., Lindblad, K.E., Maier, B., Sia, D., Puigvehi, M., Miguela, V., Casanova-Acebes, M., Dhainaut, M., et al. (2019). beta-Catenin Activation Promotes Immune Escape and Resistance to Anti-PD-1 Therapy in Hepatocellular Carcinoma. Cancer discovery 9, 1124–1141.

Sawada, M., Kiyono, T., Nakashima, S., Shinoda, J., Naganawa, T., Hara, S., Iwama, T., and Sakai, N. (2004). Molecular mechanisms of TNF-alpha-induced ceramide formation in human glioma cells: P53-mediated oxidant stress-dependent and -independent pathways. Cell Death Differ 11, 997–1008.

Shea-Donohue, T., Thomas, K., Cody, M.J., Aiping, Z., Detolla, L.J., Kopydlowski, K.M., Fukata, M., Lira, S.A., and Vogel, S.N. (2008). Mice deficient in the CXCR2 ligand, CXCL1 (KC/GRO-alpha), exhibit increased susceptibility to dextran sodium sulfate (DSS)-induced colitis. Innate Immun 14, 117–124.

Sierro, F., Evrard, M., Rizzetto, S., Melino, M., Mitchell, A.J., Florido, M., Beattie, L., Walters, S.B., Tay, S.S., Lu, B., et al. (2017). A Liver Capsular Network of Monocyte-Derived Macrophages Restricts Hepatic Dissemination of Intraperitoneal Bacteria by Neutrophil Recruitment. Immunity 47, 374–388 e376.

Sottoriva, A., Spiteri, I., Piccirillo, S.G.M., Touloumis, A., Collins, V.P., Marioni, J.C., Curtis, C., Watts, C., and Tavare, S. (2013). Intratumor heterogeneity in human glioblastoma reflects cancer evolutionary dynamics. Proceedings of the National Academy of Sciences of the United States of America 110, 4009–4014.

Tan, I.L., Arifa, R.D.N., Rallapalli, H., Kana, V., Lao, Z., Sanghrajka, R.M., Sumru Bayin, N., Tanne, A., Wojcinski, A., Korshunov, A., et al. (2021). CSF1R inhibition depletes tumor-associated macrophages and attenuates tumor progression in a mouse sonic Hedgehog-Medulloblastoma model. Oncogene 40, 396–407.

Tani, M., Fuentes, M.E., Peterson, J.W., Trapp, B.D., Durham, S.K., Loy, J.K., Bravo, R., Ransohoff, R.M., and Lira, S.A. (1996). Neutrophil infiltration, glial reaction, and neurological disease in transgenic mice expressing the chemokine N51/KC in oligodendrocytes. J Clin Invest 98, 529–539.

Verhaak, R.G., Hoadley, K.A., Purdom, E., Wang, V., Qi, Y., Wilkerson, M.D., Miller, C.R., Ding, L., Golub, T., Mesirov, J.P., et al. (2010). Integrated genomic analysis identifies clinically relevant subtypes of glioblastoma characterized by abnormalities in PDGFRA, IDH1, EGFR, and NF1. Cancer Cell 17, 98–110.

Wang, Q., Hu, B., Hu, X., Kim, H., Squatrito, M., Scarpace, L., deCarvalho, A.C., Lyu, S., Li, P., Li, Y., et al. (2018). Tumor Evolution of Glioma-Intrinsic Gene Expression Subtypes Associates with Immunological Changes in the Microenvironment. Cancer Cell 33, 152.

Wang, Q., Hu, B., Hu, X., Kim, H., Squatrito, M., Scarpace, L., deCarvalho, A.C., Lyu, S., Li, P., Li, Y., et al. (2017). Tumor Evolution of Glioma-Intrinsic Gene Expression Subtypes Associates with Immunological Changes in the Microenvironment. Cancer Cell 32, 42–56.e46.

White, J.R., Lee, J.M., Young, P.R., Hertzberg, R.P., Jurewicz, A.J., Chaikin, M.A., Widdowson, K., Foley, J.J., Martin, L.D., Griswold, D.E., et al. (1998). Identification of a potent, selective non-peptide CXCR2 antagonist that inhibits interleukin-8-induced neutrophil migration. J Biol Chem 273, 10095–10098.

Yee, P.P., Wei, Y., Kim, S.Y., Lu, T., Chih, S.Y., Lawson, C., Tang, M., Liu, Z., Anderson, B., Thamburaj, K., et al. (2020). Neutrophil-induced ferroptosis promotes tumor necrosis in glioblastoma progression. Nat Commun 11, 5424.

Yellowhair, T.R., Newville, J.C., Noor, S., Maxwell, J.R., Milligan, E.D., Robinson, S., and Jantzie, L.L. (2019). CXCR2 Blockade Mitigates Neural Cell Injury Following Preclinical Chorioamnionitis. Front Physiol 10, 324.

Yu, G., Wang, L.G., Han, Y., and He, Q.Y. (2012). clusterProfiler: an R package for comparing biological themes among gene clusters. OMICS 16, 284–287.

