## Supplemental materials for "The novel compensatory reciprocal interplay between neutrophils and monocytes drives cancer progression"

**Supplementary Materials**

**Supplementary Figures**

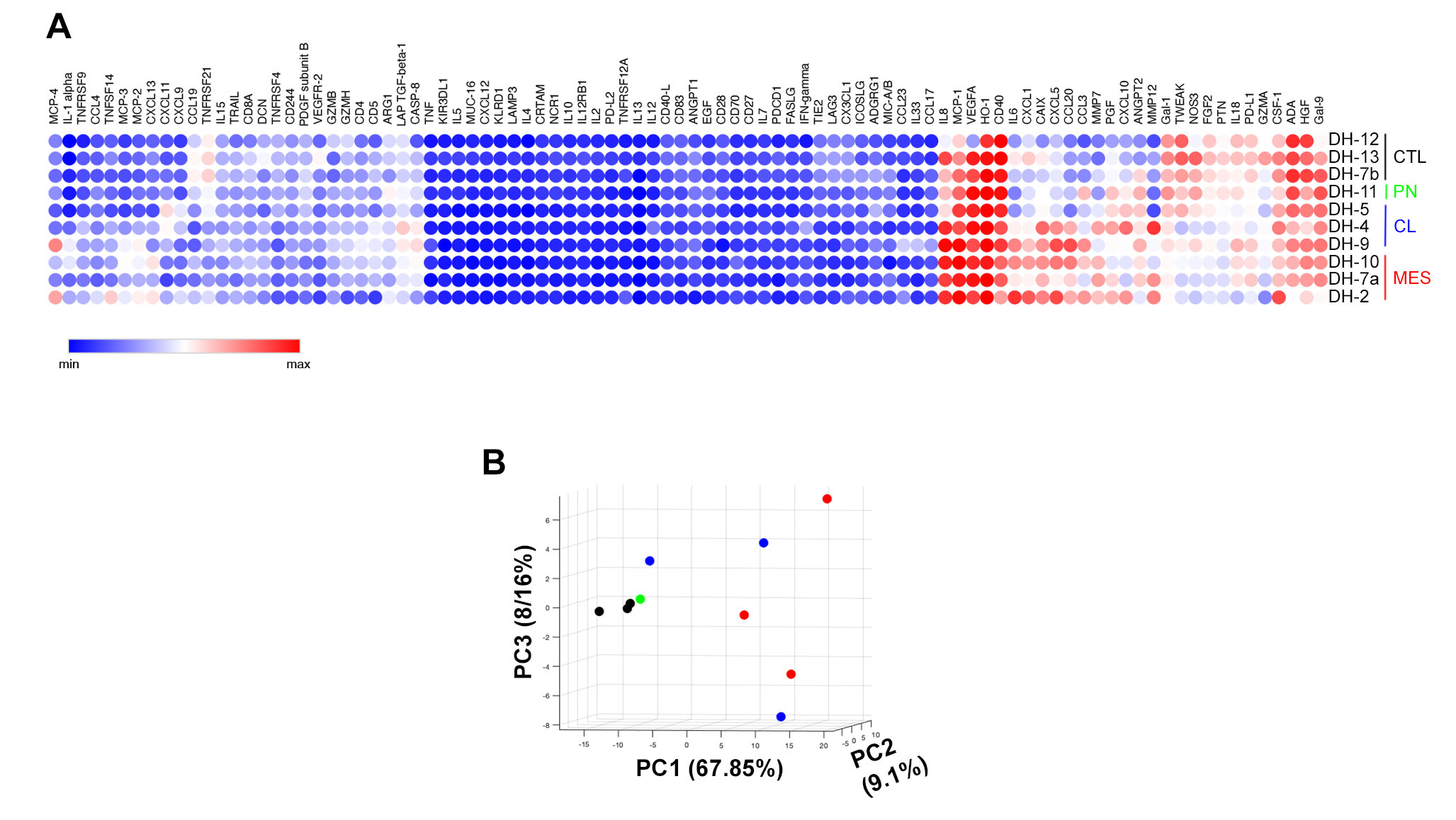

**Figure S1. Illustration of Olink multiplex proteomics with 96 immune protein targets in human control brain and GBM subsets. (A)** Heatmap showing normalized expression score of the analytes. (**B**) Principal component analysis of the results. CTL=control, PN = proneural, CL = classical, MES = mesenchymal.

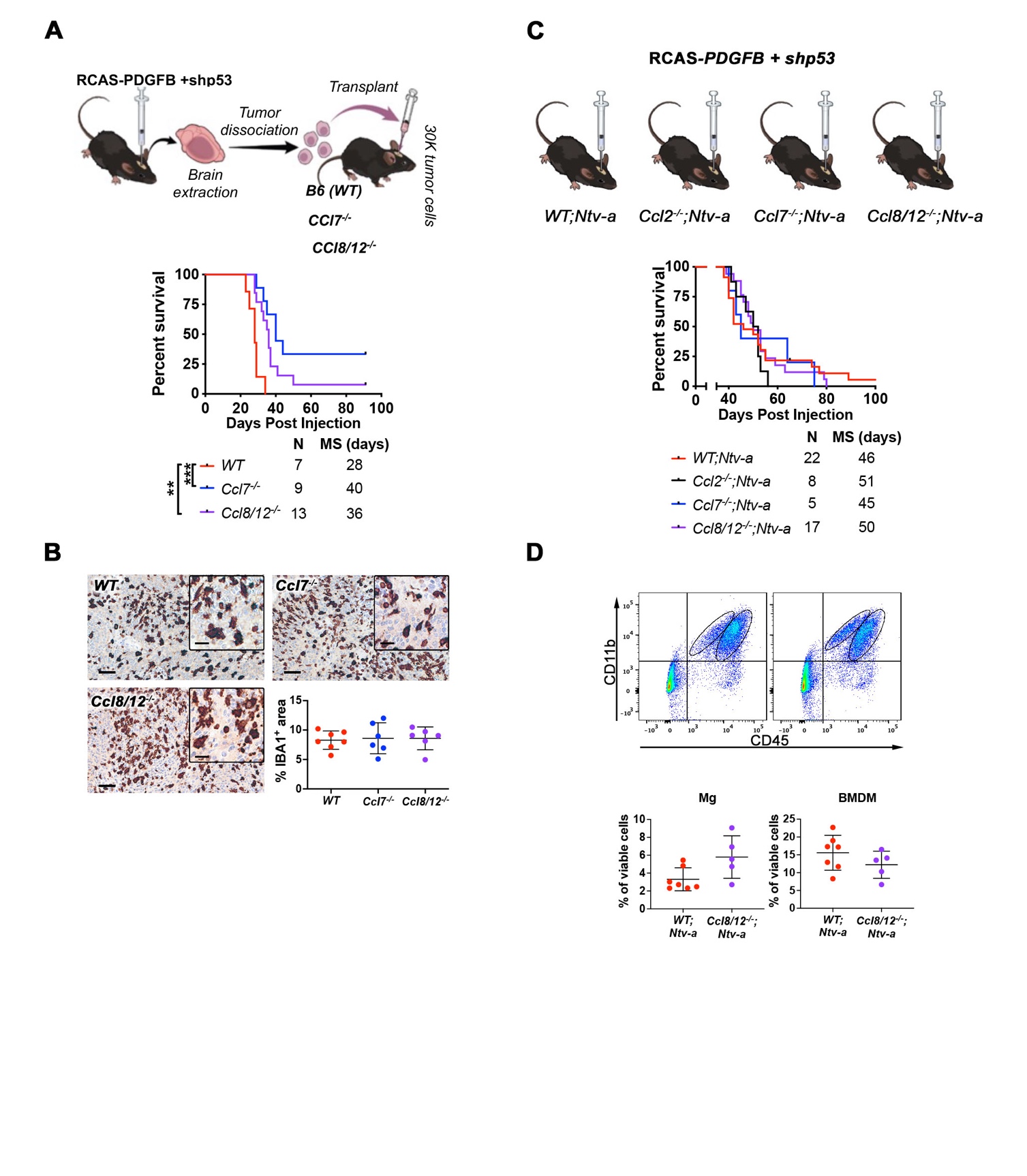

**Figure S2. Loss of *Ccl7* or *Ccl8/12* in TME but not tumor cells results in extended survival of GBM-bearing mice. (A)** Schematic illustration of orthotopic transplant of primary *PDGFB*-driven WT tumors into *Ccl7^-/-^* and *Ccl8/12^-/-^* (CCL7 or CCL8/12 are lost only in TME), and WT recipient animals and their corresponding Kaplan-Meier survival curves. Log-rank test. **(B)** Representative images and quantification of immunohistochemical staining of IBA1. **(C)** Schematic illustration of injections and Kaplan Meier-survival curves of *PDGFB*-driven tumors generated in *WT;Ntv-a*, *Ccl2^-/-^;Ntv-a, Ccl7^-/-^;Ntv-a* and *Ccl8/12^-/-^;Ntv-a* mice. **(D)** Dot plots of flow cytometry analysis of BMDM and Mg in tumors at humane endpoint. Student’s *t*-test was applied. **p<0.01, ***p<0.001. Scale bar = 50 μm, scale bar in inset = 20 μm.

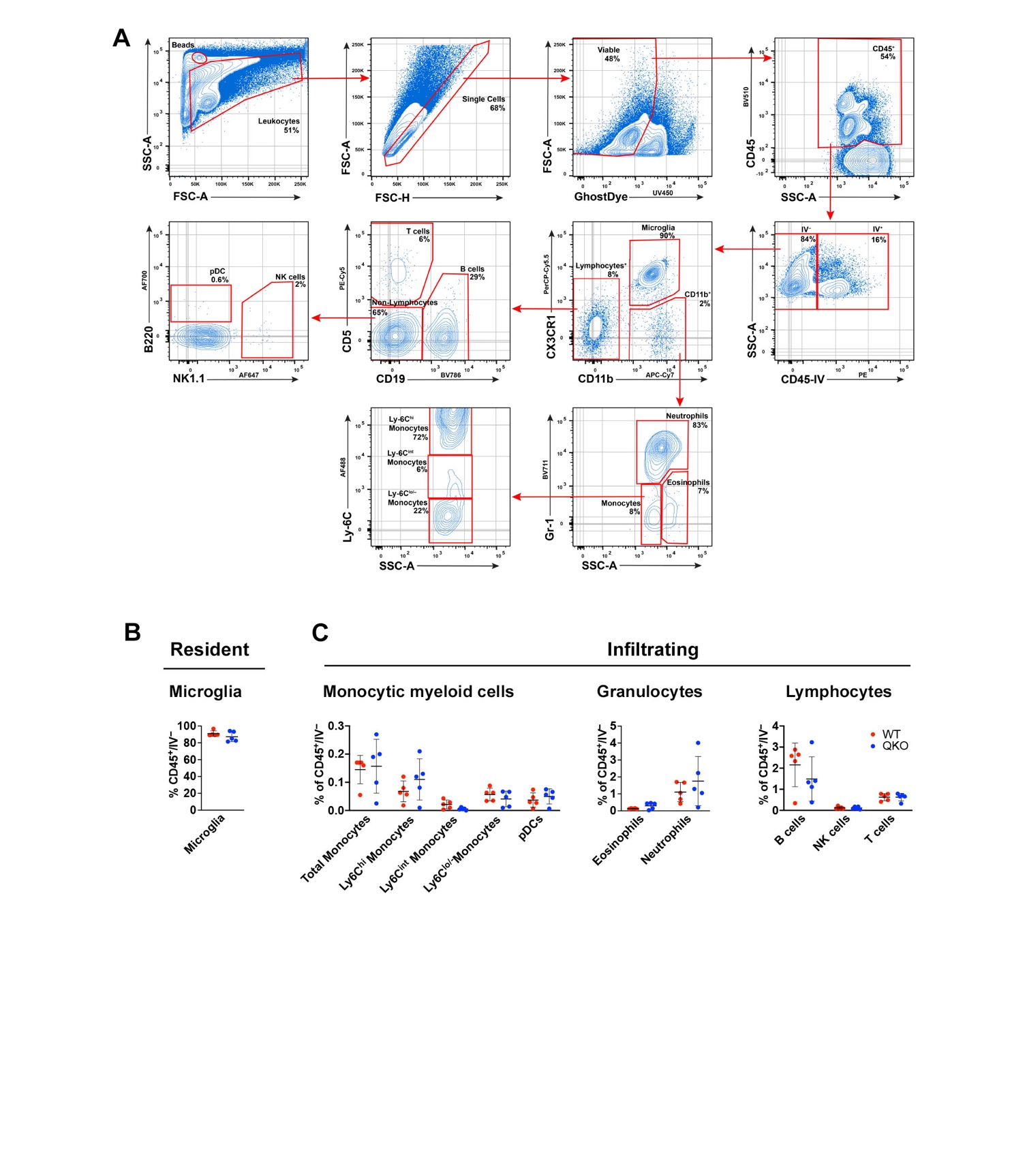

**Figure S3. Flow cytometric analysis of brain infiltrating immune cells in healthy adult mice. (A)** Gating strategy identifying subsets of immune cells. (**B**) Quantification of resident immune cells. (**C**) Quantification of brain infiltrating immune cells.

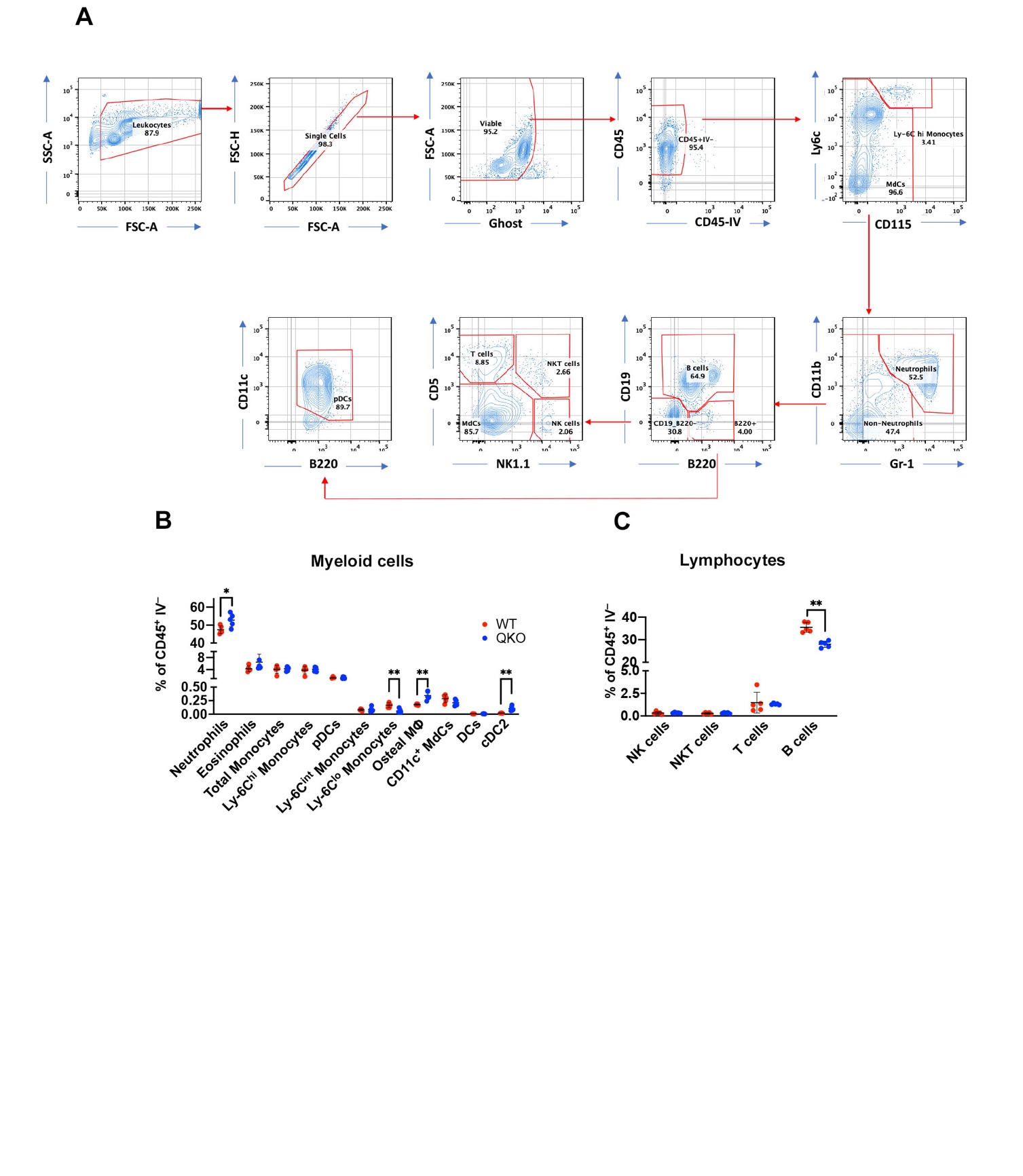

**Figure S4. Flow cytometric analysis of immune cells in the bone marrow of healthy adult mice. (A)** Gating strategy identifying subsets of immune cells. (**B**) Quantification of myeloid cells. (**C**) Quantification of lymphoid cells. Student’s *t*-test. *p<0.05, **p<0.01.

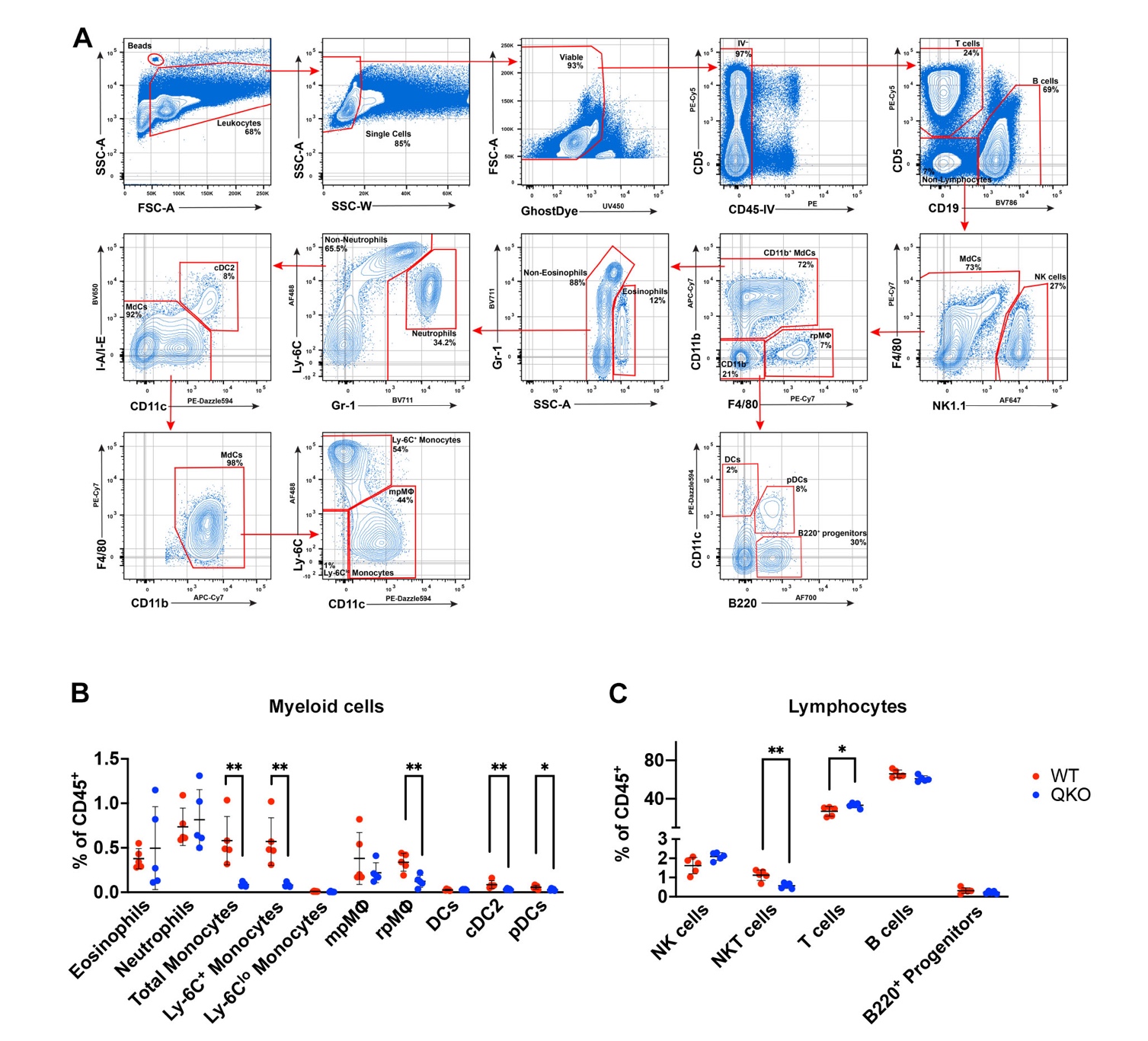

**Figure S5. Flow cytometric analysis of immune cells in spleen of healthy adult mice. (A)** Gating strategy identifying subsets of immune cells. (**B**) Quantification of myeloid cells. (**C**) Quantification of lymphoid cells. Student’s *t*-test. *p<0.05, **p<0.01.

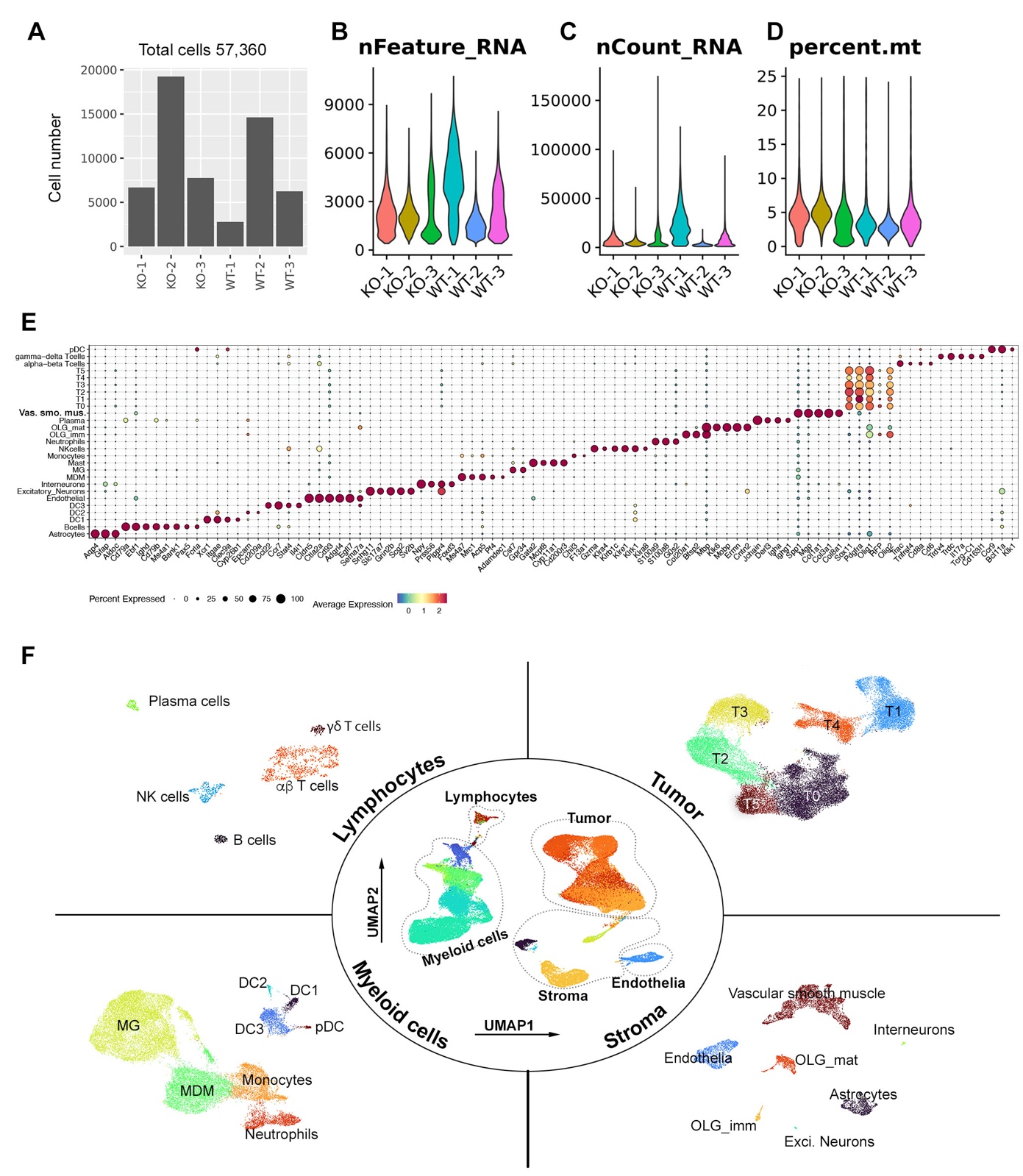

**Figure S6. Single-cell RNA seq analysis of tumors generated in *WT;Ntv-a* and *qMCP^-/-^;Ntv-a* mice**. (**A**) Total number of cells per samples after removing doublets. (**B**) Distribution of the number of genes (features) detected in each cell per sample. (**C**) Distribution of number of unique molecular identifiers (UMIs) in each cell per sample. (**D**) Distribution of percentage of mitochondrial genes expressed in each cell per sample. (**E**) Expression of select marker genes (x axis) for each annotated cell subset (y axis) colored by normalized expression level, where the size of dots represents the percentage of cells expressed. (**F**) UMAP dimensionality reductions of the stratified cells. OLG: oligodendrocytes, imm: immature, mat: mature, Exci: excitatory.

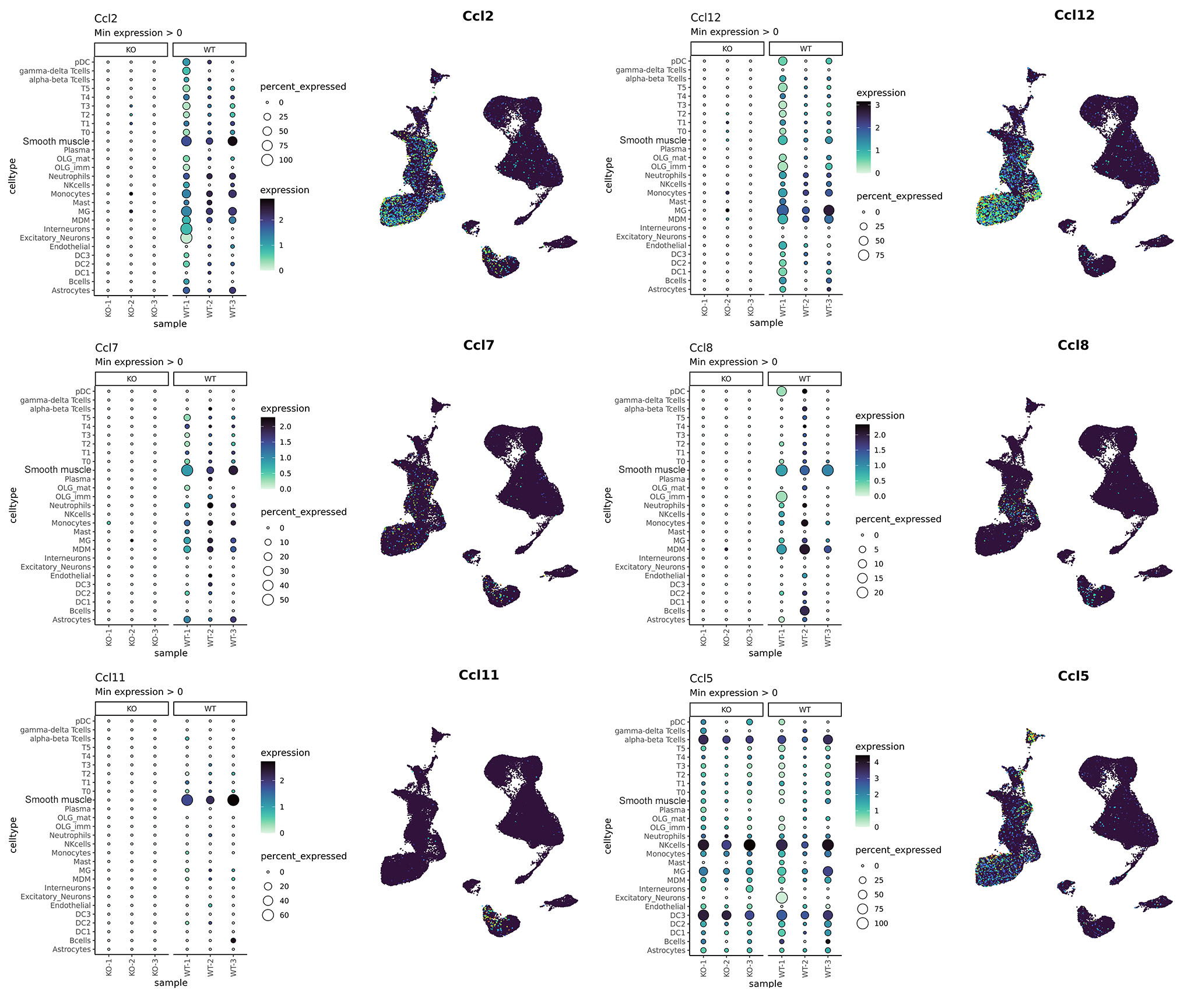

**Figure S7. Expression of MCP-related genes across cell types and samples.** Left in each panel: dot plots showing percentage of expressed cells (dot size) and normalized expression level (color gradient) across cell types (y-axes) and mouse tumor samples (x-axes). Right in each panel: UMAPs showing the expression of MCPs across all cells using the same UMAP embedding as in Figure 3B.

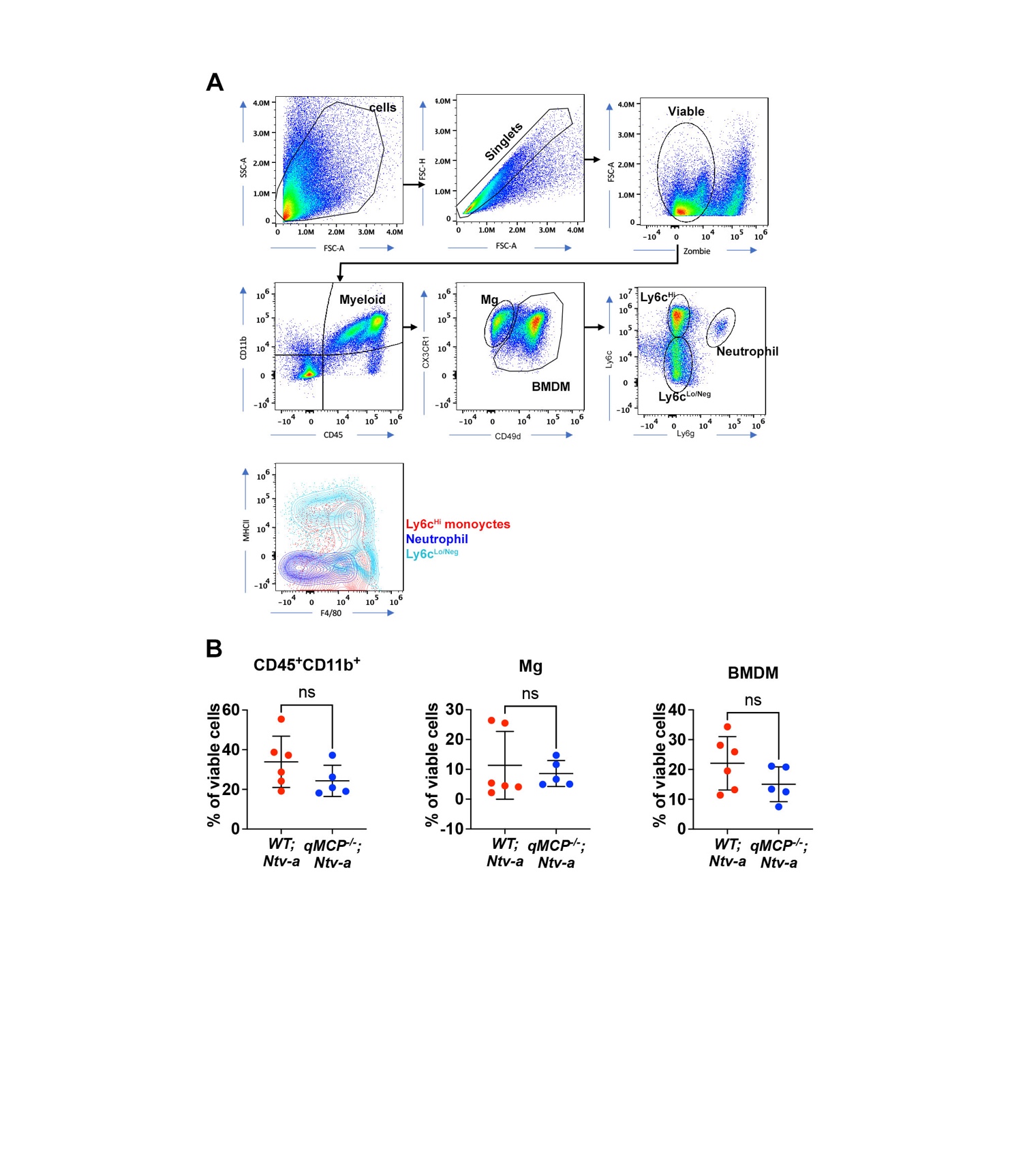

**Figure S8. Spectral flow cytometry analysis of myeloid cells in GBM. (A)** Gating strategy identifying myeloid cells in GBM. (**B**) Dot plots showing subsets of myeloid cells in the tumors. Student’s *t*-test was applied.

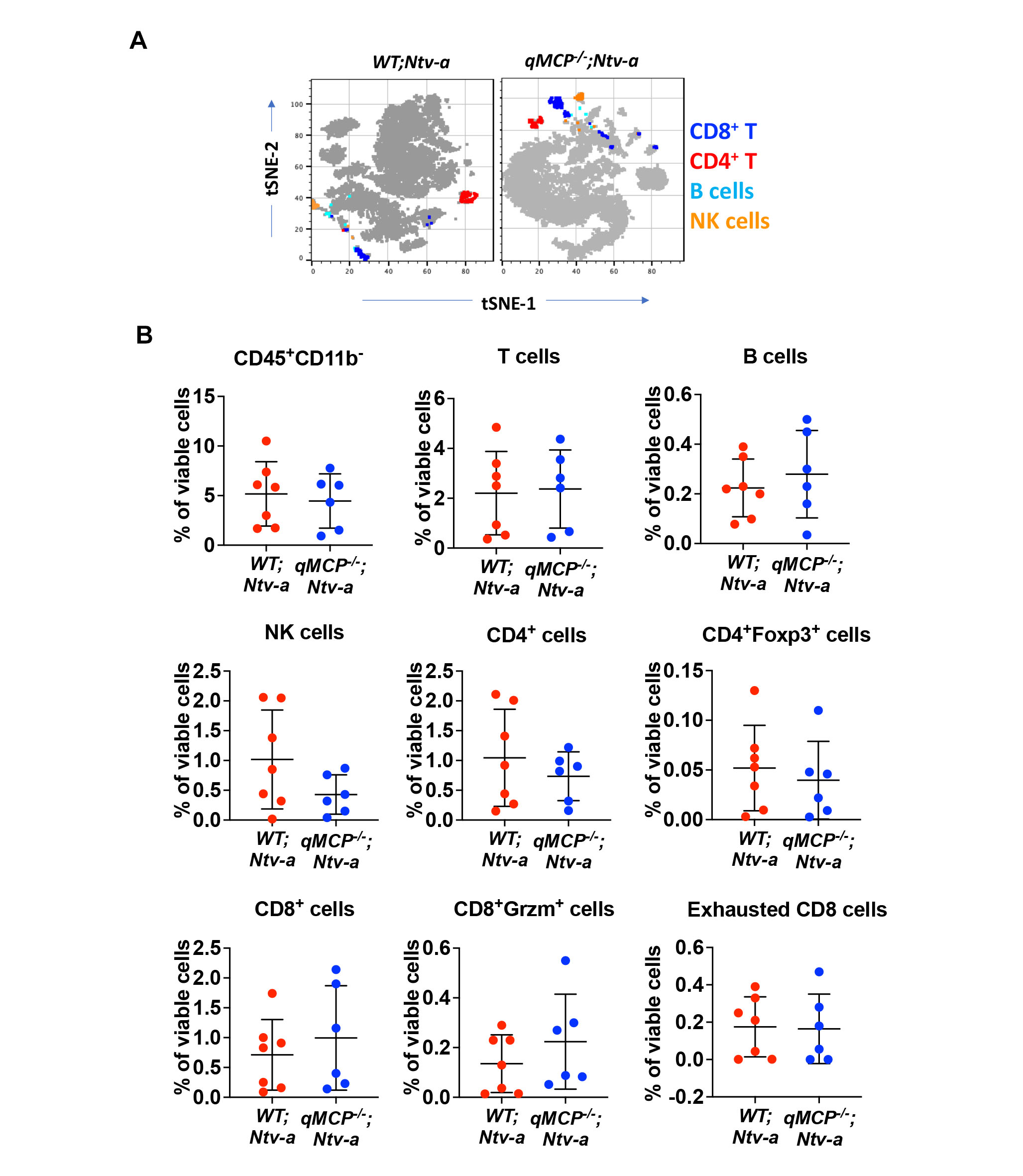

**Figure S9. Spectral flow cytometry analysis of lymphoid cells in GBM. (A)** tSNE plots showing lymphoid cells in GBM. (**B**) Dot plots quantifying subsets of lymphoid cells in the tumors. Student’s *t*-test was applied.

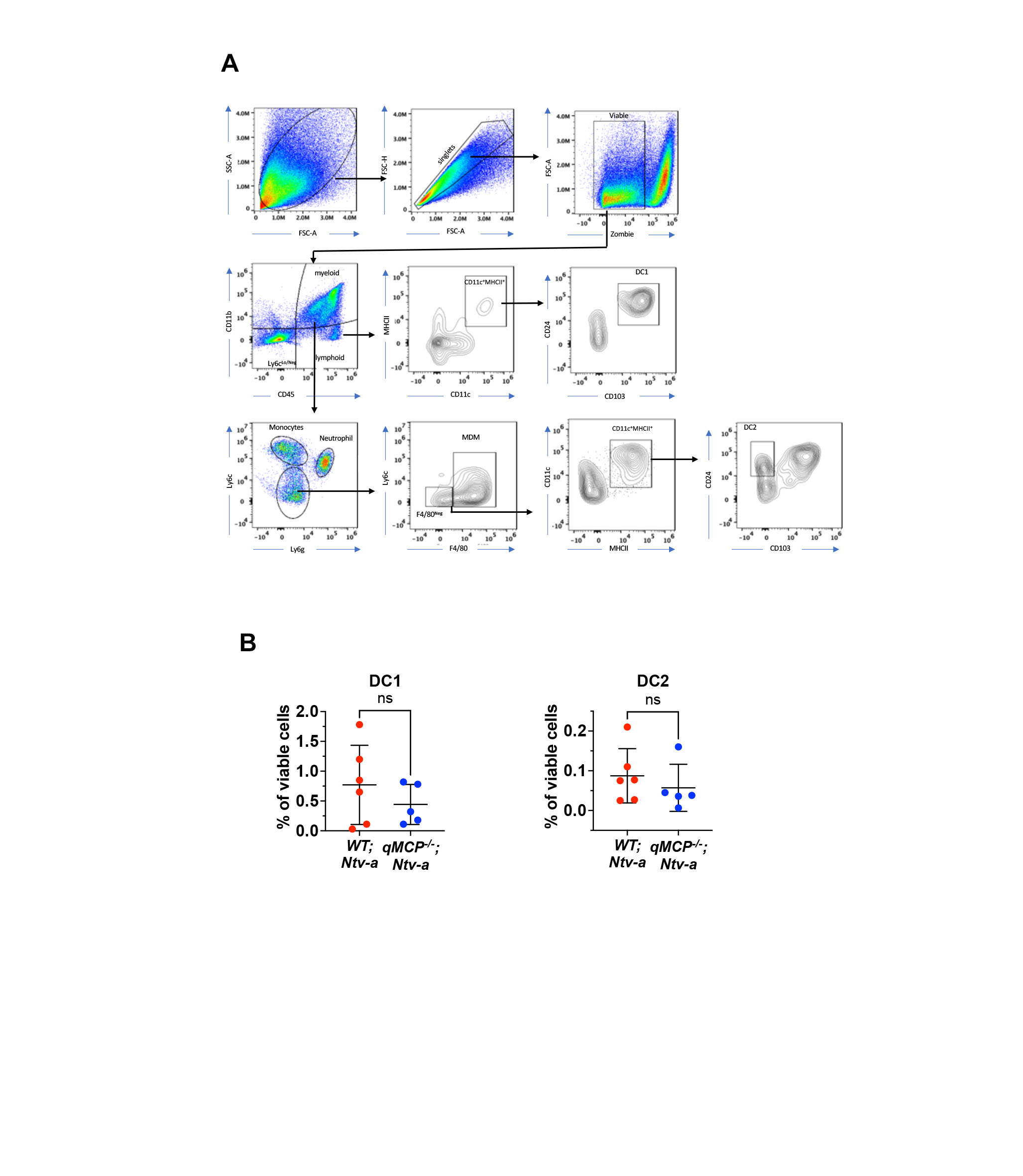

**Figure S10. Spectral flow cytometry analysis of dendritic cells (DC) in GBM. (A)** Gating strategy identifying DCs in GBM. (**B**) Dot plots quantifying DCs in the tumors. Student’s *t*-test was applied. Ns = not significant.

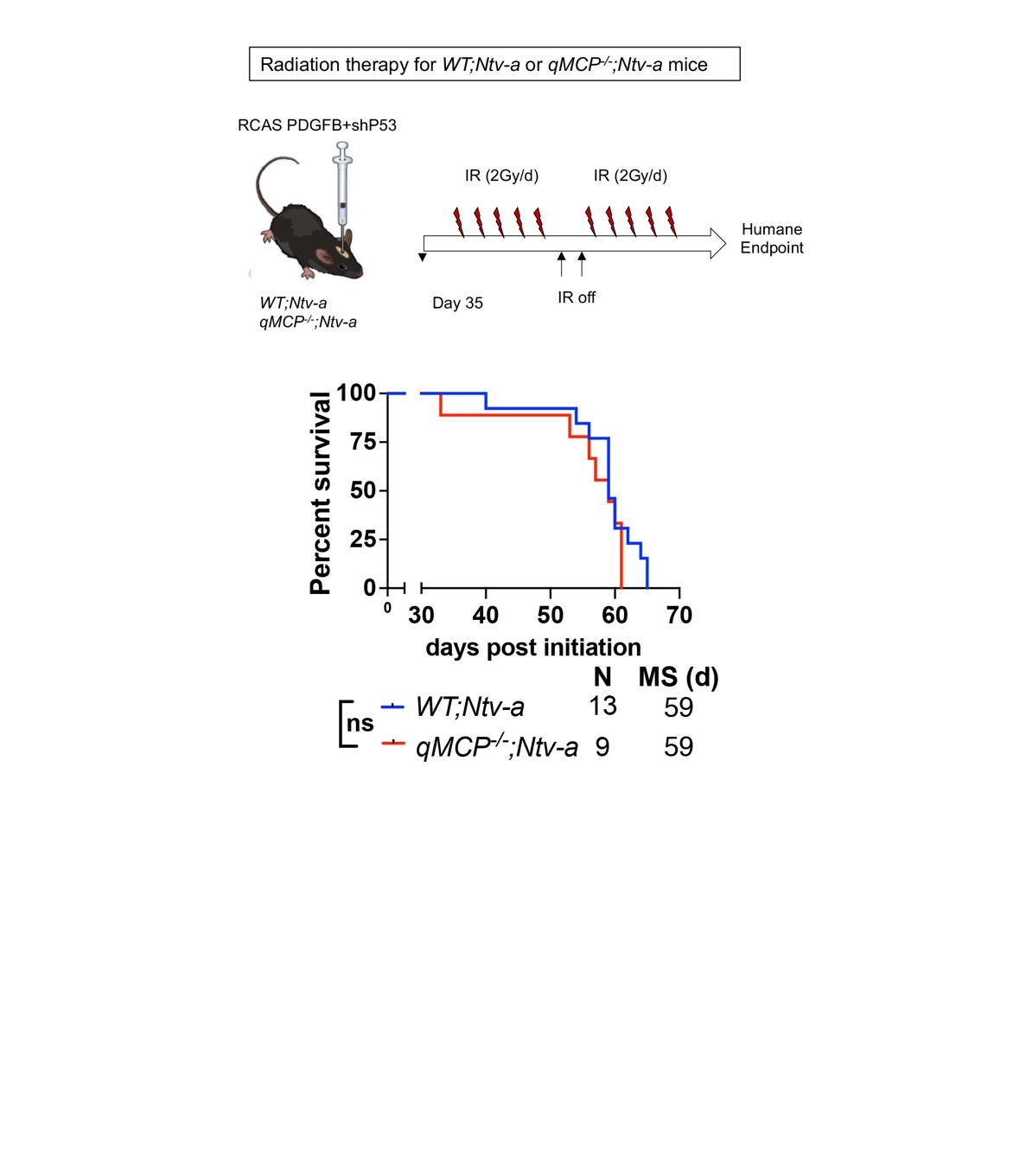

**Figure S11. Radiation therapy (RT) does not prolong the survival of *qMCP^-/-^;Ntv-a* mice*.*  (A)** Illustration of experimental design. IR = irradiation. (**B**) Kaplan-Meier survival curves of tumor-bearing *WT;Ntv-a* and *qMCP^-/-^;Ntv-a* mice.

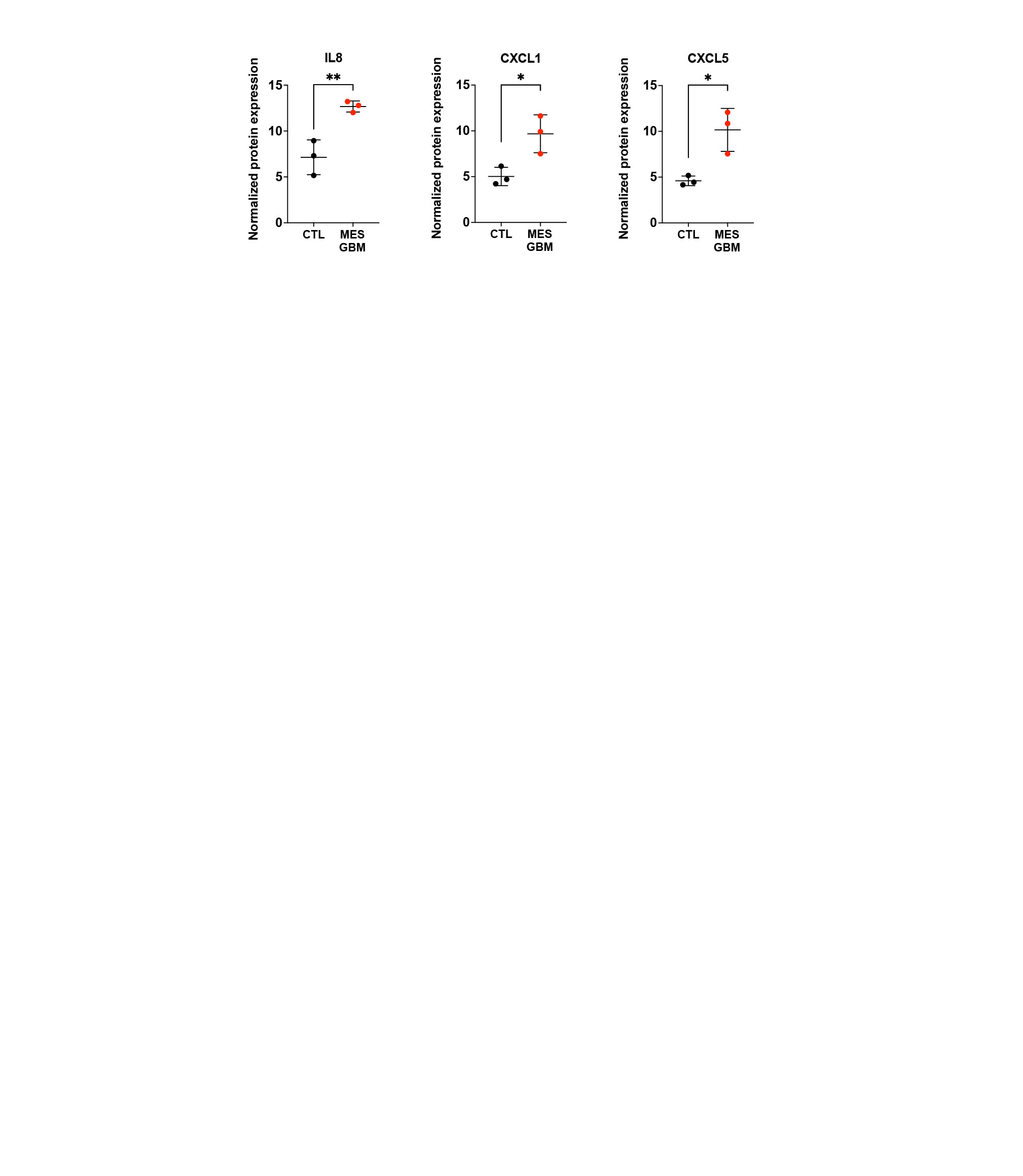

**Figure S12. Normalized protein expressions of CXCL chemokines examined by Olink proteomic assay between normal control and MES GBM.** Student's *t*-test. *p<0.05, **p<0.01.

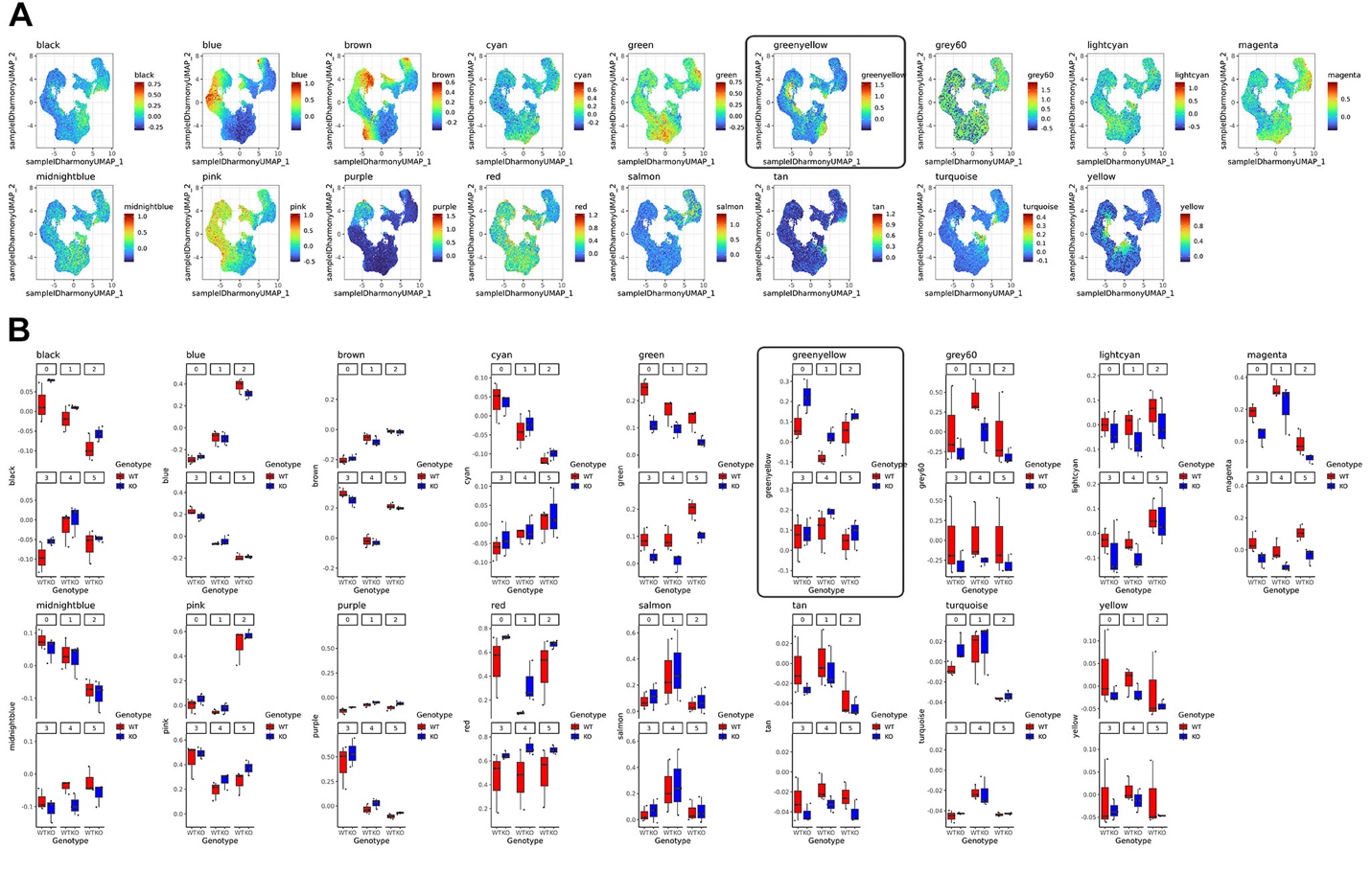

**Figure S13. Identification of cancer co-expression modules in scRNA-seq data using WGCNA. (A)** UMAPs of module scores for each cancer module identified by WGCNA. (**B**) Box plots of average module scores per sample split by genotypes (red vs. blue) and tumor clusters (0-5). “Greenyellow” module show pronounced difference between *WT;Ntv-a* (red) and *qMCP^-/-^;Ntv-a* (blue) mice, and was outlined.

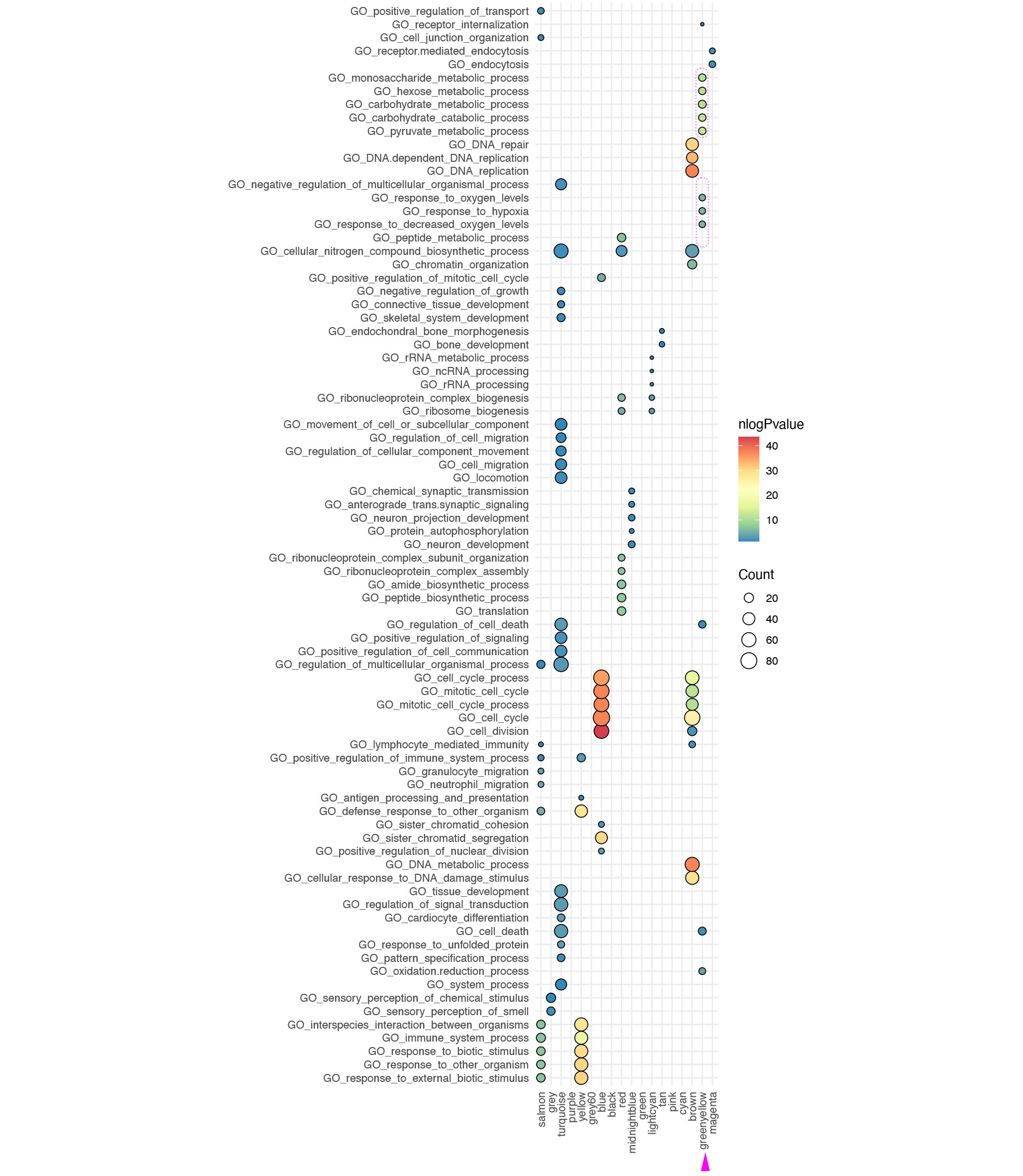

**Figure S14. Top enriched gene ontology (GO) terms for co-expressed genes identified in each WGCNA module.** Dot plot showing the top five GO biological process terms enriched for genes in each WGCNA module. “Greenyellow” module genes were enriched for hypoxic response and glycolysis GO terms (dotted lines). Size of dots represents the number of genes in the enrichment and color of dots present the -log(P-value) of the pathway enrichment.

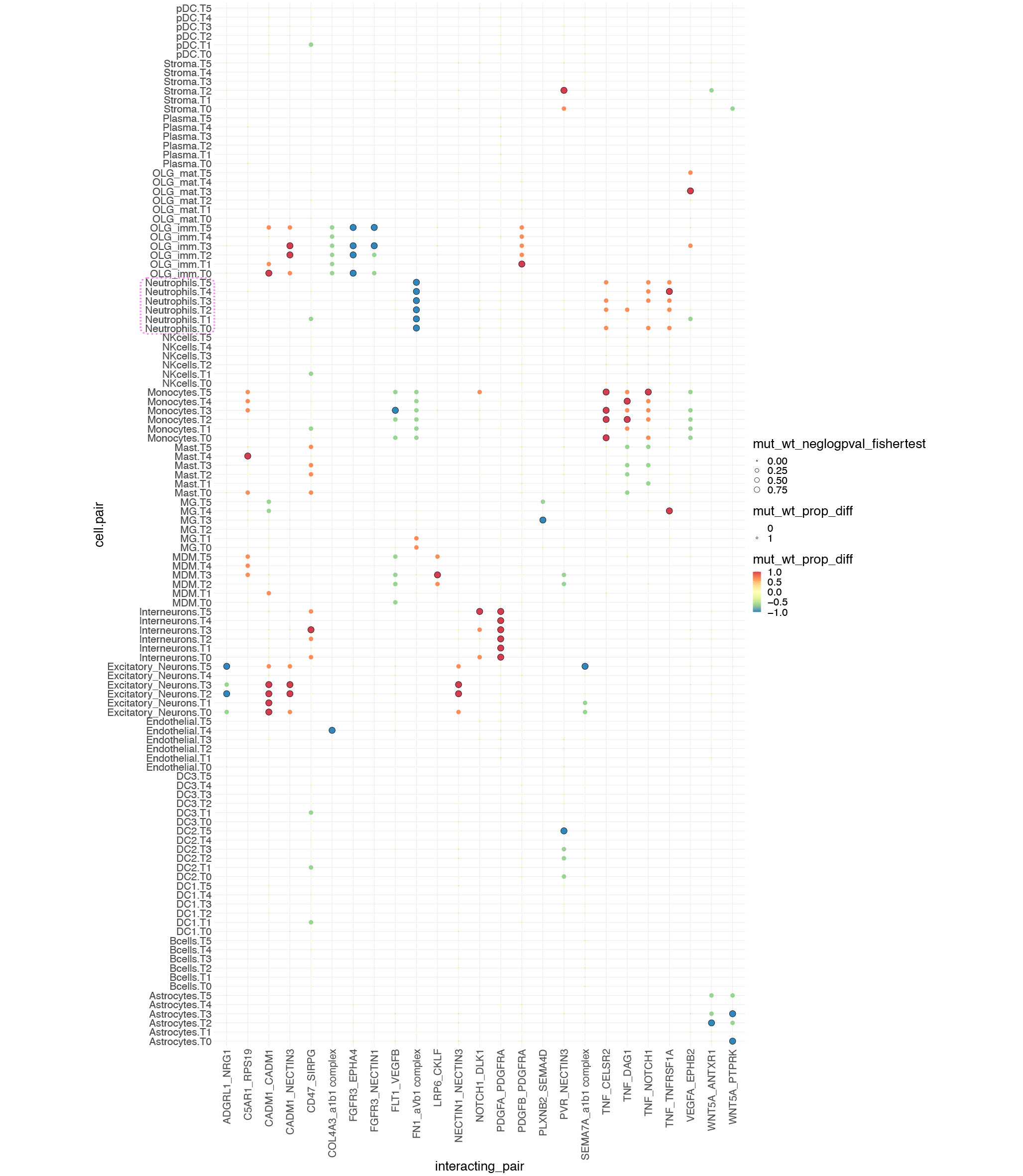

**Figure S15. Differential ligand-receptor analysis between KO and WT samples.** Dot plot showing the differential enrichment of interactions between corresponding cell-cell (y-axis) and ligand-receptor pairs (x-axis). Interactions were identified by CellPhoneDB. Color of dots represents the proportional difference in enrichment between *WT;Ntv-a* (blue) and *qMCP^-/-^;Ntv-a* (red) samples, size of dot represents the -log(P-value) of the differential enrichment (Fisher exact test), and dark circle outline represents a significant differential interaction (*P* < 0.05).

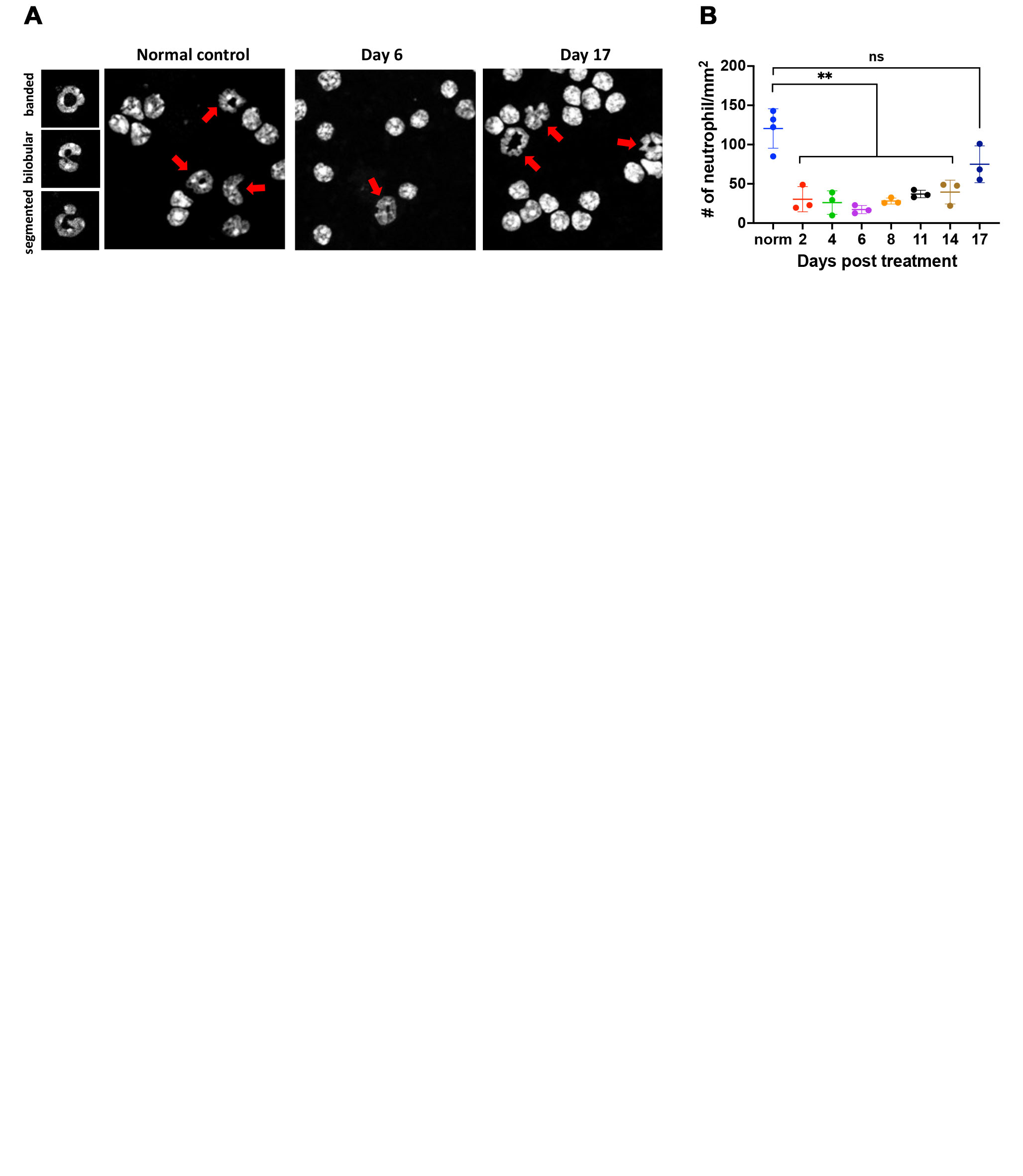

**Figure S16. Depletion of neutrophil by anti-Ly6g antibody is transient. (A)** Morphology of neutrophils analyzed by Cytospin and fluorescent microscopy. (B) quantification of blood neutrophils over time during anti-Ly6g antibody treatment in healthy adult mice.

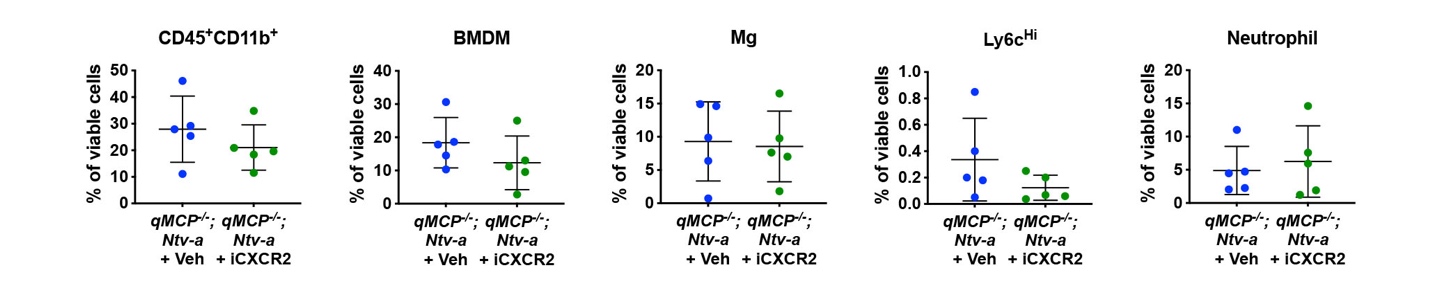

**Figure S17. FACS quantification of myeloid subtypes** **in HCC**. Student’s *t*-test was applied. BMDM = bone marrow derived myeloid cells, Mg = microglia. iCXCR2 = CXCR2 inhibitor.

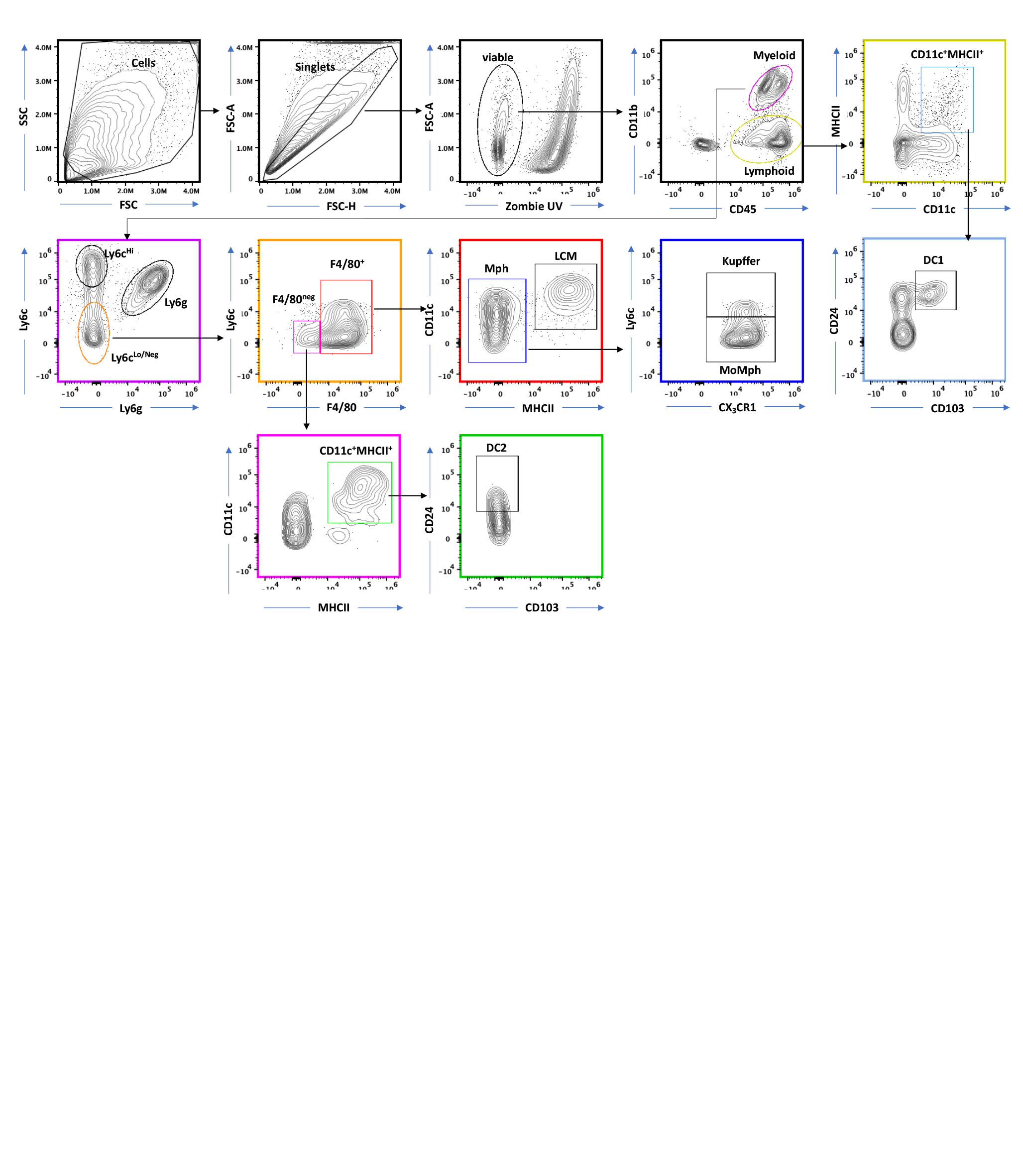

**Figure S18. Spectral flow cytometry analysis of myeloid cells in HCC.** Gating strategy is shown.

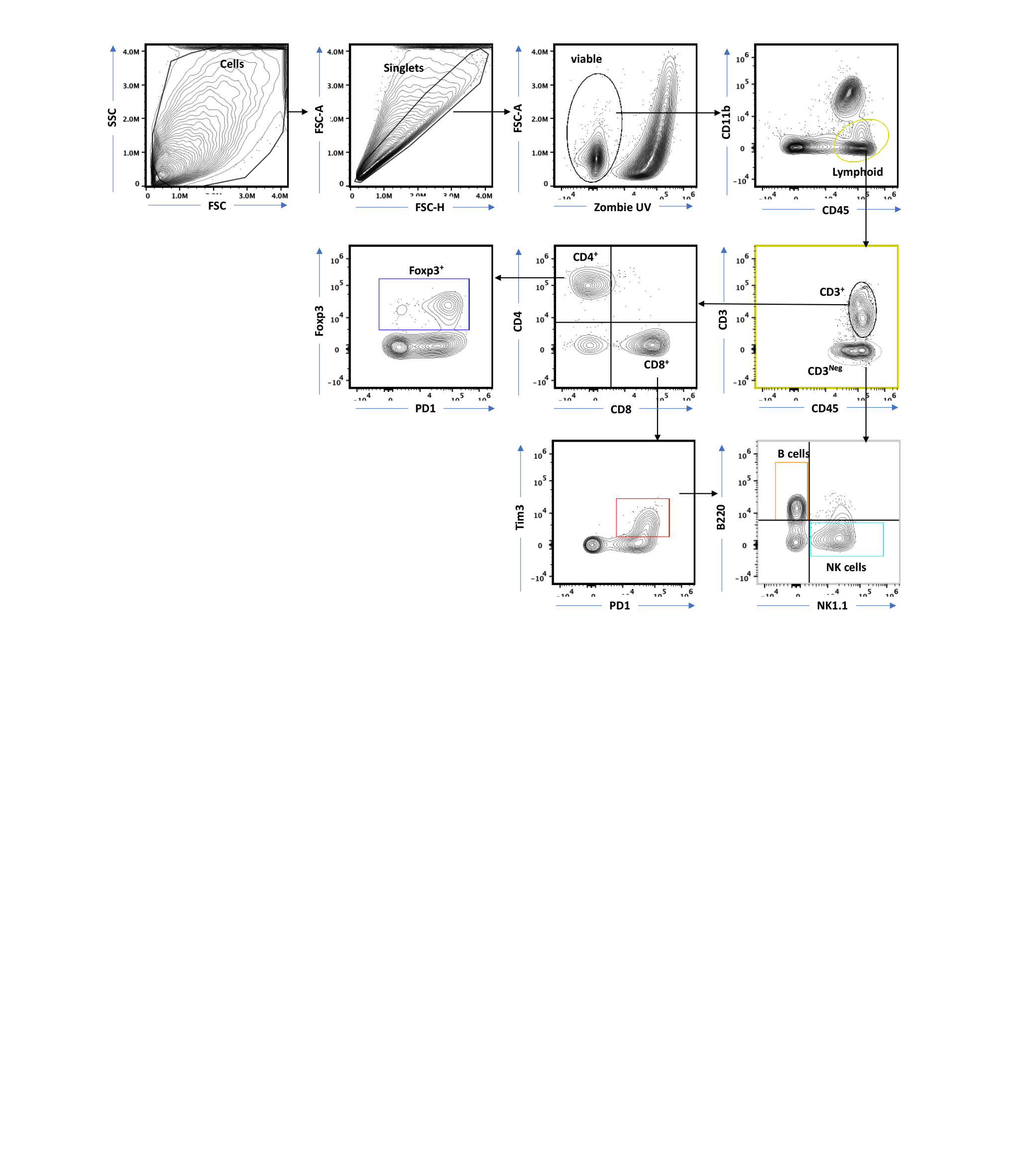

**Figure S19. Spectral flow cytometry analysis of lymphoid cells in HCC.** Gating strategy is shown.

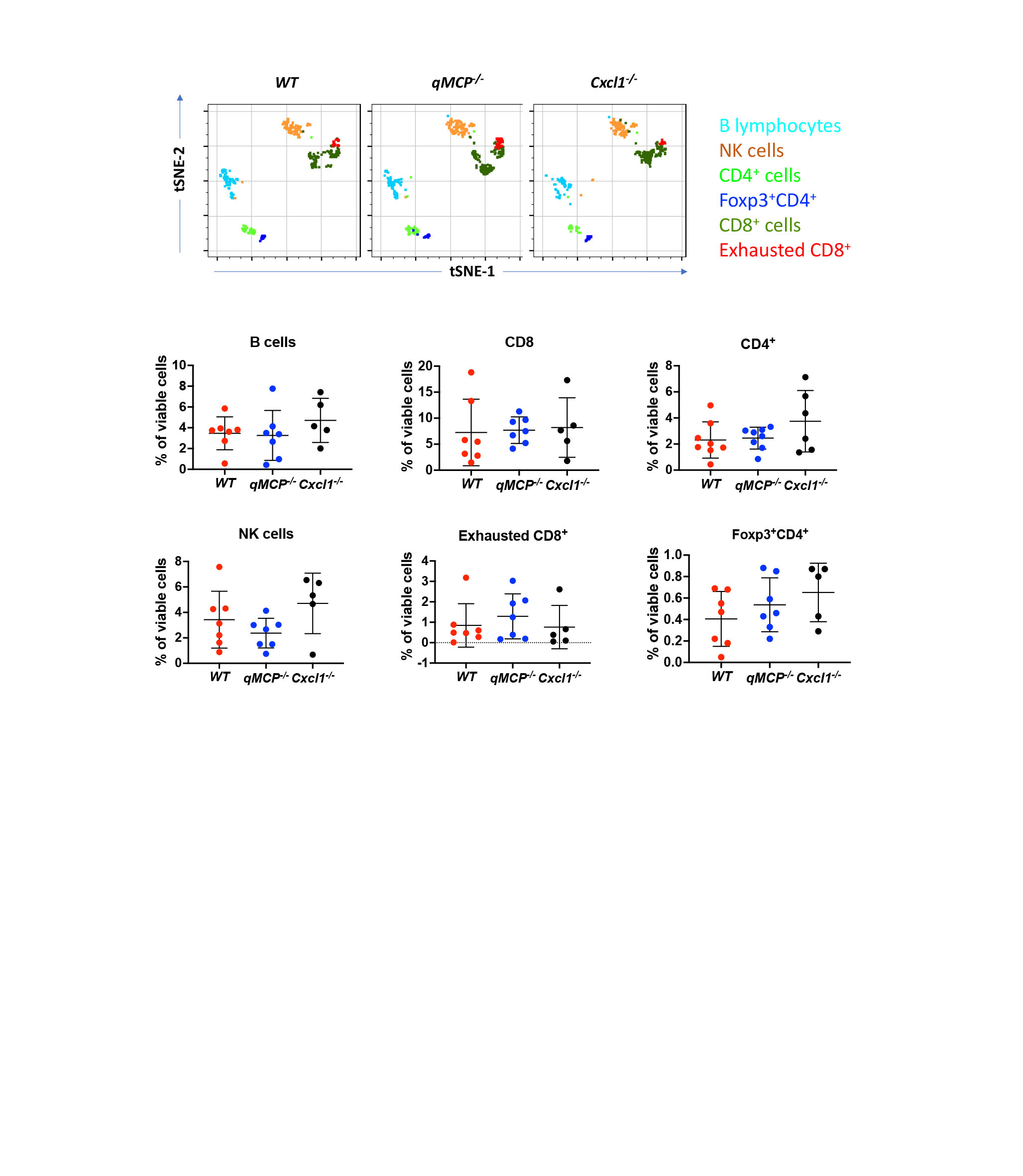

**Figure S20. Spectral flow cytometry analysis of lymphoid cells in HCC. (A)** tSNE plots illustrating lymphoid subsets of immune cells in HCC. (**B**) Quantification of lymphoid cells in HCC by spectral flow cytometry.

**Supplementary Tables**

**Table S1. Patient information.**

| **Sample #** | **Tumor diagnosis** | **Sex** | **Age** | **Sub-classification** | **Molecular driver** |
| --- | --- | --- | --- | --- | --- |
| 16302 (A2) | GBM, IDH-WT | Female | 74 | MES | NF1 mutation |
| 21810 | GBM, IDH-WT | Female | 66 | CL | EGFR amplified, TERT promoter c-146C>T, TP53 p.Y220D missense; PTEN loss; CDKN2A/B loss |
| 24286 | GBM, IDH-WT | Male | 46 | CL | EGFR amplified; EGFR G598V - subclonal, EGFRvIVa† (ex 24-27 del); ATM splice site 3576G>A; KDR R961W; CDKN2A/B loss; MTAP loss; TERT promoter -124C>T; VUS: CARD11 A687V; CEBPA V287G; CRKL T213A; GATA4 R318T; MAP3K1 T1511I; MTAP M169V; NTRK1 G18E; NTRK2 S167Y; PTEN F154I; SETD2 V932I; |
| 27419 | GBM, IDH-WT | Female | 56 | MES | NF1 splice site 2850+1G>A; CDKN2A loss; PTEN M199del; TERT -146C>T; TP53 T211_H214del; MS-Stable; 4 muts/ MB |
| 29282 | GBM, IDH-WT | Female | 58 | CL | EGFR amplified; EGFR p.A289V; PTEN p.K332Tfs*8; TERT c.-124C>T |
| 31100 | GBM, IDH-WT | Male | 75 | MES | CDKN2A loss; MTAP loss; NF1 N2387_F2388de; NF1 K1444E; PTEN loss; PTPN11 Y63C; TERT promoter -146C>T; MS-Stable; 1 mut/Mb; |
| 11849 | GBM, IDH-WT | Female | 74 | PN | PDGFRa and KIT amplifications |

**Table S2. qPCR Primers Used in Study.** The Bio-Rad qPCR primers used in the study are listed with their catalog numbers.

| **Primer** | **Bio-Rad Catalog Number** |
| --- | --- |
| Abcg2 | qMmuCID0009104 |
| Actb | qMmuCED0027505 |
| Aif1 | qMmuCED0046745 |
| Ascl1 | qMmuCED0044820 |
| Ccl11 | qMmuCED0044849 |
| Ccl12 | qMmuCED0061017 |
| Ccl2 | qMmuCED0048300 |
| Ccl7 | qMmuCED0049027 |
| Ccl8 | qMmuCED0003781 |
| Ccr1 | qMmuCID0006862 |
| Cd44 | qMmuCID0025677 |
| Cebpb | qMmuCED0050360 |
| Chi3l1 | qMmuCID0015758 |
| Cxcl3 | qMmuCED0001059 |
| Cxcl5 | qMmuCED0003886 |
| Dll3 | qMmuCID0023659 |
| Gfap | qMmuCID0020163 |
| Hprt | qMmuCED0045738 |
| Met | qMmuCID0017026 |
| Mgmt | qMmuCID0009593 |
| Olig2 | qMmuCED0003760 |
| Rps6ka3 | qMmuCID0006067 |
| Serpine1 | qMmuCID0027303 |
| Sox2 | qMmuCED0051857 |
| Stat3 | qMmuCED0044698 |
| Taz | qMmuCID0020469 |
| Tgfb1 | qMmuCID0017320 |
| Yap1 | qMmuCID0005990 |

**Table S3. Primary antibodies used in the study.**

| **Antibody** | **Application** | **Specificity** | **Manufacturer** | **Catalog Number** |
| --- | --- | --- | --- | --- |
| IBA1 | IHC/IF | Human, Mouse | Wako | 019-19741 |
| Anti-Ly6g | In vivo injection | Mouse | BioXcell | BE0075-1 |
| Anti-trinitrophenol | In vivo injection | Mouse | BioXcell | BE0089 |
| OLIG2 | IHC | Mouse | Millipore Sigma | AB9610 |
| CD31 | IHC | Mouse | Dianova | DIA-310 |
| CD44 | IHC | Human, Mouse | BD Pharmingen | 550538 |
| GFAP | IHC | Mouse | CST | 3670 |
| Elane | IHC | Human, Mouse | Bioss | bs6982R |
| Elane | IHC | Human, Mouse | AbCam | ab68672 |
| P_2_Y_12_ | IHC | Mouse | AnaSpect | SQ-ANAB-78839 |
| CD45-APC | Flow Cytometry | Mouse | BioLegend | 103112 |
| CD45-PE | Flow Cytometry | Mouse | BioLegend | 103106 |
| CD45-FITC | Flow Cytometry | Mouse | BioLegend | 103107 |
| CD45-V450 | Flow Cytometry | Mouse | BD Biosciences | 560501 |
| CD45-BV510 | Flow Cytometry | Mouse | BD Biosciences | 563891 |
| B220-BV605 | Flow Cytometry | Mouse | BioLegend | 103243 |
| B220-AF700 | Flow Cytometry | Mouse | BioLegend | 103232 |
| CD101-APC | Flow Cytometry | Mouse | Invitrogen | 17101180 |
| CD103-BUV395 | Flow Cytometry | Mouse | BD Biosciences | 748253 |
| CD11b-APC-Cy7 | Flow Cytometry | Mouse | BioLegend | 101226 |
| CD11b-PerCP-Cy5.5 | Flow Cytometry | Mouse | BD Biosciences | 550993 |
| CD11c-PE-Dazzle594 | Flow Cytometry | Mouse | BioLegend | 117348 |
| CD11c-APC | Flow Cytometry | Mouse | BD Biosciences | 550281 |
| CD19-BV785 | Flow Cytometry | Mouse | BD Biosciences | 563333 |
| CD24-BUV496 | Flow Cytometry | Mouse | BD Biosciences | 612953 |
| CD3-PE-dazzle | Flow Cytometry | Mouse | BioLegend | 100348 |
| CD4-APC-Cy7 | Flow Cytometry | Mouse | BioLegend | 100526 |
| CD49d-PE-dazzle | Flow Cytometry | Mouse | BioLegend | 103625 |
| CD5-PE-Cy5 | Flow Cytometry | Mouse | BioLegend | 100610 |
| CD8-BV510 | Flow Cytometry | Mouse | BioLegend | 100752 |
| CX3CR1-PerCP-Cy5.5 | Flow Cytometry | Mouse | BioLegend | 149009 |
| CX3CR1-BV650 | Flow Cytometry | Mouse | BioLegend | 149033 |
| CXCR2-PE | Flow Cytometry | Mouse | BioLegend | 149609 |
| F4/80-PE-Cy7 | Flow Cytometry | Mouse | BioLegend | 123114 |
| F4/80-BV711 | Flow Cytometry | Mouse | BioLegend | 123147 |
| Foxp3-FITC | Flow Cytometry | Mouse | Invitrogen | 11-5773-82 |
| Gr-1-BV711 | Flow Cytometry | Mouse | BioLegend | 108443 |
| GrzmB-PE | Flow Cytometry | Human, Mouse | Invitrogen | 12-8899-41 |
| IA/IE-BV650 | Flow Cytometry | Mouse | BD Biosciences | 563415 |
| IA/IE-Alex700 | Flow Cytometry | Mouse | BioLegend | 107622 |
| Ly6c-AF488 | Flow Cytometry | Mouse | BioLegend | 128022 |
| Ly6c-PE-Cy7 | Flow Cytometry | Mouse | BD Biosciences | 560593 |
| Ly6g-V450 | Flow Cytometry | Mouse | BD Biosciences | 560603 |
| NK1.1-AF647 | Flow Cytometry | Mouse | BioLegend | 108720 |
| NK1.1-BV711 | Flow Cytometry | Mouse | BD Biosciences | 740663 |
| PD-L1-BV605 | Flow Cytometry | Mouse | BD Biosciences | 745135 |
| PD1-BV785 | Flow Cytometry | Mouse | BioLegend | 135225 |
| Tim-3-PE-Cy7 | Flow Cytometry | Mouse | Invitrogen | 25-5870-82 |
